# Short term tomato consumption alters the pig gut microbiome towards a more favorable profile

**DOI:** 10.1101/2022.05.13.489542

**Authors:** Mallory L. Goggans, Emma A. Bilbrey, Cristian Quiroz-Moreno, David M. Francis, Sheila K. Jacobi, Jasna Kovac, Jessica L. Cooperstone

## Abstract

Diets rich in fruits and vegetables have been shown to exert positive effects on the gut microbiome. However, little is known about the specific effect of individual fruits or vegetables on gut microbe profiles. This study aims to elucidate the effects of tomato consumption on the gut microbiome, as tomatoes account for 22% of vegetable consumption in Western diets, and their consumption has been associated with positive health outcomes. Using piglets as a physiologically relevant model of human metabolism, 20 animals were assigned either to a control or tomato powder supplemented diet (both macronutrient matched and isocaloric) for 14 days. The microbiome was sampled rectally at three time points: day 0 (baseline), day 7 (midpoint), and at day 14 (end of study). DNA was sequenced using shotgun metagenomics, and reads were annotated using MG-RAST. There were no differences in body weight or feed intake between our two treatment groups. There was a microbial shift which included a higher ratio of Bacteroidota to Bacillota (formerly known as Bacteroidetes and Firmicutes, respectively) and higher alpha-diversity in tomato-fed animals, indicating a shift to a more desirable phenotype. Analyses at both the phyla and genera levels showed global microbiome profile changes (PERMANOVA P ≤ 0.05) over time, but not with tomato consumption. These data suggest that short-term tomato consumption can beneficially influence the gut microbial profile, warranting further investigation in humans.

**IMPORTANCE:** The composition of the microorganisms in the gut is a contributor to overall health, prompting the development of strategies to alter the microbiome composition. Studies have investigated the role of the diet on the microbiome, as it is a major modifiable risk factor contributing to health; however, little is known about the causal effects of consumption of specific foods on the gut microbiota. A more complete understanding of how individual foods impact the microbiome will enable more evidence-based dietary recommendations for long-term health. Tomatoes are of interest as the most consumed non-starchy vegetable and a common source of nutrients and phytochemicals across the world. This study aimed to elucidate the effect of short-term tomato consumption on the microbiome, using piglets as a physiologically relevant model to humans. We found that tomato consumption can positively affect the gut microbial profile, which warrants further investigation in humans.

## INTRODUCTION

Research has shown that the composition of the gut microbiome can be an effector of overall health (1). The composition of these gut microorganisms has been associated with a number of chronic diseases, such as cardiovascular disease (2), inflammation (3), type 2 diabetes (1), and obesity (3–5). As diet is a major modifiable factor of health, there is interest in elucidating how dietary factors can alter the microbiome (6, 7). While it is possible to use some microbiome endpoints and associate them with health (i.e., a more diverse community is favorable (1, 6, 8), and a lower Bacteroidota to Bacillota (formerly known as Bacteroidetes and Firmicutes respectively) ratio (4), the reality is that bias in sequencing approaches as well as differences in microbial communities due to lifestyle factors and location add complexity to this interpretation (9). Still, diets rich in fruits, vegetables, and whole grains have been consistently associated with a healthier microbiome (6–8, 10). However, discerning the way specific foods might affect the microbiome using intervention studies remains largely uninvestigated. Understanding the global effects that specific foods have on the microbiome helps contextualize the effect they are having towards overall health and sets a foundation towards making personalized nutritional recommendations.

Tomatoes are of interest as one such specific food because they are a common source of nutrients for many around the world. They are the second most commonly consumed vegetable (11) and are an important specialty crop across the United States. Over 12 million metric tons of tomatoes are produced in the United States each year (12), with Americans consuming about 30 pounds per person in 2018 (13). Tomatoes are a rich source of both essential nutrients (e.g., vitamins A, C), fiber, and phytochemicals (e.g., lycopene, flavonoids, phenolic acids). Tomato consumption has been linked to protection against various chronic diseases (14–16), though causality about the mechanism of action is not well understood.

We hypothesized that one mechanism by which tomatoes provide a health benefit is through their modulation of the gut microbiome. Preliminary microbiome studies in mice, feeding tomatoes or their phytochemicals, have shown positive outcomes, including increased microbial diversity, decreased abundance of *Clostridium* spp., and decreased symptoms of irritable bowel disease (17–21). Here, we aimed to elucidate the effects of short-term, consistent tomato consumption on the gut microbial ecosystem, using pigs as a physiologically relevant model for humans. To investigate this question, we fed weaned piglets (n = 20, aged 4 weeks) a diet supplemented with 10% *w*/*w* tomato powder or an iso-caloric and macronutrient-matched control diet for two weeks, sampling the gut microbiome via rectal swab at three points during the experimental period. The use of macronutrient matched diets allowed us to test the effect of tomato phytochemicals on the microbiome of studied pigs, rather than the effect of differences in nutrients, such as fiber or sugar. DNA from rectal swabs was subjected to shotgun metagenomic sequencing (i.e., the untargeted sequencing of all the DNA present in a sample (22)). The resulting reads were annotated and analyzed at both the phyla and genera levels using univariate and multivariate approaches, including the analysis of beta diversity, relative abundances of Bacteroidota, Bacilotta, their ratio, and alpha diversity.

## Results AND DISCUSSION

### Diet type did not affect animal weight

An overall scheme of the animal study design can be found in **FIG 1**. Pigs were weighed and feed intake was measured weekly. There was no difference in feed intake or animal weight over the trial (**Table S1**). Health of pigs was not altered by dietary treatment.

**FIG 1.**
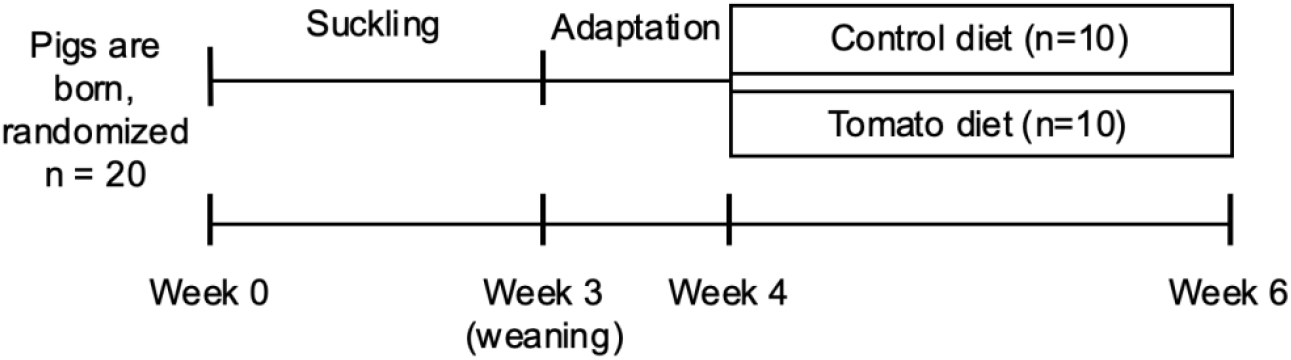
Overall animal study design. Pigs were adapted to a dry diet from weeks 3-4. Microbiome was sampled via rectal swabs when pigs were aged 4 weeks (day 0, baseline), 5 weeks (day 7, midpoint) and 6 weeks (day 14, end of study) for shotgun metagenomics.

### A median sequence depth was 2.5M reads

Each sample’s forward and reverse reads were checked for quality using FastQC version 0.11.9 (23). All sequence files passed quality checks and no samples had to be discarded. Thirteen of the 60 total samples were re-sequenced to a median of 2.3M reads per sample. For re-sequenced samples, sequences from the first and second sequencing run were merged, checked for quality and were used for further analyses. Rarefaction curves demonstrate that a similar species richness was achieved in samples with differing sequence depths (**Fig. S1**). A recent study has shown that even shallow shotgun metagenomics (<500K reads/sample) provides better annotation of taxonomic and functional composition of microbiome compared to 16S rRNA sequencing (24), providing our rationale for this sequencing approach.

### Bacillota (i.e., Firmicutes) was the predominant phylum and *Prevotella* the most abundant genus detected in the pig fecal microbiome

The mammalian gut microbiome is a complex microbial ecosystem; hence, it is beneficial to conduct analyses at more than one taxonomic rank, as the profile of each rank provides different types of information. The average human gut microbiome is dominated by Bacteroidota (formerly known as Bacteriodetes) and Bacillota (formerly known as Firmicutes), which typically account for 70-90 % of the total microbiome makeup (1). Analyses of phyla often reveal changes in the proportions of the dominant few, thus providing a broad picture of the state of the gut microbiome. Alternatively, genera are highly diverse, often with hundreds of taxa identified (25). These analyses provide a finer resolution of microbiome composition. Here, we aimed to capture modifications of the microbiome at both the phyla and genera level. For this reason, all analyses (aside from those specific to phyla) were completed at both taxonomic ranks. Across all pigs, annotation using MG-RAST and filtering for data quality resulted in identification of 45 phyla. Of those, 28 were from the domain Bacteria, comprising on average 99.3 ± 0.2% of the total reads, 10 were from Eukaryota, 5 were Archaea, 1 was Virus, and 1 was unclassified. The most prevalent phyla were Bacilotta (formerly known as Firmicutes 52.7% average abundance ± 5.5% standard deviation), Bacteroidota (formerly known as Bacteroidetes 35.4 ± 5.9), Actinomycetota (formerly known as Actinobacteria) (4.7 ± 1.8%), Pseudomonadota (formerly known as Proteobacteria) (3.9 ± 1.2%) and Fusobacteriota (formerly known as Fusobacteria) (0.43 ± 8.5×10^−4^%). Similar relative abundances of phyla were observed across samples, regardless of the diet groups. Previous studies reported conflicting results in terms of predominant phyla in pig microbiome. Some studies have shown Firmicutes to be the most abundant phyla in the pig gut microbiome after weaning (26, 27), while others have reported Bacteroidetes as the dominant phyla (28).

Annotation from MG-RAST and filtering for data quality resulted in the identification of 755 genera. Of these 755 genera, 582 were in the Bacteria domain, 89 were Eukaryota, 60 were Archaea, 23 were Viruses, and 1 was unclassified. Overall, the most prevalent genera were *Prevotella* (22.23% average abundance ± 5.4% standard deviation), *Bacteroides* (10.34 ± 1.9%), *Clostridium* (8.56 ± 1.8%), *Lactobacillus* (6.78 ± 4.6%) and *Eubacterium* (5.16 ± 1.0%). These genera were detected in similar relative abundances in each group when data were parsed by diet. Previous reports have shown *Prevotella, Bacteroides*, and *Clostridium* to be the most abundant genera in pig gut microbiomes (27), which is consistent with our findings.

### Beta diversity changed over time, but was not significantly affected by the tomato-supplemented diet

To understand the beta-diversity (differences between the microbial communities) of pigs on different diets and at different time points, all data was first visualized via principal coordinates analysis (PCoA) using the Bray-Curtis dissimilarity metric. PCoA plots (**Fig. 2**) were created for both phyla and genera separately using the relative abundances of all samples. Plots were faceted by diet to observe sample clustering by time point more easily. PC1 and PC2 together accounted for 89.1% of the variation in the phyla-level microbiome and 53.8% at the genera level. Visual clustering in PCoA scores plots at either taxonomic level was not easily observed between diets, but within the control diet, grouping was observed according to time point. It is not surprising that overall microbiome profile differences are not evident in the PCoA plots due to presence or absence of a single component of a diet (i.e., tomatoes). Global differences in microbiome composition are more likely to be observed when two completely different diets are fed, as previously shown when comparing the effect of a plant-based and animal-based diet on the microbiome (29).

**FIG 2.**
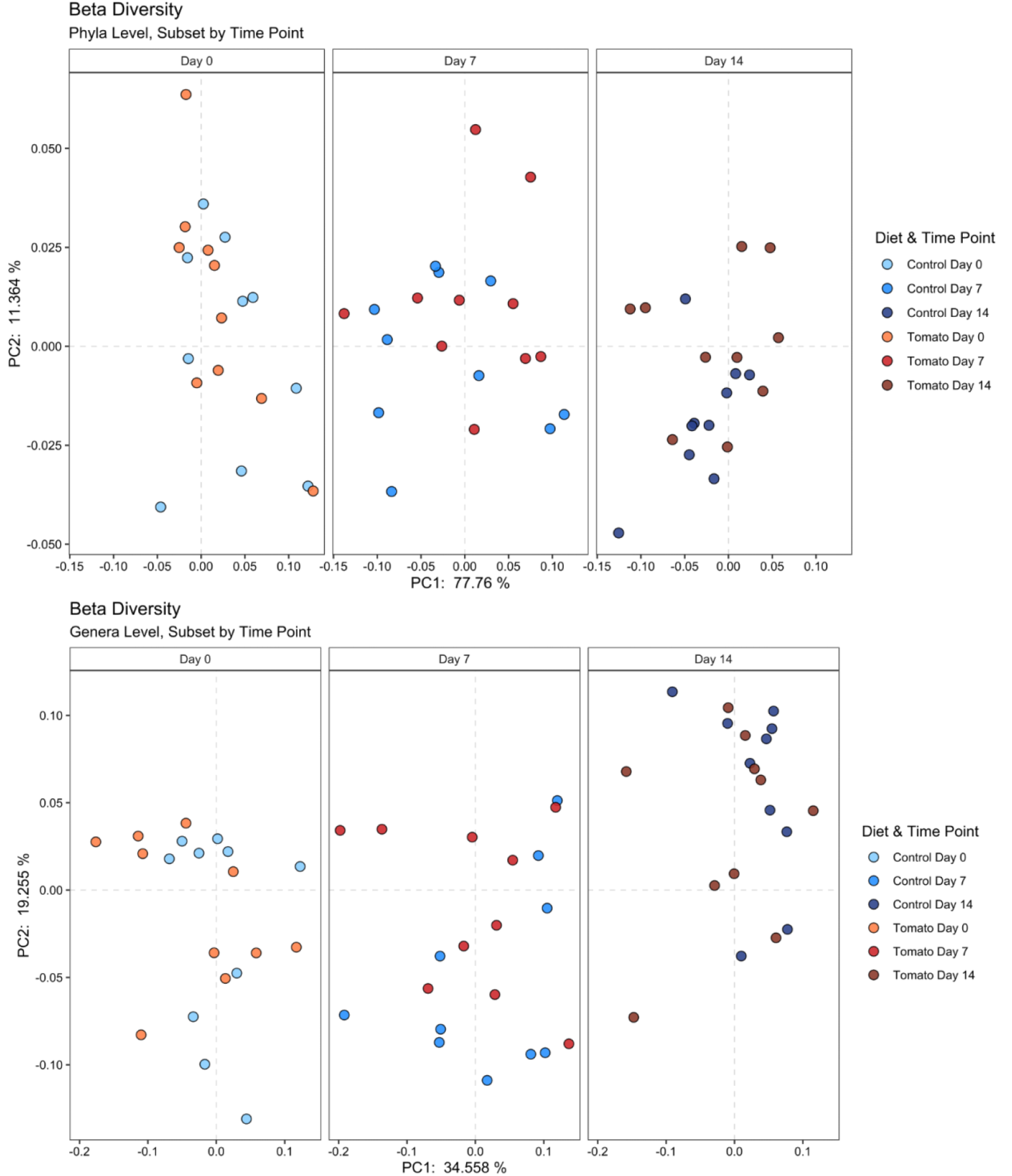
Principal coordinates analysis (PCoA) using Bray-Curtis distances showing beta-diversity of the whole microbiome at the phyla (top) level and genera (bottom) level. Each dot represents a sample collected from one pig. Plots are faceted by diet. Using repeated measures PERMANOVA (model: Beta Diversity = Diet + Time Point + Diet × Time Point + Error), only a significant effect of time point was detected at both the phyla (P = 0.020) and genera (P = 0.005) levels.

In order to determine significance of observed trends in the PCoA due to diet and time point, PERMANOVA was used (model: Beta Diversity = Diet + Time Point + Diet × Time Point

+ Error, where each pig was a plot containing three samples collected over time). These multivariate restricted permutation tests are a useful approach for assessing differences in beta-diversity because they allow for the investigation of the gut microbiome as a whole, instead of focusing on individual taxa. The PERMANOVA model p-values are recorded in **Table 1**. At both the phyla and genera levels, we found the interaction term to be non-significant (Phyla P = 0.510; Genera P = 0.360) and therefore we removed it from the model. The new model was then tested and revealed an overall significant effect of time point (Phyla P = 0.020; Genera P = 0.005) but not diet (P = 0.270) on the gut microbiome (**Table 1**).

**TABLE 1.** Results from restricted permutation tests via PERMANOVA to investigate differences in beta-diversity at the phyla and genera taxonomic levels. The full model tested the variance explained by the diet, time point, and their interaction on the dissimilarity matrix, calculated with Bray-Curtis distances.

| Variable | Phyla | Genera |
| --- | --- | --- |
| Diet | 0.270 | 0.060 |
| Time Point | 0.020<sup>a</sup> | 0.005<sup>b</sup> |
| Diet × Time Point | 0.510 | 0.360 |

<sup>a</sup> Indicates significant model effect at $P \leq 0.05$ .
<sup>b</sup> Indicates significant model effect at $P \leq 0.01$ .

These data can be interpreted in that, at both taxonomic ranks, the microbiomes of pigs were significantly changing over the two-week intervention, but the effect of diet on beta diversitywas not significant. In another study using a mouse model, the microbiomes were compared between a group fed a high fat diet supplemented with tomato powder and a high fat-only diet group. Using clustering by unweighted UniFrac dissimilarity, a significant difference was detected between diet group microbiomes (17). However, using weighted UniFrac distances, no separation of tomato and control groups was observed in the pigs. It is possible that using a dissimilarity measure that incorporates evolutionary relatedness may have been a contributor to the detected significant effects. However, a direct comparison with our study is difficult because mice are known to be different than pigs in their microbiome composition (30).

### Inverse relationship between Bacteroidota and Bacillota abundances was detected over time in the control-fed pigs, but not tomato-fed pigs

In addition to multivariate approaches to understand microbiome data, univariate methods to examine differences in specific taxa are valuable. As previously stated, the phyla Bacteroidota (i.e., Bacteriodetes) and Bacillota (i.e., Firmicutes) and their relationship have been implicated in obesity and high fat diets (31, 32). With these *a priori* interests, changes in these two phyla were assessed individually across diets and time points using repeated measures ANOVA. Results indicated a significant model effect of time point for both phyla (Bacteroidota P = 0.024; Bacillota P = 0.001); whereas diet and the interaction term were non-significant. After *post hoc* analyses to determine which pair-wise groups differed, significant alteration in the abundance of both Bacteroidota and Bacillota was found between day 0 and day 14 in control-fed pigs (Bacteroidota P = 0.044; Bacillota, P = 0.03). No significant differences between time points within the tomato-fed pigs were observed. Box plots of the two phyla demonstrate the inverse relationship between Bacteroidota and Bacillota abundances over time in the control-fed pigs (**Fig. 3a)**.

**FIG 3.**
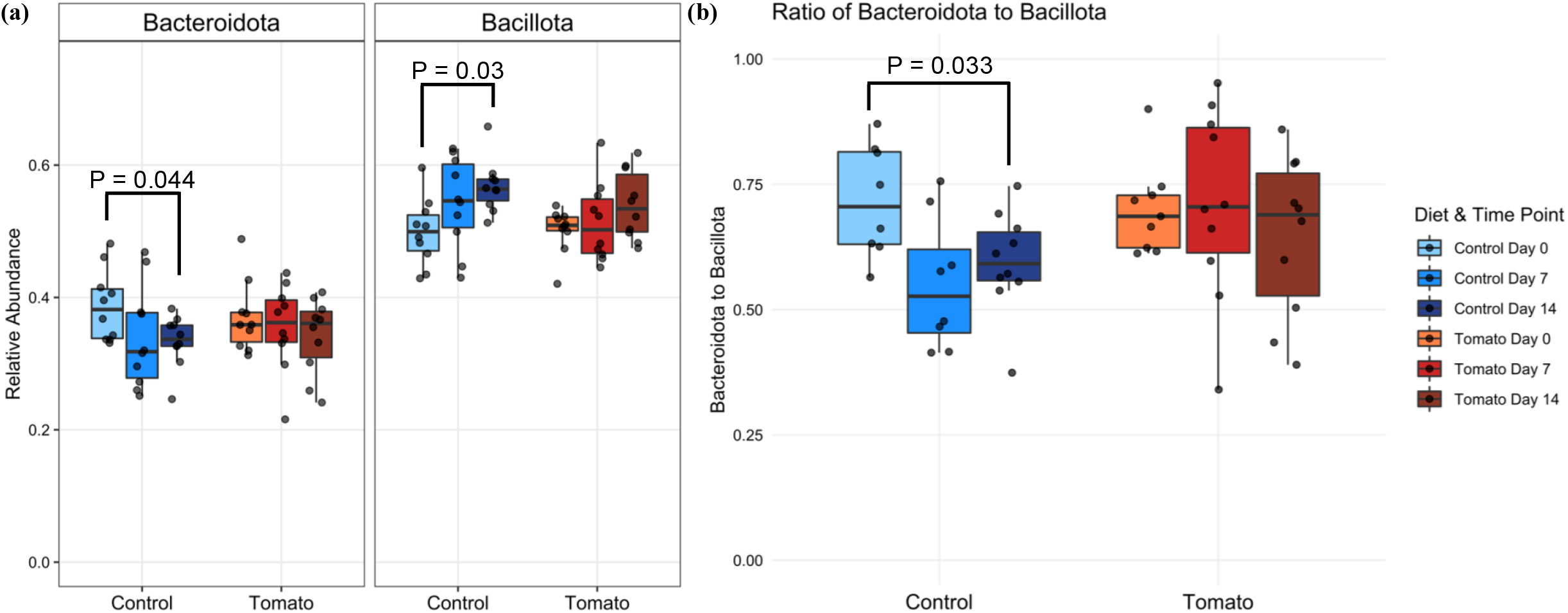
(a) Comparing relative abundances across time points and between diets for two phyla: Bacteroidota and Bacillota. Using repeated measures ANOVA, a significant model effect of time point was found for both phyla (Bacteroidota: P = 0.024, Bacillota: P = 0.001). *Post hoc* findings of significant differences between Day 0 Control and Day 14 Control were found for both Bacteroidota (P = 0.044) and Bacillota (P = 0.03). No significant effects of diet or time point-by-diet interaction were detected for either phylum. (b) Comparing the ratio of the relative abundance of Bacteroidota to that of Bacillota across time points and between diets. A significant effect of time via repeated measures ANOVA (P = 0.009) led to *post hoc* comparisons and a significant difference in the ratio of Bacteroidota/Bacillota between Day 0 and Day 14 in control-fed pigs only (P = 0.033). There was no significant effect of diet or time point-by-diet interaction.

Additionally, the ratio of Bacteroidota to Bacillota in the gut microbiome is a commonly assessed metric because of its correlation to obesity (4, 31, 32). Therefore, differences in Bacteroidota/Bacillota were also tested via repeated measures ANOVA with diet, time point, and their interaction as factors. This analysis revealed a significant difference due to time (P = 0.009), with a non-significant effect of diet (P = 0.728) and time-by-diet interaction (P = 0.436). *Post hoc* analyses using pairwise comparisons (and adjusting for multiple comparisons using the Benjamini-Hochberg procedure (33)) showed a significant difference only in the control-fed group between day 0 and day 14 (P = 0.033) (**Fig. 3b**). There were no statistically significant changes in Bacteroidota/Bacillota detected within tomato-fed pigs. The significant Bacteroidota/Bacillota decrease found in the control-fed group at day 14 versus baseline corresponds with the significant decrease in Bacteroidota and increase in Bacillota mentioned above.

These data together suggest that incorporation of tomato into the diet can help prevent the alteration of the microbial profile to maintain higher Bacteroidota/Bacillota ratio, which is considered a more desirable phenotype. Low Bacteroidota/Bacillota ratio in the gut has been linked to an obese host (4, 34, 35), suggesting a higher Bacteroidota/Bacillota ratio is more desirable. In our control pigs, Bacteroidota/Bacillota decreased over time, whereas the ratio remained unchanged in tomato-fed pigs, so it follows that tomato consumption may be playing a role in maintaining a more desirable Bacteroidota/Bacillota ratio. It has been suggested that altering this ratio may directly affect risk of obesity, as there is some evidence that taxa in the Firmicutes phylum have an increased capacity for energy harvest (5, 36). The role of Bacteroidota/Bacillota in predicting or influencing obesity and the mechanisms underlying this relationship, including diet, are worth further investigation.

Diet is known to have a major influence on the gut microbiome in general (6, 8), and limited studies showed that certain dietary patterns or components affect Bacteroidota/Bacillota ratio. Some studies have demonstrated that fiber, starch, and other plant polysaccharides can increase Bacteroidota/Bacillota ratio (8, 37, 38). Tomato powder does provide a source of these carbohydrates, although our control diet was macronutrient matched to the tomato diet, suggesting differences we see here are a function of the small molecule phytochemicals from tomato. Some bacteria are known to metabolize tomato phytochemicals, such as rutin, quercetin, and chlorogenic acid (7, 39). Adding a food with unique phytochemicals to the diet introduces a new source of nutrients for the microbiome and encourages growth of certain bacteria, suggesting the mechanism that phytochemicals indirectly influence the makeup of the microbiome. Effects shown here could be partially or wholly induced by tomato phytochemicals; however, it is also possible that certain polysaccharides in tomatoes provide benefits, preventing the change in Bacteroidota, Bacillota, and Bacteroidota/Bacillota ratio seen in the control-fed animals over time. A study that fed tomato powder to mice with induced liver cancer saw a decreased level of Bacteroidota and an increased level of Bacillota, resulting in a lower Bacteroidota/Bacillota ratio (18). However, these animals were double knockouts deficient in beta carotene oxygenase 1 and 2, which is known to exert physiological affects beyond metabolism of carotenoids, challenging the translation of these results to other mammals (40, 41).

### Several phyla were detected in significantly higher relative abundance in tomato-fed pigs compared to control pigs after 14 days of feeding

In addition to assessing Bacteroidetes and Firmicutes, which were of *a priori* interest, we assessed changes in each of the 45 detected phyla across time points and between diet groups. Differences between relative abundances of individual taxa were determined by compositional analyses using the *ALDEx2* package in R (42– 45). Within control-fed pigs, there were no significant changes in relative abundance of any phylum over time. While we would expect to see differences due to time in Bacteroidota and Bacillota, as was discovered with repeated measures ANOVA, we suspect that due to the multiple testing corrections incurred to test the 45 phyla, this test is conservative in its estimate of changes in taxa relative abundance. Within tomato-fed pigs, 1 phylum (unclassified (Bacteria-derived)) of the 45 detected was significantly altered over time. When comparing diet groups, there were no significant phyla-level differences at day 0, 1 phylum (unclassified (Bacteria-derived)) on day 7, and 5 phyla (Nematoda, Apicomplexa, Deinococcus-Thermus, Pseudomonadota (i.e., Proteobacteria), and unclassified (Bacteria-derived)) on day 14. The relative abundance of each of these phyla was found to be higher in the tomato-fed group than in the control, apart from Deinococcus-Thermus for which the opposite was true. The full list of p-values for all phyla level comparisons can be found in Supplemental Table 5.

No significant differences at day 0 is expected, as no intervention had yet occurred, and microbiome compositions should be relatively consistent between pigs. Providing an explanation for the functional implications of changes in phyla at the other two time points is challenging to describe, as most have not been extensively studied in the context of the gut microbiome and each contain diverse genera and species that vary in function.

To get closer to understanding functional implications of differences in taxa across time points and between diet groups, the same compositional analyses were conducted using *ALDEx2* at the genus level. Significant differences were detected in relative abundances of 4 genera across time in control-fed pigs. These were *Oribacterium, Streptococcus, Lactococcus*, and *Granulicatella*; all of which were detected in a higher relative abundance with time. In tomato-fed pigs compared to control-fed pigs, four genera were found to have significantly increased in relative abundance over time: *Staphylococcus, Alphatorquevirus*, Lambda-like viruses, and an unclassified group (Bacteria-derived).

In the context of the gut microbiome, changes in *Lactococcus* (phylum Firmicutes) and *Staphylococcus* (phylum Firmicutes) abundances is of interest. Some *Lactococcus* species and strains have shown potential to act as a probiotic in the gut and provide some health benefits in animal studies (46, 47). In contrast, this genus has also been associated with body fat accumulation in mice fed a high fat diet (48). More work is needed to determine its exact role. Here we report an increase in *Lactococcus* relative abundance over time within the microbiomes of the control-fed pigs, resulting in a significant difference between diet groups at day 14. Many species within the *Staphylococcus* genus are known to be typical commensal inhabitants of the human and pig skin microbiomes (49, 50). However, there are some species which can cause pathogenesis in humans (51). Without further knowledge of the species present in these samples, it is impossible to say whether increases in *Staphylococcus* abundance in tomato-fed pigs should be viewed as negative. However, it should be noted that no pigs showed signs of diseases throughout the study.

Furthermore, significant differences were assessed between diet groups for each genus. As in phyla-level analyses, no significant differences in abundance of genera were noted between diet groups at day 0. At day 7, an unclassified group (Bacteria-derived) was significantly different between diets, consistent with the single phylum (unclassified (Bacteria-derived) for which a difference was detected in the phyla-level analyses. Analyses of differences at day 14 showed 14 genera significantly different in relative abundance. These were *Alphatorquevirus, Brugia, Loa, Malassezia, Plasmodium, Propionibacterium, Rosiflexus, Saccharomyces, Staphylococcus, Stenotrophomonas, Streptococcus, Vanderwaltozyma*, Lambda-like viruses, and unclassified (Bacteria-derived). All were significantly higher in tomato-fed vs. control group, except for *Rosiflexus* and *Streptococcus*, which were higher in the control group. There is evidence that *Propionibacterium* are early colonizers of the infant gut (52), with their enrichment protective against necrotizing enterocolitis (53), and acting a probiotic (54). Similarly, some *Saccharomyces* species have also been shown to be probiotic, increasing the abundance of Bacteroidota and decreasing Bacilotta (55), while others act along the gut-brain axis in reducing irritable bowel disease severity (56). Increased *Streptococcus* has been associated with increased localized inflammation (57), while other strains have been shown to be probiotic (58). However, it is currently difficult to contextualizing these findings because of the diversity of species within each genus. The full list of p-values for all genera level comparisons can be found in Supplemental Table 6.

### Tomato-fed pigs had a significantly higher fecal microbiome alpha diversity at a phylum, but not at a genus levels

The microbiome is a complex collection of organisms, so it is important to analyze differences in the community based not only on single phyla and genera, but also by examining the overall diversity present. Therefore, using the Shannon index, alpha-diversity was calculated at the phyla-and genera-level for each sample to provide a measure of taxonomic diversity within each sample. Diet and time point group averages were then compared with a repeated measures ANOVA (**FIG. 4**).

**FIG 4.**
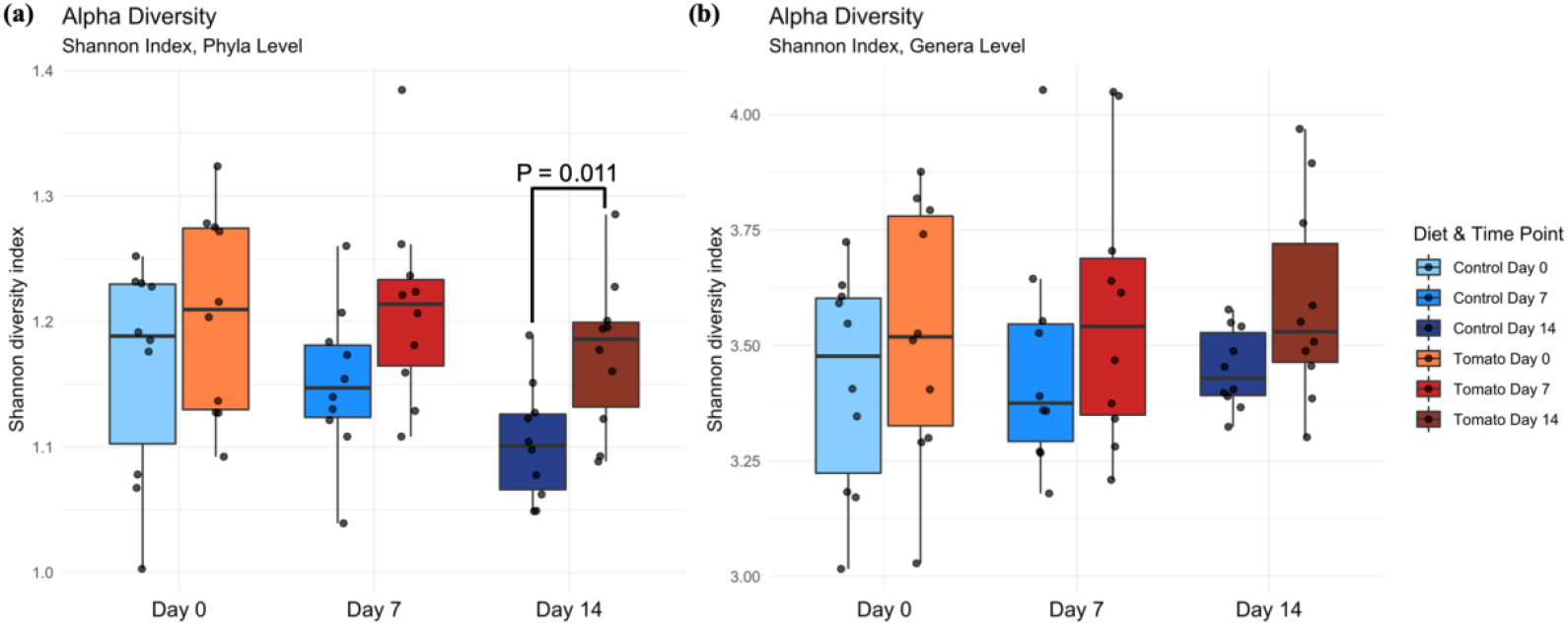
Alpha diversity as measured by the Shannon diversity index at the (a) phyla and (b) genera level. (a) A statistically significant effect of diet was found via repeated measures ANOVA (P = 0.004), and a *post hoc* difference (P = 0.011) was found at the phyla level between control and tomato fed pigs at day 14. (b) No significant differences were observed at the genera level.

Comparison of phyla-level alpha-diversity between diets and time points showed a significant effect of diet on alpha-diversity (P = 0.004) but no significant effect of time (P = 0.086) or diet-by-time interaction (P = 0.791). *Post hoc* analyses by pairwise comparison revealed a statistical difference between control-and tomato-fed pigs at day 14 (P = 0.011), with higher alpha-diversity in the tomato-fed animals (**Fig. 4a**). This aligns with our univariate *ALDEx2* analyses, as significant differences in 5 phyla were observed between the diets at day 14. Consumption of tomato has previously been shown to affect alpha-diversity. Mice consuming high-fat diets supplemented with tomato powder had higher levels of alpha-diversity than those who did not consume tomato powder (17, 18). Higher alpha-diversity is desirable, as a more diverse gut microbiome has been associated with more benefits for the host and better resilience to pathogens (25).

The repeated measures ANOVA investigating the effect of diet, time point, and their interaction on alpha-diversity at the genera level showed no significant differences (**Fig. 4b**). The lack of observed effect has been similarly noted in human interventions with single foods, including broccoli (59). Another study showed that walnut consumption significantly increased alpha-diversity in rats (60). Again, few studies have been conducted with single plant food interventions for comparison to our results here.

The gut microbiome has a large amount of functional redundancy at the genera and species level, meaning multiple microorganisms contribute the same metabolic functions (25). For example, there are numerous different organisms, when annotated at the genus level, that metabolize carbohydrates, others that metabolize proteins, and some that overlap and metabolize both macromolecules. This provides stability and resiliency to the microbial ecosystem of the gut through a consistent use of nutrients and output of metabolites, even if the exact genera or species presence is changing. Dietary causes of change in alpha-diversity typically occur from repeated habits or patterns that are sustained and dominated by one macronutrient, such as consistent high fat intake, because this limits the available nutrients for microbes (25).

In summary, we have found that supplementation of the diet with 10% tomato powder (as compared to a macronutrient-matched control) has the ability to modulate the gut microbiome in pigs. Animals on tomato-containing diets had higher alpha diversity, a higher Bacteroidota/Bacillota ratio, higher abundance of Bacteroidota (i.e., Bacteroidetes), and lower abundance of Bacillota (i.e., Firmicutes), consistent with a more beneficial microbial phenotype. The effect of tomato consumption on the gut microbiome in humans warrants further investigation at a functional level to improve the understanding of the effect of tomato-rich diet on functional resilience of human gut microbiome.

## METHODS

### Experimental Diet Production

Processing tomatoes (*Solanum lycopersicum* L.) used in this study were grown at the North Central Agricultural Research Station of Ohio State University (OSU) in Fremont, OH. A hybrid tomato derived from the cross OH8245 × OH8243 (61)was used. Tomatoes were grown using conventional horticultural practices, mechanically harvested using a Guaresci harvester (Guaresci, Sp.A, Pilastri, Italy), and sorted to include ripe fruits only. Tomatoes were transported to the Columbus, OH, campus of OSU and processed at the Food Industries Center Pilot Plant, where fruits were immediately washed, diced, and frozen, as previously described (62). Frozen tomatoes were freeze-dried and dry material ground into a fine powder using a vertical chopper mixer (62). Tomato powder was stored in vacuum sealed bags at -20 ° C until use.

The basal diet (**Table 2**) was formulated with a nutrient make-up appropriate for nursery pigs weighing 7-11 kg according to the National Research Council (63). To the basal diet, the tomato powder was added at 10% *w*/*w*. To create the control diet, the basal diet was supplemented with milk protein isolate (90% purity, 13%, protein), powdered sugar (70%, sugar), pectin (3.4%, soluble fiber) and cellulose (13.6%, insoluble fiber) to create a macronutrient match to the tomato diet (**Table 2**). These ingredients were formulated to match the ratios of nutrients typically found in tomato powder as reported by FoodData Central (64). This supplement was added at 10% *w*/*w* to match the addition of the tomato powder.

**TABLE 2.**
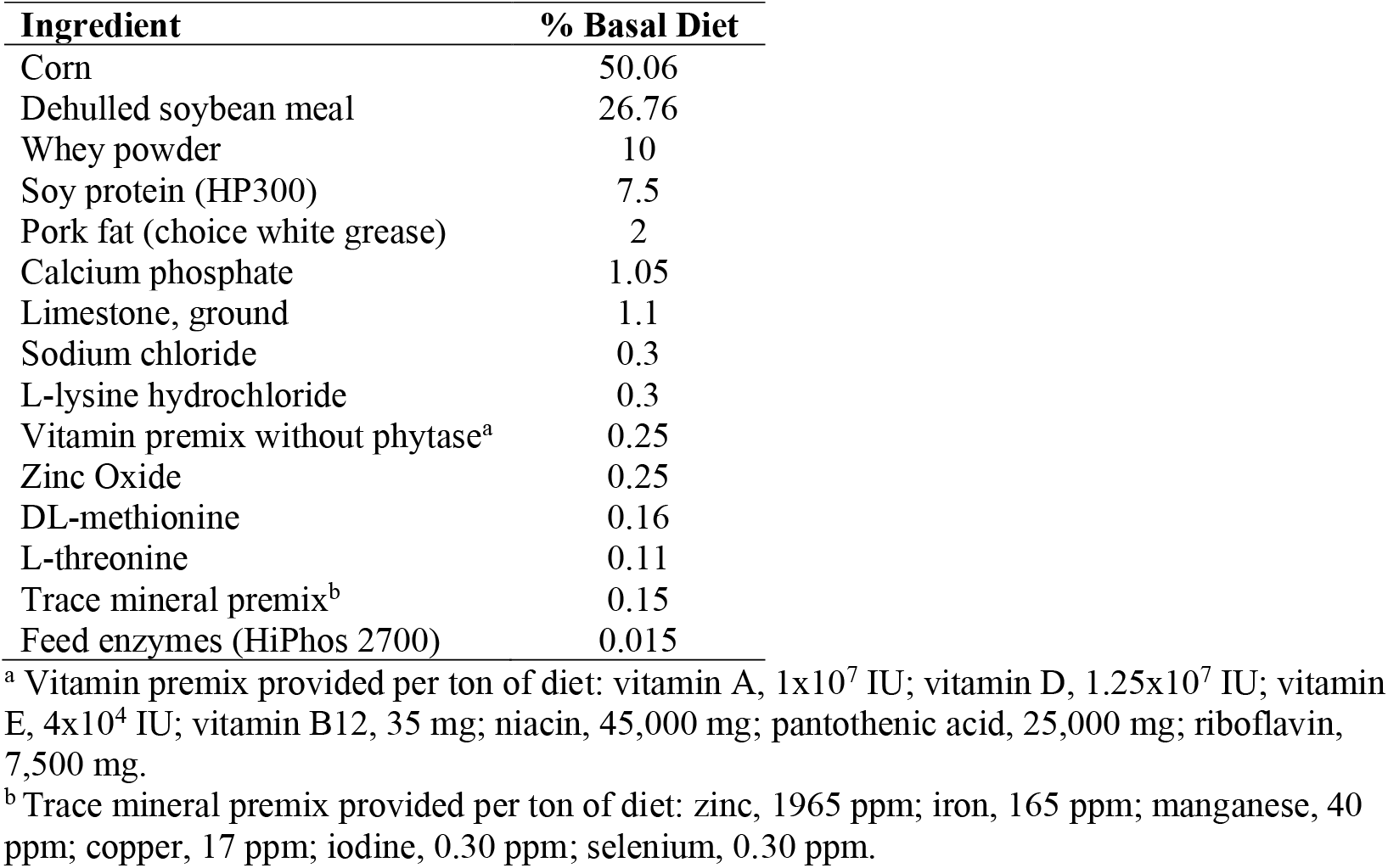
Composition of basal diet on an as-fed basis. This diet delivered 3,381 kcal/kg, 22.7& crude protein, 1.35% standardized ileal digestible lysine, 34% ileal digestible methionine:lysine, 57% ileal digestible methionine and cysteine:lysine, 0.8% calcium, and 0.67% phosphate.

| Ingredient | % Basal Diet |
| --- | --- |
| Corn | 50.06 |
| Dehulled soybean meal | 26.76 |
| Whey powder | 10 |
| Soy protein (HP300) | 7.5 |
| Pork fat (choice white grease) | 2 |
| Calcium phosphate | 1.05 |
| Limestone, ground | 1.1 |
| Sodium chloride | 0.3 |
| L-lysine hydrochloride | 0.3 |
| Vitamin premix without phytase<sup>a</sup> | 0.25 |
| Zinc Oxide | 0.25 |
| DL-methionine | 0.16 |
| L-threonine | 0.11 |
| Trace mineral premix<sup>b</sup> | 0.15 |
| Feed enzymes (HiPhos 2700) | 0.015 |

<sup>a</sup> Vitamin premix provided per ton of diet: vitamin A, $1 \times 10^7$ IU; vitamin D, $1.25 \times 10^7$ IU; vitamin E, $4 \times 10^4$ IU; vitamin B12, 35 mg; niacin, 45,000 mg; pantothenic acid, 25,000 mg; riboflavin, 7,500 mg.
<sup>b</sup> Trace mineral premix provided per ton of diet: zinc, 1965 ppm; iron, 165 ppm; manganese, 40 ppm; copper, 17 ppm; iodine, 0.30 ppm; selenium, 0.30 ppm.

### Animal Study Design

Twenty male pigs born to six sows in summer 2019 at the OSU Swine Facility in Dublin, OH were used in this study. Male pigs were selected to allow sampling of prostatic tissue for a secondary study. At weaning twenty male pigs were selected according to weight and randomly assigned to dietary treatment. A scheme of the overall study design can be found in **Fig. 1**.

To prevent diet mixing and cross-contamination of microbiomes through contact, only pigs consuming the same diets were allowed to have contact. The two diet groups were housed across the room from each other and divided by a walkway. Pens had sufficient space between railings for nose-to-nose contact with other pigs, though not enough space to allow a pig to leave its own pen. After successful weaning from mother’s milk, all pigs consumed the basal diet to acclimate to solid food from week 3 to 4. Pigs at 4 weeks of age began consuming the experimental diets assigned. Feeders were attached to the front of the pens and allowed pigs to eat *ad libitum*. Pigs were weighed weekly to monitor growth and were checked daily to ensure health. Apart from feeding, weighing, and swabbing, human contact with pigs was minimized to limit influences on the gut microbiome of pigs. This study was approved by the OSU Office of Responsible Research Practices (IACUC #2019A00000060).

### Sample Collection

The microbiome was sampled 3 times during this study via rectal swabs: prior to beginning experimental diets (day 0, aged 4 weeks), after one week of consuming assigned diets (day 7, the study midpoint, aged 5 weeks), and after two weeks of dietary intervention (day 14, end of study, aged 6 weeks) (**Fig. 1**). Swabs used for collection were sterile DNA/RNA Shield Collection Tubes (Zymo Research, Irvine, CA, United States) and were stored at -80 °C after collection prior to sequencing.

### Sample Processing and Sequencing

Swabs were sent to CosmosID, Inc. (Rockville, MD, United States) for DNA extraction and sequencing. Samples were sequenced via 150 bp paired-end shotgun sequencing, using an Illumina HiSeq4000 instrument (San Diego, CA, United States). Unopened collection tubes were used as negative controls. Samples with reads lower than 1.8M reads were re-sequenced and merged with the prior sequences, allowing increased microbiome coverage.

### Quality of Sequences

Quality of sequences was analyzed using FastQC version 0.11.9 (23). Sequences were trimmed during annotation in MG-RAST version 4.0.3 (65) if they contained more than 5 bases that were below a minimum Phred quality score of 20. Full metadata for MG-RAST parameters can be found at https://www.mg-rast.org/linkin.cgi?project=mgp93233.

### Sequence analyses and taxonomy identification

Raw fastq files were made publicly available via the NCBI Sequence Read Archive (SRA), project number PRJNA601162. Annotated files are available through MG-RAST (project mgp93233), and annotated taxa can be found in the Supplementary Tables S3 and S4. Sample reads were annotated via the MG-RAST open-access pipeline (65) using the RefSeq database (66). No assembly was completed prior to annotation. Sequences were screened for host DNA using the NCBI *Sus scrofa* v10.2 genome and, if identified, were removed. Sequences from Bacteria, Archaea, Eukaryota, and viruses were kept for further analysis. Phyla and genera were filtered to exclude taxa that were present in less than 67% of tested samples.

### Statistical analysis

All data analysis was performed in R version 4.0.3 (67) using RStudio (68) and results were considered significant at P ≤ 0.05. All code used to conduct analyses can be found in the tomato-pig-microbiome repository at www.github.com/CooperstoneLab. All figures were created using *ggplot2* (69). Microbiome profiles at both the phyla and genera taxonomic level were analyzed. Data was normalized using relative abundance to account for differences in sequencing depth, since rarefaction is no longer recommended as a normalization tool due to high potential for data loss (70). Relative abundance was calculated by dividing the number of counts for any one taxon by the total number of counts at that taxonomic level per sample.

Interactive Krona plots (Fig. S1) were created using R packages *phyloseq* (71) and *psadd* (72) to visualize the microbiome composition. To assess sufficiency of sequencing depth, rarefaction curves were created using the package *ranacapa* (73) with a window size of 60,000 counts (**Fig. S2**).

To understand overall microbiome differences between diet groups and across time points, beta diversity was calculated using the R package *vegan* and functions “vegdist” and “cmdscale” then visualized using PCoA with a Bray-Curtis dissimilarity matrix. Significance of separation between treatments was tested via restricted permutation tests using Permutational Multivariate Analysis of Variance (PERMANOVA) (74) with the R package *vegan* using the function “adonis2” (75) and the “how” function from the package *permute* (76) (model: Beta Diversity ∼ Diet + Time Point + Diet×Time Point + Error where each pig was a plot containing 3 samples collected over time). The argument “by” was set to “margin” to assess how much each individual term contributes to the model. The permutations were restricted within each pig as a time series for which the same permutation was used across pigs (R code available in supplemental data).

To examine differences in relative abundances of individual microorganisms across groups, univariate analyses were conducted using the R package *ALDEx2* (42–44). This specific package was used because it is designed to analyze high-throughput sequence data as compositional data (i.e., it accounts for total reads and uses a data transformation for statistical testing), allowing direct comparison of samples without an effect of total number of reads (43, 45). Raw taxa counts (as compared to relative abundance data) were used and center log ratio (CLR) transformed for these analyses (42, 43). Parametric tests were used for these analyses as our data met assumptions for normality.

Alpha-diversity of each sample was calculated from counts using the Shannon index in the R package *vegan* with the function “diversity” (75). The Shannon index alpha-diversity group means were compared using repeated measures two-way ANOVA (model: Alpha-Diversity ∼ Diet + Time Point + Diet×Time Point + Error). *Post hoc* analyses for significant model terms were completed using pairwise comparison via *t*-test to determine where differences originated.

The ratio of the phyla Bacteroidota (i.e., Bacteroidetes) to Bacilotta (i.e., Firmicutes) was determined for each sample by dividing relative abundance of Bacteroidota by that of Bacilotta, each as a percentage of the total phyla. Differences between the ratios were tested between diets and time points using two-way repeated measures ANOVA given our *a priori* interest in these phyla, followed by a pairwise comparison via *t*-test as a *post hoc* analysis. Additionally, the relative abundance of Bacteroidota and Bacilotta phyla were separately tested using two-way repeated measured ANOVA with a post-hoc test of pairwise comparison by *t*-test.

## Supporting information

Kronas plots

## ACKNOWLEDGMENTS

This research was financially supported by USDA Hatch funds (OHO01470 and PEN04646/Accession 1015787), USDA-NIFA National Needs Fellowship (2014-38420-21844), Ohio Agricultural Research and Development Center Seed Grant (to MLG), and Foods for Health, a focus area of the Discovery Themes Initiative at The Ohio State University. The funders had no role in study, design, data collection and interpretation, or the decision to submit the work for publication. The authors thank to M. Laura Rolon for providing the starting R code for microbiome data analyses.

## AUTHOR CONTRIBUTIONS

Conception and study design: MLG, SKJ, JK, and JLC; Data collection: MLG, DMF, SKJ, JLC; Data analysis and interpretation: MLG, EAB, CQ-M, JK, JLC; Drafting the article: MLG, EAB, JLC; Revision and review of the article: MLG, EAB, CQ-M, DMF, SKJ, JK, JLC; Responsibility for final content: JLC.

## SUPPLEMENTAL MATERIAL

### Supplemental Figures

**SUPPLEMENTAL FIGURE 1.**
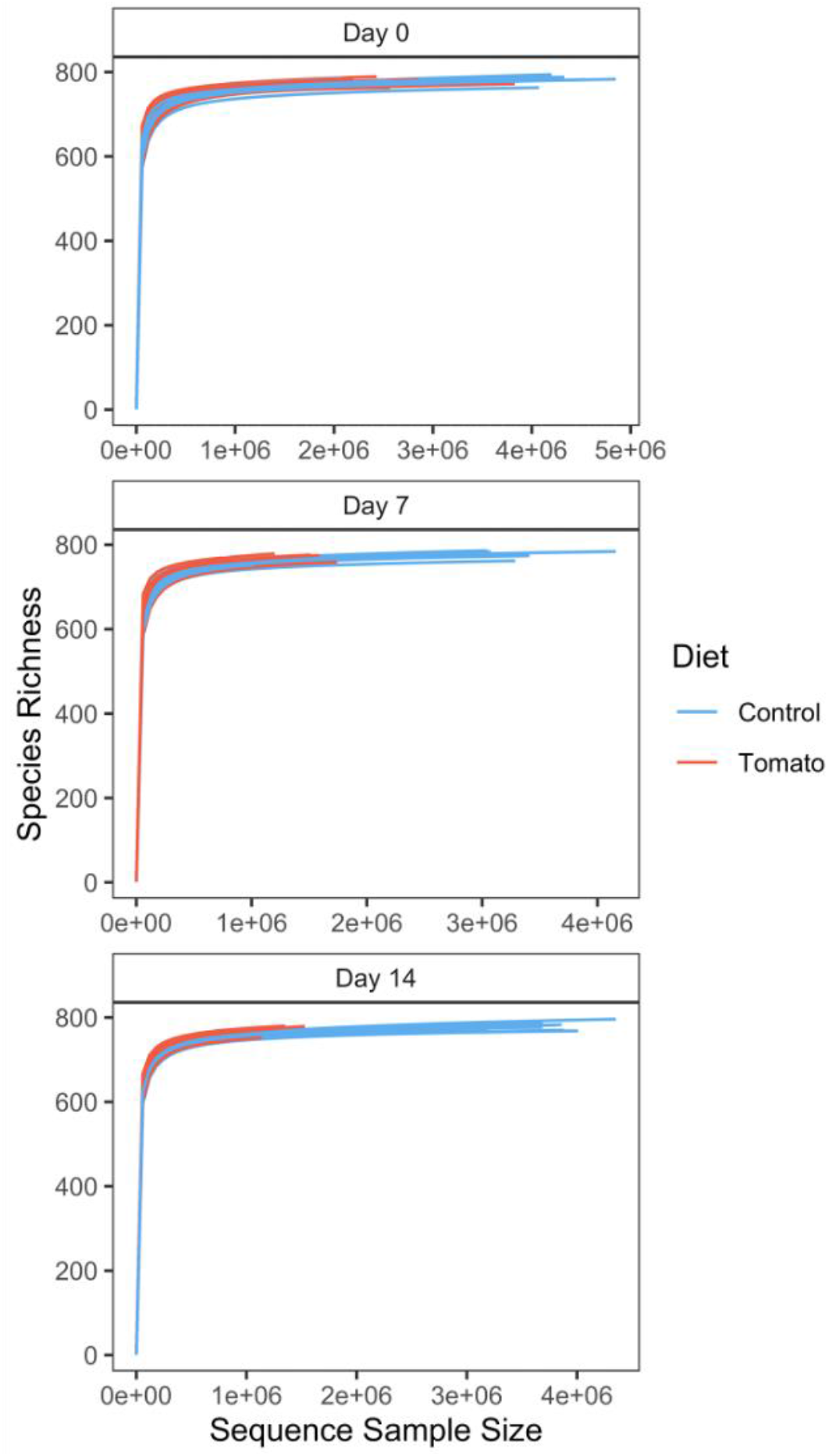
Rarefaction curves showing species richness relative to sequence sample size, by sampling day and diet.

Supplemental Tables: Provided in Excel document: Goggans_etal_2021_tomato_pig_microbiome_WGS

**SUPPLEMENTAL TABLE 1**. Weights (kg) of pigs at study day 0 (aged 4 weeks), study day 7 (aged 5 weeks), and study day 14 (aged 6 weeks). There were no significant differences between diets at any time point by unpaired t-tests.

**SUPPLEMENTAL TABLE 2**. Sample metadata, containing full sample name and each variable.

**SUPPLEMENTAL TABLE 3**. Taxonomic identification annotated at the phyla level via MG-RAST.

**SUPPLEMENTAL TABLE 4**. Taxonomic identification annotation at the genera level via MG-RAST.

**SUPPLEMENTAL TABLE5**. Output from ALDEeX2 univariate analysis at the phyla level, significant taxa after a multiple testing correction are indicated with a yellow highlight. Abbreviations: rab.all: median clr value for all samples in the feature; rab.win.Control: median clr value for the control group; rab.win.Tomato: median clr value for the tomato group; dif.btw: median difference in clr values between S and NS groups; diff.btw: median difference in clr values between tomato and control groups; diff.win: median of the largest difference in clr values within tomato and control groups; overlap: proportion of effect size that overlaps 0 (i.e. no effect); we.ep: Expected P value of Welch’s t test; we.eBH: Expected Benjamini-Hochberg corrected P value of Welch’s t test; wi.ep: Expected P value of Wilcoxon rank test; wi.eBH: Expected Benjamini-Hochberg corrected P value of Wilcoxon test; kw.ep: Expected P value of Kruskal-Wallace test; kw.eBH: Expected Benjamini-Hochberg corrected P value of Kruskal-Wallace test; glm.ep: Expected P value of glm test; glm.eBH: Expected Benjamini-Hochberg corrected P value of glm test

**SUPPLEMENTAL TABLE 6**. Output from ALDEeX2 univariate analysis at the genera level, significant taxa after a multiple testing correction are indicated with a yellow highlight. Abbreviations: rab.all: median clr value for all samples in the feature; rab.win.Control: median clr value for the control group; rab.win.Tomato: median clr value for the tomato group; dif.btw: median difference in clr values between S and NS groups; diff.btw: median difference in clr values between tomato and control groups; diff.win: median of the largest difference in clr values within tomato and control groups; overlap: proportion of effect size that overlaps 0 (i.e. no effect); we.ep: Expected P value of Welch’s t test; we.eBH: Expected Benjamini-Hochberg corrected P value of Welch’s t test; wi.ep: Expected P value of Wilcoxon rank test; wi.eBH: Expected Benjamini-Hochberg corrected P value of Wilcoxon test; kw.ep: Expected P value of Kruskal-Wallace test; kw.eBH: Expected Benjamini-Hochberg corrected P value of Kruskal-Wallace test; glm.ep: Expected P value of glm test; glm.eBH: Expected Benjamini-Hochberg corrected P value of glm test

## REFERENCES

1. Park W. 2018. Gut microbiomes and their metabolites shape human and animal health. J Microbiol 56:151–153.

2. Battson ML, Lee DM, Weir TL, Gentile CL. 2018. The gut microbiota as a novel regulator of cardiovascular function and disease. The Journal of Nutritional Biochemistry 56:1–15.

3. Tuddenham S, Sears CL. 2015. The intestinal microbiome and health: Current Opinion in Infectious Diseases 28:464–470.

4. Ley RE, Turnbaugh PJ, Klein S, Gordon JI. 2006. Human gut microbes associated with obesity. 7122. Nature 444:1022–1023.

5. Turnbaugh PJ, Ley RE, Mahowald MA, Magrini V, Mardis ER, Gordon JI. 2006. An obesity-associated gut microbiome with increased capacity for energy harvest. Nature 444:1027–1031.

6. Jeffery I, O’Toole P. 2013. Diet-Microbiota Interactions and Their Implications for Healthy Living. Nutrients 5:234–252.

7. Tuohy KM, Conterno L, Gasperotti M, Viola R. 2012. Up-regulating the Human Intestinal Microbiome Using Whole Plant Foods, Polyphenols, and/or Fiber. J Agric Food Chem 60:8776–8782.

8. Turnbaugh PJ, Ridaura VK, Faith JJ, Rey FE, Knight R, Gordon JI. 2009. The Effect of Diet on the Human Gut Microbiome: A Metagenomic Analysis in Humanized Gnotobiotic Mice. Science Translational Medicine 1:6ra14–6ra14.

9. Magne F, Gotteland M, Gauthier L, Zazueta A, Pesoa S, Navarrete P, Balamurugan R. 2020. The Firmicutes/Bacteroidetes Ratio: A Relevant Marker of Gut Dysbiosis in Obese Patients? Nutrients 12:1474.

10. Kopf JC, Suhr MJ, Clarke J, Eyun S, Riethoven J-JM, Ramer-Tait AE, Rose DJ. 2018. Role of whole grains versus fruits and vegetables in reducing subclinical inflammation and promoting gastrointestinal health in individuals affected by overweight and obesity: a randomized controlled trial. Nutr J 17:72.

11. United States Department of Agriculture, Economic Research Service. 2019. Potatoes and tomatoes are the most commonly consumed vegetables.

12. FAOSTAT. 2016. Food and Agriculture Organization of the United Nations. FAO, Rome, Italy. http://www.fao.org/faostat/en/#data.

13. USDA Economic Research Service. 2018. Vegetables and Pulses Yearbook Tables.

14. Sesso HD, Liu S, Gaziano JM, Buring JE. 2003. Dietary Lycopene, Tomato-Based Food Products and Cardiovascular Disease in Women. The Journal of Nutrition 133:2336–2341.

15. Burton-Freeman BM, Sesso HD. 2014. Whole Food versus Supplement: Comparing the Clinical Evidence of Tomato Intake and Lycopene Supplementation on Cardiovascular Risk Factors. Advances in Nutrition 5:457–485.

16. Rowles JL, Ranard KM, Applegate CC, Jeon S, An R, Erdman JW. 2018. Processed and raw tomato consumption and risk of prostate cancer: a systematic review and dose– response meta-analysis. Prostate Cancer Prostatic Dis 21:319–336.

17. Li C-C, Liu C, Fu M, Hu K-Q, Aizawa K, Takahashi S, Hiroyuki S, Cheng J, von Lintig J, Wang X-D. 2018. Tomato Powder Inhibits Hepatic Steatosis and Inflammation Potentially Through Restoring SIRT1 Activity and Adiponectin Function Independent of Carotenoid Cleavage Enzymes in Mice. Mol Nutr Food Res 62:1700738.

18. Xia H, Liu C, Li C-C, Fu M, Takahashi S, Hu K-Q, Aizawa K, Hiroyuki S, Wu G, Zhao L, Wang X-D. 2018. Dietary Tomato Powder Inhibits High-Fat Diet–Promoted Hepatocellular Carcinoma with Alteration of Gut Microbiota in Mice Lacking Carotenoid Cleavage Enzymes. Cancer Prev Res 11:797–810.

19. Scarano A, Butelli E, De Santis S, Cavalcanti E, Hill L, De Angelis M, Giovinazzo G, Chieppa M, Martin C, Santino A. 2018. Combined Dietary Anthocyanins, Flavonols, and Stilbenoids Alleviate Inflammatory Bowel Disease Symptoms in Mice. Front Nutr 4:75.

20. Liso M, De Santis S, Scarano A, Verna G, Dicarlo M, Galleggiante V, Campiglia P, Mastronardi M, Lippolis A, Vacca M, Sobolewski A, Serino G, Butelli E, De Angelis M, Martin C, Santino A, Chieppa M. 2018. A Bronze-Tomato Enriched Diet Affects the Intestinal Microbiome under Homeostatic and Inflammatory Conditions. Nutrients 10:1862.

21. Ephraim E, Jewell D. 2021. The Influence of Fiber Source on Circulating Concentration of Age-Related Metabolites and the Gut Microbial Composition in Senior Cats. Current Developments in Nutrition 5:13–13.

22. Quince C, Walker AW, Simpson JT, Loman NJ, Segata N. 2017. Shotgun metagenomics, from sampling to analysis. Nat Biotechnol 35:833–844.

23. Andrews S. 2010. FastQC: A Quality Control Tool for High Throughput Sequence Data. http://www.bioinformatics.babraham.ac.uk/projects/fastqc.

24. Hillmann B, Al-Ghalith GA, Shields-Cutler RR, Zhu Q, Gohl DM, Beckman KB, Knight R, Knights D. 2018. Evaluating the Information Content of Shallow Shotgun Metagenomics 3:12.

25. Lozupone CA, Stombaugh JI, Gordon JI, Jansson JK, Knight R. 2012. Diversity, stability and resilience of the human gut microbiota. Nature 489:220–230.

26. Kim HB, Borewicz K, White BA, Singer RS, Sreevatsan S, Tu ZJ, Isaacson RE. 2011. Longitudinal investigation of the age-related bacterial diversity in the feces of commercial pigs. Veterinary Microbiology 153:124–133.

27. Xiao L, Estellé J, Kiilerich P, Ramayo-Caldas Y, Xia Z, Feng Q, Liang S, Pedersen AØ, Kjeldsen NJ, Liu C, Maguin E, Doré J, Pons N, Le Chatelier E, Prifti E, Li J, Jia H, Liu X, Xu X, Ehrlich SD, Madsen L, Kristiansen K, Rogel-Gaillard C, Wang J. 2016. A reference gene catalogue of the pig gut microbiome. Nat Microbiol 1:16161.

28. Alain B. Pajarillo E, Chae J-P, P. Balolong M, Bum Kim H, Kang D-K. 2014. Assessment of fecal bacterial diversity among healthy piglets during the weaning transition. J Gen Appl Microbiol 60:140–146.

29. David LA, Maurice CF, Carmody RN, Gootenberg DB, Button JE, Wolfe BE, Ling AV, Devlin AS, Varma Y, Fischbach MA, Biddinger SB, Dutton RJ, Turnbaugh PJ. 2014. Diet rapidly and reproducibly alters the human gut microbiome. Nature 505:559–563.

30. Ziegler A, Gonzalez L, Blikslager A. 2016. Large Animal Models: The Key to Translational Discovery in Digestive Disease Research. Cellular and Molecular Gastroenterology and Hepatology 2:716–724.

31. Ridaura VK, Faith JJ, Rey FE, Cheng J, Duncan AE, Kau AL, Griffin NW, Lombard V, Henrissat B, Bain JR, Muehlbauer MJ, Ilkayeva O, Semenkovich CF, Funai K, Hayashi DK, Lyle BJ, Martini MC, Ursell LK, Clemente JC, Van Treuren W, Walters WA, Knight R, Newgard CB, Heath AC, Gordon JI. 2013. Gut Microbiota from Twins Discordant for Obesity Modulate Metabolism in Mice. Science 341:1241214.

32. Hildebrandt MA, Hoffmann C, Sherrill–Mix SA, Keilbaugh SA, Hamady M, Chen Y, Knight R, Ahima RS, Bushman F, Wu GD. 2009. High-Fat Diet Determines the Composition of the Murine Gut Microbiome Independently of Obesity. Gastroenterology 137:1716-1724.e2.

33. Benjamini Y, Hochberg Y. 1995. Controlling the False Discovery Rate: A Practical and Powerful Approach to Multiple Testing. Journal of the Royal Statistical Society: Series B (Methodological) 57:289–300.

34. Ismail NA, Ragab SH, ElBaky AA, Shoeib ARS, Alhosary Y, Fekry D. 2011. Frequency of Firmicutes and Bacteroidetes in gut microbiota in obese and normal weight Egyptian children and adults. aoms 3:501–507.

35. Koliada A, Syzenko G, Moseiko V, Budovska L, Puchkov K, Perederiy V, Gavalko Y, Dorofeyev A, Romanenko M, Tkach S, Sineok L, Lushchak O, Vaiserman A. 2017. Association between body mass index and Firmicutes/Bacteroidetes ratio in an adult Ukrainian population. BMC Microbiol 17:120.

36. Backhed F, Ding H, Wang T, Hooper LV, Koh GY, Nagy A, Semenkovich CF, Gordon JI. 2004. The gut microbiota as an environmental factor that regulates fat storage. Proceedings of the National Academy of Sciences 101:15718–15723.

37. Barczynska R, Slizewska K, Litwin M, Szalecki M, Zarski A, Kapusniak J. 2015. The effect of dietary fibre preparations from potato starch on the growth and activity of bacterial strains belonging to the phyla Firmicutes, Bacteroidetes, and Actinobacteria. Journal of Functional Foods 19:661–668.

38. De Filippo C, Cavalieri D, Di Paola M, Ramazzotti M, Poullet JB, Massart S, Collini S, Pieraccini G, Lionetti P. 2010. Impact of diet in shaping gut microbiota revealed by a comparative study in children from Europe and rural Africa. Proceedings of the National Academy of Sciences 107:14691–14696.

39. Edwards CA, Havlik J, Cong W, Mullen W, Preston T, Morrison DJ, Combet E. 2017. Polyphenols and health: Interactions between fibre, plant polyphenols and the gut microbiota. Nutr Bull 42:356–360.

40. Guo X, Wu L, Lyu Y, Chowanadisai W, Clarke SL, Lucas EA, Smith BJ, He H, Wang W, Medeiros DM, Lin D. 2017. Ablation of β,β-carotene-9′,10′-oxygenase 2 remodels the hypothalamic metabolome leading to metabolic disorders in mice. The Journal of Nutritional Biochemistry 46:74–82.

41. Wu L, Guo X, Hartson SD, Davis MA, He H, Medeiros DM, Wang W, Clarke SL, Lucas EA, Smith BJ, von Lintig J, Lin D. 2017. Lack of β, β-carotene-9′, 10′-oxygenase 2 leads to hepatic mitochondrial dysfunction and cellular oxidative stress in mice. Mol Nutr Food Res 61:1600576.

42. Fernandes AD, Reid JN, Macklaim JM, McMurrough TA, Edgell DR, Gloor GB. 2014. Unifying the analysis of high-throughput sequencing datasets: characterizing RNA-seq, 16S rRNA gene sequencing and selective growth experiments by compositional data analysis. Microbiome 2:15.

43. Fernandes AD, Macklaim JM, Linn TG, Reid G, Gloor GB. 2013. ANOVA-Like Differential Expression (ALDEx) Analysis for Mixed Population RNA-Seq. PLoS ONE 8:e67019.

44. Gloor GB, Macklaim JM, Fernandes AD. 2016. Displaying Variation in Large Datasets: Plotting a Visual Summary of Effect Sizes. Journal of Computational and Graphical Statistics 25:971–979.

45. Gloor GB, Macklaim JM, Pawlowsky-Glahn V, Egozcue JJ. 2017. Microbiome Datasets Are Compositional: And This Is Not Optional. Front Microbiol 8:2224.

46. Berlec A, Perše M, Ravnikar M, Lunder M, Erman A, Cerar A, Štrukelj B. 2017. Dextran sulphate sodium colitis in C57BL/6J mice is alleviated by Lactococcus lactis and worsened by the neutralization of Tumor necrosis Factor α. International Immunopharmacology 43:219–226.

47. Kimoto-Nira H, Mizumachi K, Nomura M, Kobayashi M, Fujita Y, Okamoto T, Suzuki I, Tsuji NM, Kurisaki J, Ohmomo S. 2007. Lactococcus sp. as Potential Probiotic Lactic Acid Bacteria. JARQ 41:181–189.

48. Parks BW, Nam E, Org E, Kostem E, Norheim F, Hui ST, Pan C, Civelek M, Rau CD, Bennett BJ, Mehrabian M, Ursell LK, He A, Castellani LW, Zinker B, Kirby M, Drake TA, Drevon CA, Knight R, Gargalovic P, Kirchgessner T, Eskin E, Lusis AJ. 2013. Genetic Control of Obesity and Gut Microbiota Composition in Response to High-Fat, High-Sucrose Diet in Mice. Cell Metabolism 17:141–152.

49. Costello EK, Lauber CL, Hamady M, Fierer N, Gordon JI, Knight R. 2009. Bacterial Community Variation in Human Body Habitats Across Space and Time. Science 326:1694–1697.

50. Kloos WE. 1980. Natural Populations of the Genus Staphylococcus. Annu Rev Microbiol 34:559–592.

51. Lowy FD. 1998. Staphylococcus aureus Infections. The New England Journal of Medicine 13.

52. Rocha Martin VN, Schwab C, Krych L, Voney E, Geirnaert A, Braegger C, Lacroix C. 2019. Colonization of Cutibacterium avidum during infant gut microbiota establishment. FEMS Microbiology Ecology 95.

53. Colliou N, Ge Y, Sahay B, Gong M, Zadeh M, Owen JL, Neu J, Farmerie WG, Alonzo F, Liu K, Jones DP, Li S, Mohamadzadeh M. 2017. Commensal Propionibacterium strain UF1 mitigates intestinal inflammation via Th17 cell regulation. Journal of Clinical Investigation 127:3970–3986.

54. Lan A, Bruneau A, Philippe C, Rochet V, Rouault A, Hervé C, Roland N, Rabot S, Jan G. 2007. Survival and metabolic activity of selected strains of Propionibacterium freudenreichii in the gastrointestinal tract of human microbiota-associated rats. Br J Nutr 97:714–724.

55. Yu L, Zhao X, Cheng M, Yang G, Wang B, Liu H, Hu Y, Zhu L, Zhang S, Xiao Z, Liu Y, Zhang B, Mu M. 2017. Saccharomyces boulardii Administration Changes Gut Microbiota and Attenuates D-Galactosamine-Induced Liver Injury. Sci Rep 7:1359.

56. Constante M, De Palma G, Lu J, Jury J, Rondeau L, Caminero A, Collins SM, Verdu EF, Bercik P. 2021. Saccharomyces boulardii CNCM I-745 modulates the microbiota–gut– brain axis in a humanized mouse model of Irritable Bowel Syndrome. Neurogastroenterology & Motility 33.

57. Boer CG, Radjabzadeh D, Medina-Gomez C, Garmaeva S, Schiphof D, Arp P, Koet T, Kurilshikov A, Fu J, Ikram MA, Bierma-Zeinstra S, Uitterlinden AG, Kraaij R, Zhernakova A, van Meurs JBJ. 2019. Intestinal microbiome composition and its relation to joint pain and inflammation. Nat Commun 10:4881.

58. Iyer R, Tomar SK, Kapila S, Mani J, Singh R. 2010. Probiotic properties of folate producing Streptococcus thermophilus strains. Food Research International 43:103–110.

59. Kaczmarek JL, Liu X, Charron CS, Novotny JA, Jeffery EH, Seifried HE, Ross SA, Miller MJ, Swanson KS, Holscher HD. 2019. Broccoli consumption affects the human gastrointestinal microbiota. The Journal of Nutritional Biochemistry 63:27–34.

60. Byerley LO, Samuelson D, Blanchard E, Luo M, Lorenzen BN, Banks S, Ponder MA, Welsh DA, Taylor CM. 2017. Changes in the gut microbial communities following addition of walnuts to the diet. The Journal of Nutritional Biochemistry 48:94–102.

61. Berry SZ, Gould WA, Wiese KL. 1991. “Ohio 8245” Processing Tomato. HortScience 26:1083.

62. Cooperstone JL, Tober KL, Riedl KM, Teegarden MD, Cichon MJ, Francis DM, Schwartz SJ, Oberyszyn TM. 2017. Tomatoes protect against development of UV-induced keratinocyte carcinoma via metabolomic alterations. Sci Rep 7:5106.

63. National Research Council. 2012. Nutrient Requirements of Swine: Eleventh Revised Edition 11th edition. The National Academies PRess, shinton, DC.

64. 2018. Tomato powder. SR Legacy. United States Department of Agriculture.

65. Meyer F, Paarmann D, D’Souza M, Olson R, Glass E, Kubal M, Paczian T, Rodriguez A, Stevens R, Wilke A, Wilkening J, Edwards R. 2008. The metagenomics RAST server – a public resource for the automatic phylogenetic and functional analysis of metagenomes. BMC Bioinformatics 9:386.

66. Pruitt KD, Tatusova T, Maglott DR. 2007. NCBI reference sequences (RefSeq): a curated non-redundant sequence database of genomes, transcripts and proteins. Nucleic Acids Research 35:D61–D65.

67. R Core Team. 2021. R: A language and environment for statistical computing (4.0.3). R Foundation for Statistical Computing, Vienna, Austria. https://www.R-project.org/.

68. RStudio Team. 2021. RStudio: Integrated Development Environment for R (1.4.1103). RStudio, PBC.http://www.rstudio.com/.

69. Wickham H. 2016. ggplot2: Elegant Graphics for Data Analysis. Springer-Verlag, New York. https://ggplot2.tidyverse.org.

70. McMurdie PJ, Holmes S. 2014. Waste Not, Want Not: Why Rarefying Microbiome Data Is Inadmissible. PLoS Comput Biol 10:e1003531.

71. McMurdie PJ, Holmes S. 2013. phyloseq: An R Package for Reproducible Interactive Analysis and Graphics of Microbiome Census Data. PLoS ONE 8:e61217.

72. Pauvert C. psadd: Additions to phyloseq package for microbiome analysis. https://github.com/cpauvert/psadd.

73. Kandlikar GS, Gold ZJ, Cowen MC, Meyer RS, Freise AC, Kraft NJB, Moberg-Parker J, Sprague J, Kushner DJ, Curd EE. 2018. ranacapa: An R package and Shiny web app to explore environmental DNA data with exploratory statistics and interactive visualizations. F1000Res 7:1734.

74. Anderson MJ. 2017. Permutational Multivariate Analysis of Variance (PERMANOVA). Wiley StatsRef: Statistics Reference Online 1–15.

75. Oksanen J, Blanchet FG, Friendly M, Kindt R, Legendre P, McGlinn D, Minchin PR, O’Hara RB, Simpson GL, Solymos P, Stevens MHH, Eduard S, Wagner H. 2019. vegan: Community Ecology Package. (R package version 2.5-6.). https://CRAN.R-project.org/package=vegan.

76. Simpson G, R Core Team, Bates D, Oksanen J. 2021. permute (0.9-7). https://CRAN.R-project.org/package=permute.

