## Supplementary material for "Short term tomato consumption alters the pig gut microbiome towards a more favorable profile": Kronas plots: per_category.html

Javascript must be enabled to view this page.

magnitude
magnitudeUnassigned

Control\_D0
Control\_D14
Control\_D7
Tomato\_D0
Tomato\_D14
Tomato\_D7

372205843686531126404228252299351029158814069698

5603887791217343520

5603887791217343520

5603887791217343520

5603887791217343520

5603887791217343520

5603887791217343520

369851673664992926234770250560881022018613959886

1645516987125481212857796923

1645516987125481212857796923

1645516987125481212857796923

650570885120476724692854

458750333649338318252046

1918205514711384644808

995098997428736133104069

495150263709362815862014

11671109827871384482

740773657674321370

9451031756727319385

2147196014791461700818

1870850143926713182091111757443344726137

1870850143926713182091111757443344726137

23901822462819853723323678973115624

2206185814091546655910

2206185814091546655910

925887226942728631074238

2212202917761769756941

12041278957974396573

1625154411921313545790

421738713017323014101934

10691140979928349489

10691140979928349489

511533457471214292

511533457471214292

22462030175119859291262

1017913768884428585

122911179831101501677

2052192916161646729939

2052192916161646729939

1577714536123721211665608016

1577714536123721211665608016

351329632352245310641528

351329632352245310641528

447103525528168543491250721404

28312223411451040873780714417

889873777493682730284145

750055376165664916722842

1925518127146821492567769371

1925518127146821492567769371

764170236151600435034015

764170236151600435034015

472041543211344513521843

472041543211344513521843

893288277504738531474266

627362655202504823043020

26592562230223378431246

323330012437245211881573

323330012437245211881573

688157364880543026343442

441237823132358117132221

21221723151215987811043

347231236251140178

28802323216918918921227

28802323216918918921227

1794167614151487651905

1488134011541189468710

306336261298183195

567250234107446326843749

372833312657279819352633

19441692145016657491116

28422276197219927961230

28422276197219927961230

1600315182127511350653288836

899481646882730328984567

17351701145815315431317

264126322430247710341554

1346155210381162474709

128711339431033379689

1345116710911214438577

1345116710911214438577

2116197415891573641987

2116197415891573641987

1588158412491293557794

1588158412491293557794

31022694233923938821389

31022694233923938821389

357531692724304712851737

357531692724304712851737

22752119153517608491185

22752119153517608491185

32532586217126889291429

32532586217126889291429

1809180915221425642896

1809180915221425642896

552853054085429918152548

376935602776294212101734

1759174513091357605814

517145883428367414862022

517145883428367414862022

568153724526460120222914

22602117178817498111121

342132552738285212111793

27152518201922627741167

27152518201922627741167

496443393370166215

496443393370166215

963090767192720833744509

963090767192720833744509

29539379104534949669804813720

29539379104534949669804813720

278329142336214910471297

278329142336214910471297

278329142336214910471297

12321121934963540738

12321121934963540738

12321121934963540738

713770785368508422103027

713770785368508422103027

713770785368508422103027

807409501828617254285050149912287369

807409501828617254285050149912287369

40028281053035216545803414711

575854014886277012292133

754613462424576566263069139227267950

701058985450266614222575

812491701025493176584646210291317603

812491701025493176584646210291317603

16317211662994091732773624654703

173373185582953652375156452996986

228214165644136444935384588673879

942769366863345724032471435387

11843610745980488816263007843602

35020320432344326287883813046

780673604629371479

780673604629371479

780673604629371479

508314663035059345881436619632

508314663035059345881436619632

508314663035059345881436619632

508314663035059345881436619632

608861504658423618992426

2385221368159051576465549111

788272735675538922613055

10929100927353779930794320

2080174714681400573720

1141096939794853730634857

1141096939794853730634857

1728138612591246465721

1728138612591246465721

1728138612591246465721

968283078535729125984136

968283078535729125984136

968283078535729125984136

925609280568619715524234948600

446349213473322016331937

446349213473322016331937

446349213473322016331937

446349213473322016331937

880978788465146683324071646663

729572815506599436484312

729572815506599436484312

729572815506599436484312

13791135219759945447885912

13791135219759945447885912

504148083594349617562284

465530326308230208

734572885205504324913016

25124118419496125

689654450413215279

784373345579518027003157

784373345579518027003157

965876634645350393

816860556596307344

404238972985270113061658

953636610529336336

10671065794709401426

28532983199620128921188

28532983199620128921188

28532983199620128921188

563155676542306456922868832094

563155676542306456922868832094

29073221221921569871373

1517715491111311065855176631

461549253442336414931927

13159171126218624722302

296394256229149127

438043933038299614071755

255962520219572228091591417084

2029222215221294749895

515384719038463376521387920284

515384719038463376521387920284

515384719038463376521387920284

515384719038463376521387920284

767872875709571522152999

1951317639150371387849997760

11158103847868834430824304

1812168813631542512819

713963145162507219142629

26322638233519747711222

160612409891127386551

13943134131319110139848911763

13943134131319110139848911763

13943134131319110139848911763

13943134131319110139848911763

11571037886828366455

747573968473577766469099

531149803832353414772209

1392214153103681022644005778

1392214153103681022644005778

1392214153103681022644005778

1392214153103681022644005778

645068844898491620602777

411943183166290313501695

33532951230424079901306

736747538755283551932288031206

32449341892326124803982712827

32449341892326124803982712827

32449341892326124803982712827

29143311162105322486884911395

33063073220823179781432

1786150713891554520787

1786150713891554520787

1786150713891554520787

1786150713891554520787

962397478082712536004623

30603025243922739461346

30603025243922739461346

30603025243922739461346

656367225643485226543277

656367225643485226543277

656367225643485226543277

29816299442255121711893312969

499250613800376517342636

499250613800376517342636

499250613800376517342636

24824248831875117946719910333

2398624058180841733069199941

1543715403117241126643666528

854986556360606425533413

838825667616280392

838825667616280392

10840113378173807932664498

10840113378173807932664498

10840113378173807932664498

10840113378173807932664498

10840113378173807932664498

26892816224221038701084

26892816224221038701084

26892816224221038701084

26892816224221038701084

26892816224221038701084

528845062739160377911543521121

528845062739160377911543521121

528845062739160377911543521121

1548144213161225426735

1548144213161225426735

513364918537844365661500920386

2274221469164121595863978513

760871515409548721963313

726272905517510522202950

443341033321327513151759

929191727185674128813851

1558431481841132991098224534961036

1558431481841132991098224534961036

1558431481841132991098224534961036

1558431481841132991098224534961036

2100120997158951584058288153

2529123525175881708872969408

804077926529541225013355

787547185057013557712267631340

2275724020162741571170488780

20862095158816609091091

20862095158816609091091

20862095158816609091091

20862095158816609091091

20862095158816609091091

1708214881128341295959517820

1708214881128341295959517820

1708214881128341295959517820

1708214881128341295959517820

319928082330243210871363

22171782165416528351044

1028790740800373562

652655174667458619772647

411239843443348916792204

28938255061859119823764610780

154211300194481023438305528

154211300194481023438305528

154211300194481023438305528

154211300194481023438305528

643660084375433618292496

643660084375433618292496

643660084375433618292496

400537072738280811371582

2431230116371528692914

708164974768525319872756

708164974768525319872756

708164974768525319872756

708164974768525319872756

1442163679950468118232538

1442163679950468118232538

1442163679950468118232538

112153314738320287791099

1646161713061245508707

956916976077783271392

2180211617211837702955

1847179114321516578779

333325289321124176

1026937846816342484

1026937846816342484

26273246931918718297728410655

26273246931918718297728410655

140581295710051953737935637

140581295710051953737935637

567055524260400516832307

1780171512961196473743

660856904495433616372587

12215117369136876034915018

2087193516701592629850

2087193516701592629850

1012898017466716828624168

1012898017466716828624168

1348191211221031721025083778153940

1348191211221031721025083778153940

1348191211221031721025083778153940

584358474586452518332591

584358474586452518332591

529464996536397373611494420244

529464996536397373611494420244

760306531062189606222100431105

416237393305390811301753

393293544235451312651232317847

32539261292343325449755111505

328281233267115185

328281233267115185

328281233267115185

328281233267115185

328281233267115185

1868441521024429145302371268984055392387250737

12842354122713288804297940799637079344963305

378303873628781263001144214916

28174287202156019558847311176

1954520318151411369359007900

862984026419586525733276

9656100167221674229693740

9656100167221674229693740

1569715806116151135043826209

1569715806116151135043826209

1569715806116151135043826209

12432168118582658490165911486735855304795394

512375278438170377411558120567

11159115518371767033264248

481451764288398715922243

352643605725511260841066314076

32900230124922816323710492363130361

647659804912499317522823

1970071775791367841423045530778983

12551911769086467898073530448555

1740052174822012294431171231474828625903

5328910744939816399653075926146

30736354412320921888930512114

29699283952260322031823312447

1818222517441287572907

31640131154821807323111493676121625

430714114231539307931212217586

491363555246350259346230140529169572

36913426606925850921072576836125285

1753811634301194711244604385063821

2291602372751642201427385894676400

460940041961212986086359853713067021796619

1990281909091426541449275304881364

1490402160383110753681410898500996728215

34244523866121878021911267503122122

2252221340171781683967269914

667669642926447401357716160065251680

1791492141031710124031380223493534562751

958428813772302688222483040573

444774813732733292151338216656

444774813732733292151338216656

32232732876623924521899694328124816

1524221500301126291043154309059785

2501126164182101597277399315

138981441910458976641625484

928049706668980629862790235782

381924108728968259571143514450

939097317583704727174025

939097317583704727174025

1851722200063213190971383940606034722472

1819207196229612911321359569596274707554

1730516085130361322548457385

1521022251149291114649157533

31448318492416821516895912430

31448318492416821516895912430

33304373030855229974723263549366771293672

31859162895192219541822189918963761235755

1445211356631043291073634030157917

913558944568800666182753638275

1755616963141221384553507692

439404318430915305501240216963

25682254692046118995854711664

417738293302322812371956

2132120476169301656864239598

1433413869115391103744086473

698766075391553120153125

356659358521273736255479106580146786

366333632547252910911356

366333632547252910911356

2107122141021630081495226334085758

390434012029231258521195814756

662026550451080477191938827650

783283046323540924153091

613156158747400448481855225476

363203858728974256941102714785

1422841410561081811034284214959672

923219063570196679092705839242

2066020053156161477659038494

29303303682236920743918811936

422434373310281206288353172368154848

422434373310281206288353172368154848

422434373310281206288353172368154848

1889391655941182211413625862162141

13784012893595798959133791351803

14357122639450891646196039

28042739219518758711213

857883076229570027393763

555583567539367233686257224476

143581979799461121950335413

149942318127591189934728781554802509039

149942318127591189934728781554802509039

302998342395254098180441112219110656

302998342395254098180441112219110656

11964251470364935836548340442583398383

2055513026391827821398576096872810

29156321642100120958826110741

553959600042410837151512202870162192

2085362912461435291312817910464847

143831172540112092850894683650358

553927173365595196434454437435

392020465670324254800226471011041341623545

33191315967850380551618495799069261367469

693784345920482223533065

439056773996313615182056

25472757192416868351009

3241903809982764676194776396371217514

3018193768109745126180166387280207212

22371418731955014610909110302

2555631601209871615571039765

150824161390984482649

1580020050131291027445075993

824891356468489721143123

27832231942928289771215273874501321043724

1713118574131521063844556549

27660921924354288456015167494456771037175

510724875834942313021406418788

1613142513081046443681

407134058829736282091232816526

874667453898204712931581

12815312614781279751373690374613

12790012590381059750323680574538

2532442201059875

601073599182449284415131197208256076

479434878839337324672584430485

435004404435930295432421628472

444347443407292416282013

2157322202159491481568218737

1975620271144601345661537889

1817193114891359668848

2369824652183111693872319815

10136108517973709331624094

135621380110338984540695721

364464362511268793252334115431148518

131051360710045898646855507

645986396147729432961980325642

11859113428460824335754757

25955125836919107118120582466106544

1535115232114881060449026068

446764517133206295721351718462

446764517133206295721351718462

987199585873688690052836440059

832567988061892580592368633749

1546315978117961094646786310

750877226231615826693452

493851134298411418312456

493851134298411418312456

493851134298411418312456

493851134298411418312456

2570260919332044838996

2570260919332044838996

2570260919332044838996

2570260919332044838996

556957374288431017742411

556957374288431017742411

556957374288431017742411

556957374288431017742411

1626820714

350836572768268711101480

2045205415121603657917

119294610826379437951159411420436588541

581753144022403917292338

581753144022403917292338

581753144022403917292338

581753144022403917292338

1657791524061184621476216586277047

724568695130585626933321

724568695130585626933321

483745153369397718312257

24082354176118798621064

308530732376270710241358

308530732376270710241358

308530732376270710241358

556352804196497721002500

19315617218488107

19315617218488107

537051244024479320122393

25342518186521699311074

126811279191099421536

1568147912401525660783

1591613017103661578353176016

1591613017103661578353176016

1879163913641958714777

367032262620327512091544

579743173430698419832000

457038352952356614111695

10877110119277630969094592252739

130111309399981158355866803

253424091978233311401388

1634183413031763808931

278524101797225910661260

605864404920522825723224

136971165485041451752555835

947984729964451491

269521801793240414291536

245020521699231512771428

760564384283883420982380

441514096631321362631574119003

597257224081522622692596

238521551751244412481470

26211240671867620400858410738

651542443761276337

893284806370743233643862

277392551220381254961546216507

229422472366219512411234

573350674028529435813722

374133852675345619462241

1464132710141430820869

309726752129252312901552

922839640890509627

873780996266805949025113

175118731263164911731149

1017399677426905038784591

22841998163121649311090

788979695795688629473501

304827982136270110091273

304827982136270110091273

1412142710581374482651

1636137110781327527622

264922571981254410101087

264922571981254410101087

24482141182523389091000

20111615620610187

1950217920146471614467878753

1950217920146471614467878753

1683177412601641633751

772685584768313363

818845693795334453

529566404463205232

405405307355179196

1093596588384872537915094

436039873015339713321664

117421997261390651338104692554763

109008917891320471266404406651176

497623932685293777602358724573

463143705636577919621907

401422999675430677581992020457

354534642981309112241563

1444149612461132481646

587565196846365485332033226389

506074433039786418741781822971

557452434411465517082283

25752395216820048061135

490495389347147214

490495389347147214

1964190315761675733884

1964190315761675733884

1539153313101317575690

425370266358158194

644960345442549521262703

644960345442549521262703

332132642992293211761437

31282770245025639501266

666710525550228328

666710525550228328

666710525550228328

666710525550228328

480031421855370098541831175266275640

20251147771701816799849811373

20251147771701816799849811373

696846905656581329623942

1328310087113621098655367431

319032772522234112081449

319032772522234112081449

808731613581290323

23822546190917609181126

1912717876143511506264778724

1040197597852830634334698

358831632748292413111671

1536132710551137418605

22122191175119217281040

30653078229823249761382

751469215486569725563391

539050243871403518712430

2124189716151662685961

1212119610131059488635

1212119610131059488635

2109419058161981640471939668

438539553614355217142283

280025532393224310621523

1585140212211309652760

444741843509344414921942

444741843509344414921942

700762345265551723723263

651522515588265331

650451474559237378

570652614276437018702554

525546853810389116152180

149913109721121470635

375633752838277011451545

1048392217515804435284661

339431242548271311501492

29702783220123309591228

424341347383191264

804715589705383447

804715589705383447

628553824378462619952722

628553824378462619952722

795171275999586026573515

593952344310422118982522

593952344310422118982522

2012189316891639759993

1895179515361513705883

1179815312654110

524224892238970412851768524089

373413446327412289861261016630

297632804222385236961038913500

535045663474370715012149

2228185515531583720981

1508114459115581229950757459

537249843912412216782420

554507403461230586

877822734687329473

827881466509702928383980

456044223653371517252315

456044223653371517252315

388737963075314614211896

673626578569304419

1530913486111531191650226696

1530913486111531191650226696

978184526745732829263977

300825822310243812171565

25202452209821508791154

812556786260257632482766437741

300321412085233610011442

2238170214371729623979

765439648607378463

119776410081124545732

119776410081124545732

413536693823348918572395

413536693823348918572395

339962885925405265481141715366

339962885925405265481141715366

996980127048769134304797

996980127048769134304797

641155785106504324553215

641155785106504324553215

1705114183114121232048677012

573549644333446318742427

27082042221921579311358

860871774860570020623227

325730282608285112181614

325730282608285112181614

22361628176218468741168

22361628176218468741168

613975390348011445962185627542

613975390348011445962185627542

472838694282360020002303

720964035333545624812979

329732102617252512081517

401933513343304015131959

2162119969160401529373379211

627154085172441722782996

1425211693112241026550396577

26922452187219248661203

26922452187219248661203

26922452187219248661203

1004688337695790835664789

1004688337695790835664789

22262066169016426831038

614452084735492523002993

1429133810451151496636

24722122519087122

1251551113039898626711950666109296

1251551113039898626711950666109296

473481454381176257

600753524253621820103160

19891651171917938091083

475238233540367516452355

20517018917986112

3847387035012075414676568

9733844472731366536586565

1297100111191038505743

544435480445234270

313828362382284310041587

1628140214141223562804

655652542642

719066515306552024073077

502247553742378716632240

488545133711370417512331

729467125236825232204167

1563142513101375602930

542847123994626116972913

399833003123331713512061

542250504345500618682571

3078526746268601609461722155868

425935623056371913671998

269193264269108205

11522110068751981139895712

384031572912288412401677

450993933635898356101665522579

33929223927997170

33929223927997170

447603904435659353311655822409

968687839930465580

654755415052512623232962

1018687398029801937935179

27059240772173921256997713688

25202224270718649220277572372101770

32643234247524119661389

32643234247524119661389

32643234247524119661389

29302303452234223440951113239

312034512482283110301582

312034512482283110301582

408741513003310313301744

408741513003310313301744

1939020009146471497262168481

627464704579514720512848

131161353910068982541655633

120412579751128384655

120412579751128384655

1501147712351406551777

1501147712351406551777

826038319560581592392467332294

514555157637979368521538820495

514555157637979368521538820495

2571425978184561839875999588

2571425978184561839875999588

543456414146398916862211

543456414146398916862211

31793147228522809081241

31793147228522809081241

31793147228522809081241

1829117660139181365953907475

887384876671656725393628

887384876671656725393628

941891737247709228513847

941891737247709228513847

26282409190921017331038

26282409190921017331038

26282409190921017331038

30028292092258322511874412094

1881518682143591416455257673

562355684135410315872123

664066435042510820662779

655264715182495318722771

11213105278224834732194421

27692645205922058011171

844478826165614224183250

827277350860399771342144733000

571655274218429116082251

571655274218429116082251

382241562981311612291690

849825672692273404

29733331230924249561286

731896382553200697271861029059

457139563465370514522265

686185986949735660221715826794

1712101599191251311287855805476655

10371038840800351459

10371038840800351459

10371038840800351459

1455413810109431099453746791

313929392293232510561454

313929392293232510561454

11415108718650866943185337

257525732006200010891380

191177182167122107

22062117168717288361028

315127342217222610361325

11791062927953528661

2113220816311595707836

771447139056133571292612634150

22352185162016658811211

125013189801011527740

985867640654354471

433394402486401437

433394402486401437

2139920104157981605177279492

734065615105548326383207

965593327336715235824384

1421721591507995

888853717691329402

337431862481257510991404

606059334561464821102835

606059334561464821102835

2041518729145881457965088582

630757294597451120182665

828377035772582626523441

559750924011405017642361

22820520819274115

25162027181618597021023

25162027181618597021023

551550693907418016632437

26832440174819597641075

28322629215922218991362

974990707112719831244302

26332636203120159081233

363731402705268911841576

31812982213722258991338

298312239269133155

530949813983408319302369

11781017905872401563

22392210171016908281003

1892175413681521701803

158212849821018438619

12571002748764345472

32528223425493147

1931161413641362642843

1053863732777327440

878751632585315403

12418111307872886334274881

12418111307872886334274881

135513499951095469627

11601136907975381622

713965014495498219812733

2764214414751811596899

342843205025925269921242916480

858666653690359487

858666653690359487

278672614621406221101036313731

595583452555231390

602552472455234330

2389227017401789753974

759724627561240362

366535042932291514141884

1691661121365490

129913389311101445621

505490455428195256

1437126610951146584704

276240196203116127

260266222204115139

793694506577295392

560352934326455223482940

1596132512001058494666

367933892749291613291761

2072011401545589

34431027224792173

481464464502180265

922860699699362445

481490421431179227

24420817220589123

10871094864906381546

1841621321286783

290257227242111144

555952383866419217072262

1655151110511220518705

842912702716270370

2038191913711546596806

1024896742710323381

148412001106976447617

148412001106976447617

929670679534265356

555530427442182261

2298222472168861743176469937

874187286520681130453960

23422509182818628351069

765826499596259327

551442410446294334

1966190914401548629821

1506129910521075459595

1611174312911284569814

1424113744103661062046015977

603056024243429718282441

2218207015731732714943

599360724550459120592593

730768295426560022543340

34762996247626379991516

751755586575251362

344390292306143203

402358293361110148

1979149313051395495803

29963012230522449611426

615597446458186266

23812415185917867751160

835821645719294398

10010775934250

735714570626252348

14322975122282778775888943352135332685014240

13861030118429868460274911795234150764849239

13861030118429868460274911795234150764849239

12902103649245930833154764

12902103649245930833154764

876821280726915506190564489823080253235751

876821280726915506190564489823080253235751

44038483255378252184130032669533921392995

44038483255378252184130032669533921392995

457803674337264290581204816190

457803674337264290581204816190

630288467810385734431422138296199539

40989630021224835427733788074125770

794576058849236547471788526320

14093510701088144993383233747449

1899815466143811331048156691

1899815466143811331048156691

1899815466143811331048156691

25232412218219738241016

1647513054121991133739915675

1075508937371366728122711737767

1075508937371366728122711737767

427132132964284811991612

427132132964284811991612

743667465476532321642960

743667465476532321642960

958437941462926646412375433195

33274257211965721647773511349

1510513925109771013239425293

518741083746350013551907

2098418032146121459551997527

2129317628139341476755237119

2290461926341578971578726062483394

2212831858781514171520455841480440

586647444399407416862236

362368251347115158

550443764148372715712078

2143591801151460361470805637277708

1708514030113251165942005943

559246274088384915462092

31616262412175521016796910796

1604413517110821110040145919

720864845001489619882784

631745188839875421101583722521

12099104758787839732634436

911783626464668325043344

657056244983478319002452

1134591777626776131314282

2638227619041945780966

531442913635359613491996

2558203317861670777904

1816915931132101341055457064

583051594515420515692209

10581019982891356496

10581019982891356496

776367566480582722102954

776367566480582722102954

776367566480582722102954

1063518781871970715752563637149

1063518781871970715752563637149

29618238551912319803699810109

29618238551912319803699810109

12858120689079893433884822

777171375408545420382939

508749313671348013501883

638755189543768428381525022218

824464385041550619272892

472683880932381315101121716482

836366486346582221062844

278722660421254186531008413321

278722660421254186531008413321

11749104638681747837575286

11749104638681747837575286

1847179015691308770959

990286737112617029874327

20822038153415117551044

20822038153415117551044

20822038153415117551044

140411410311039966455726991

140411410311039966455726991

12344124069769840149826180

1697169712701263590811

595877701937345443881506320469

595877701937345443881506320469

595877701937345443881506320469

595877701937345443881506320469

595877701937345443881506320469

429093407025769300451267616787

129712319621010519629

129712319621010519629

129712319621010519629

129712319621010519629

176621346799111210749736658

279124241832209210241205

279124241832209210241205

279124241832209210241205

148711104380791001539495453

148711104380791001539495453

2069147512741522654807

1280295686805849332954646

1605117910151255746898

1605117910151255746898

1605117910151255746898

1605117910151255746898

2234518193138811567364388602

2234518193138811567364388602

1653135310661319491719

1653135310661319491719

1644127411041445753942

1644127411041445753942

1904815566117111290951946941

140710759671313719853

1764114491107441159644756088

567854298135900371081994824428

1831681291383967

1831681291383967

1831681291383967

1831681291383967

1831681291383967

23311597193418919711219

23311597193418919711219

699444935896552562

699444935896552562

187119263287193169

512325672609359393

16321153999995419657

16321153999995419657

16321153999995419657

807776611471164275

807776611471164275

807776611471164275

884838521930

884838521930

719728573419145245

719728573419145245

339624803138261914081640

22691560194519059141111

37924117820994111

181121104944863

181121104944863

198120741154648

198120741154648

18901319176716968201000

18901319176716968201000

18901319176716968201000

11279201193714494529

11279201193714494529

435324272349315343

435324272349315343

692596921365179186

692596921365179186

1719014168108101165574449047

36073186224626189791442

141913579341113429653

141913579341113429653

24924616617083111

11701111768943346542

747590453552177258

747590453552177258

747590453552177258

11801078741794321473

724682452454200285

724682452454200285

25819216920076123

25819216920076123

1982041201404565

1982041201404565

2611611181595258

2611611181595258

2611611181595258

566447399463158287

566447399463158287

566447399463158287

20218116020866115

36426623925592172

1981611351194879

1981611351194879

1981611351194879

1981611351194879

643849353906409944064703

643849353906409944064703

2322151371627494

2322151371627494

632547386459160270

632547386459160270

376626302185225735733527

320153157185469403

913660513531628656

106266861463314701396

431329240279276303

418335290255316321

228144123118234201

217193144122130172

1771481041345075

501497349365190235

501497349365190235

13071046849856409577

24326217421594125

2051751481527198

28618217614569116

573427351344175238

466042453118317613661863

754607517495237314

382238250225114153

333199209184102128

493941411225

1149558823446

1149558823446

1111521091015055

1111521091015055

147122100873960

147122100873960

390636382601268111291549

390636382601268111291549

1541801201114488

151913119461028436546

12551219832860368558

532503369347157190

446425334335124167

595493391438190238

595493391438190238

164140861144471

164140861144471

431353305324146167

431353305324146167

1126701615742297435

1126701615742297435

1126701615742297435

1126701615742297435

314024481929205510611425

603484388403190287

603484388403190287

603484388403190287

603484388403190287

896761576565261390

896761576565261390

896761576565261390

896761576565261390

261220221234262296

261220221234262296

261220221234262296

261220221234262296

1380983744853348452

806150582436

806150582436

806150582436

1300922694795324416

252142921334075

252142921334075

2651851441878785

2651851441878785

345262208223100117

345262208223100117

43833325025297139

43833325025297139

2363316095130271338468318032

9311574753333

9311574753333

9311574753333

9311574753333

313625191414

313625191414

313625191414

313625191414

15929068491222359497

836501451673207294

836501451673207294

836501451673207294

756405398549152203

756405398549152203

756405398549152203

682457425483191276

682457425483191276

682457425483191276

682457425483191276

22827513616695117

22827513616695117

19466161717

19466161717

20922913015078100

20922913015078100

2061313817111901114959716908

2503181415661734729944

2503181415661734729944

2503181415661734729944

382305322312108192

382305322312108192

382305322312108192

90128731088793

90128731088793

90128731088793

42311927613

42311927613

42311927613

948069015190503533973561

948069015190503533973561

948069015190503533973561

4583198116821269584739

413615131326

413615131326

449310233202125118

449310233202125118

4093163514341054446595

4093163514341054446595

728595668669295340

728595668669295340

728595668669295340

1155712566791267405

1155712566791267405

1155712566791267405

1650135011041204498621

1650135011041204498621

713585413448203258

937765691756295363

394489328270168187

1942361461288599

1942361461288599

1942361461288599

2002531821428388

2002531821428388

1381741131004961

627969423427

565410304349166232

565410304349166232

565410304349166232

565410304349166232

565410304349166232

136910508921075457614

538460371463202274

538460371463202274

538460371463202274

538460371463202274

831590521612255340

374426281814

374426281814

374426281814

794546495584237326

794546495584237326

794546495584237326

417137893126347114071877

22772086185419467641000

22772086185419467641000

9741003757811385453

9741003757811385453

1303108310971135379547

1303108310971135379547

1846165912291481620855

1846165912291481620855

1846165912291481620855

591493371423190267

125511668581058430588

484443442322

16171320125

16171320125

16171320125

322730241117

322730241117

322730241117

23960262691555416983779214389

23960262691555416983779214389

23960262691555416983779214389

268526962244257510981483

231735312838

231735312838

1467764632246

1467764632246

23972504201523856191076

23972504201523856191076

1199813096429323

1199813096429323

21275235731331014408669412906

15025181929059860348516039

14321722615

848325373016

14111174558233780030004506

81662278474418151502

421730142527414112475960

525245371926

771783143211490

733383377503236242

7412367681746

27812163167218887834979

500276283213181177

1077728532948318387

33495801817577

774939662727

157127621184755

784555772034

426675801615

772741973257

28926414526783114

23553562188

95616391192716278520

95616391192716278520

1541121457441172251185394331970475

465436541925

465436541925

465436541925

465436541925

465436541925

699068275404518119892962

699068275404518119892962

2322220718011705608963

1922179114691411511778

397403305290124155

3462391852176188

24020416119269109

526524481385135234

28727423223291138

126147105953154

40041633229497185

40041633229497185

24552544210419237541108

848894733643243358

848894733643243358

1607165013711280511750

1741771431245776

9791050803776321464

757666862246

379347359294111164

1391511081174266

1391511081174266

1391511081174266

2074192513911436585825

2074192513911436585825

1610147810761131479650

464447315305106175

736647499519171253

736647499519171253

736647499519171253

736647499519171253

736647499519171253

637622520475198290

637622520475198290

21220719017568120

21220719017568120

21220719017568120

425415330300130170

425415330300130170

425415330300130170

1457031375941107661123104094266945

11971085954913277524

11971085954913277524

11971085954913277524

11971085954913277524

399939223543335610211709

399939223543335610211709

890907810783220389

890907810783220389

2141213219081858571922

2141213219081858571922

968883825715230398

968883825715230398

749170216025592117793054

749170216025592117793054

749170216025592117793054

35113312300829287581432

398037093017299310211622

498952623852394413262030

498952623852394413262030

498952623852394413262030

412943583178334711121689

860904674597214341

758875165836567820163026

758875165836567820163026

758875165836567820163026

708615512490155251

854843632678232352

472421332321112184

1059921758778283422

652697519513170263

385448292300119171

740723631527191322

495495370375113169

580652490481203239

467434378343120196

722745529499195286

454522393373123171

1829018340138581373352277463

1829018340138581373352277463

13150133409647965038405260

27124523426369126

12879130959413938737715134

514050004211408313872203

514050004211408313872203

10796104588517784629444295

10796104588517784629444295

10796104588517784629444295

511049353948358714262064

568655234569425915182231

331982783424945276551004121264

331982783424945276551004121264

703658621597199315

703658621597199315

32495271762432427058984220949

24017195871786920316745816550

435637493600369211542535

412238402855305012301864

581555615643236432641631123580

29031281452160621412796711644

413539343012297111021558

413539343012297111021558

12985127029659959535585136

12985127029659959535585136

770270345604561320093018

36022938260523338431325

410040962999328011661693

420944753331323312981932

1738186512651320548737

24712610206619137501195

27332262682018620518785611119

31383013241525327901182

31383013241525327901182

2419423255177711798670669937

1730916532126341268551397123

11711133912971320474

404841443040305411521701

1666144611851276455639

1792174314441334488817

1792174314441334488817

1792174314441334488817
