## Supplementary material for "Short term tomato consumption alters the pig gut microbiome towards a more favorable profile": Kronas plots: per_sample.html

Javascript must be enabled to view this page.

magnitude
magnitudeUnassigned

Control\_D0\_P1
Control\_D0\_P10
Control\_D0\_P2
Control\_D0\_P3
Control\_D0\_P4
Control\_D0\_P5
Control\_D0\_P6
Control\_D0\_P7
Control\_D0\_P8
Control\_D0\_P9
Control\_D14\_P1
Control\_D14\_P10
Control\_D14\_P2
Control\_D14\_P3
Control\_D14\_P4
Control\_D14\_P5
Control\_D14\_P6
Control\_D14\_P7
Control\_D14\_P8
Control\_D14\_P9
Control\_D7\_P1
Control\_D7\_P10
Control\_D7\_P2
Control\_D7\_P3
Control\_D7\_P4
Control\_D7\_P5
Control\_D7\_P6
Control\_D7\_P7
Control\_D7\_P8
Control\_D7\_P9
Tomato\_D0\_P11
Tomato\_D0\_P12
Tomato\_D0\_P13
Tomato\_D0\_P14
Tomato\_D0\_P15
Tomato\_D0\_P16
Tomato\_D0\_P17
Tomato\_D0\_P18
Tomato\_D0\_P19
Tomato\_D0\_P20
Tomato\_D14\_P11
Tomato\_D14\_P12
Tomato\_D14\_P13
Tomato\_D14\_P14
Tomato\_D14\_P15
Tomato\_D14\_P16
Tomato\_D14\_P17
Tomato\_D14\_P18
Tomato\_D14\_P19
Tomato\_D14\_P20
Tomato\_D7\_P11
Tomato\_D7\_P12
Tomato\_D7\_P13
Tomato\_D7\_P14
Tomato\_D7\_P15
Tomato\_D7\_P16
Tomato\_D7\_P17
Tomato\_D7\_P18
Tomato\_D7\_P19
Tomato\_D7\_P20

48508974073418365254045331983320996433024225712523600592420006120873883252962312966143527283860981381372336875733880576319059736866684009842289088620058193040318415991618675643071387328609316526413409991101961324124702766051285977592550121923352567847243469338264212812740243210276889713518096688541004733153010179775611772981140296109406575777937303717278491208247120154874946321866761587603178269715102171742361

149199387163352153363920646251404813334850485841610131717105453019426227300179135101436180214134729799911960115236040591223

149199387163352153363920646251404813334850485841610131717105453019426227300179135101436180214134729799911960115236040591223

149199387163352153363920646251404813334850485841610131717105453019426227300179135101436180214134729799911960115236040591223

149199387163352153363920646251404813334850485841610131717105453019426227300179135101436180214134729799911960115236040591223

149199387163352153363920646251404813334850485841610131717105453019426227300179135101436180214134729799911960115236040591223

149199387163352153363920646251404813334850485841610131717105453019426227300179135101436180214134729799911960115236040591223

27062505177227442219239311184132257018011743379155901831190319041506235829732670166285532602444108714341101167715524823100192514376611464127310892529125822476461022662778916675968618112238550415121217992987124649791251981720

27062505177227442219239311184132257018011743379155901831190319041506235829732670166285532602444108714341101167715524823100192514376611464127310892529125822476461022662778916675968618112238550415121217992987124649791251981720

27062505177227442219239311184132257018011743379155901831190319041506235829732670166285532602444108714341101167715524823100192514376611464127310892529125822476461022662778916675968618112238550415121217992987124649791251981720

32323126131938834915715034416325817026919616438926553018626927413337337816221816820224888254293195112342143110493167466135131106120105112147848573104270125116130162130200125121

10168103922611141081016781045728318549266359715103741235132321225242271624

10168103922611141081016781045728318549266359715103741235132321225242271624

146111012101781813411115101489206101315151071615101991014771310610107343343136715932212926302624352120922425

146111012101781813411115101489206101315151071615101991014771310610107343343136715932212926302624352120922425

2242423211612121121287241264272424441361243392242565

2242423211612121121287241264272424441361243392242565

2972072382973333151311293111392461432411821453632525151572602541093403581412001501772325423427017297328120934801514405579708361378451603973226958891135101879387

2972072382973333151311293111392461432411821453632525151572602541093403581412001501772325423427017297328120934801514405579708361378451603973226958891135101879387

238322741511242518312044961398222261638148536215321163517391515124118282787240113887222887206692512169331475130439428461632124254911221130979203610911781511891556658811563821534103731240012421092876857108448491051856599

1791191086917361155148859930711184122210402742452312081263106284512932268194810014741614133667680365612859212931288120575530063786047912606701149372574420493553382602395846214323865769539585688529677643421

11334235413513612127122312321371122224421

5215155494115511813631114393742242111312343424363352161131722

1718184180316331067138349929721069112697326694434114511979817681220219118779123791526125359271858012238502001148112069521055680039511876231066176384290285343173340242660107161769600395393591363413514307

68486087809887951048959577749577365566861849486697981745867921367258887653796842721921831272052052052621521781061629516714119093161255127111

81398873699265151171127774832236312310964231173938529603205391023821482215374728067524350794621502526252040203714121358359496430413314

81398873699265151171127774832236312310964231173938529603205391023821482215374728067524350794621502526252040203714121358359496430413314

4372874135465193952584107382142903114223002772992682642833002441815164361872712071312856912572663451363261733935952633878822910012116811513510112664521751962542022984230278144131

2951812703322992871392445871472231953232141912262061632022201638134228213517315887203483341702017524311895270164218611267270935792778748241261268314314140531058989

23977351641373321676856112328112322513532422321313263722

3671111213571116116211180611391125824027411311121121126249610812057

521830491402967554218162823333927353015303911546418391415254172823322311212314219641635121011888818351517201592317

11279811646101112207142042564421713612115656696397172362222776282561

77839482716042101853836604436302322405339292181662642312743118449731751361850715413346135443271127816192518133345371917

74381417088693935013375788515364764564406360583815489234032385010148877033924564131791993038111865464418282013674424213460553633

1015555126108244811532833114133131852144115146621113611142262

4113164339195372131343412053212212445143121332835323712652357681524

18545815164221410762912774627458206262562209157163865294741012648433713611

1613532116211193331218347019311172132211734924669817655421911301418272328994416121051135141236102219158

86211122911023642244291468201610281421431113365925141111223413

41116202819935392273420662152426731831813722347328444132581455613012578545

94222221033571253344421424412545313624113361212141

5615125131564112592591191426737265232142543213346152542592

16582107151740018090197741765496981382120260101181371214231199471354315745168631395811258124911399613050586024165175168766140661039764281402129561464493481570550511591998766358142171208115340306767754599348748793991444239325107304017047711115748706747194886396529671205009

852672275233114474585384244564241841579214211814628112

852672275233114474585384244564241841579214211814628112

852672275233114474585384244564241841579214211814628112

852672275233114474585384244564241841579214211814628112

852672275233114474585384244564241841579214211814628112

157501006416451171241869816747907213075191769546129441344718843127901491715804131751063511831132081234255362285616457828213289991360731319928191395289021485547991503492836041134511145414539290463874394332346143757417137224801286916197305111038299717989576052497067504711

3414226432934787528736531846266440531937238524544381270833502741280618322586259124471237719533001836280919841124237364049062087282510193828193514112628322637906151283185074611628858318261174669556135652323361353517121563102717351187

3414226432934787528736531846266440531937238524544381270833502741280618322586259124471237719533001836280919841124237364049062087282510193828193514112628322637906151283185074611628858318261174669556135652323361353517121563102717351187

63276668116914878984876701015475864551485268141388759814738692059308533854213746111593015181126191639168305158264825183516

63276668116914878984876701015475864551485268141388759814738692059308533854213746111593015181126191639168305158264825183516

335122373227471951713562179825863955188923092384428026543275265527611781253825392379122370573213177727281937108623046204847205727409863743189313982554316536756061253183572811518598128101135653548132651813303350916641538100917001171

420247487606819412207355539264349362648343428443350259272295346142123838627539725816333065652160471132563241126374424549661562088195135881111367853184613361420242191147217107

475324460390433511268366614281333343511380371402499286370345411165507436191339270140307893102604141323462691853683454219018710111217510610510414110942194281196170250192171184184

2456166622803723391926391323186528021344162716793121193124761810191212361896189916229165312239113111992140978316674663885163718557222834138310871812239627054509101526535881618619595858466453948428727462919117211556911299880

8324338447139818194098721048540633776922556815938557566597661794182110382142183245439688313962631396424394052719581752477211825215214415819920213728413388380328389216560308286323176

8324338447139818194098721048540633776922556815938557566597661794182110382142183245439688313962631396424394052719581752477211825215214415819920213728413388380328389216560308286323176

8324338447139818194098721048540633776922556815938557566597661794182110382142183245439688313962631396424394052719581752477211825215214415819920213728413388380328389216560308286323176

4712814944235044042003815502723863594823114224333403103493173411336134452013872651803836933522442013242725713239328039382156869211410998731476445202199194122271161141159128

361152350290477415209491498268247417440245393505217256248344453494903762204451892165007029189544111513270634242443793696665244901046413769431781291959428914714516448

40821543236650551325145552333134643249231746957333432826137043811555952426141333324358374304154532131573297105570290400651297574831091151071947041200166195123318164158215129

40821543236650551325145552333134643249231746957333432826137043811555952426141333324358374304154532131573297105570290400651297574831091151071947041200166195123318164158215129

102729885951466511411477748510559114116957085807932152139619472511281760379628149572112861781638151620223424232212504544387240314026

102729885951466511411477748510559114116957085807932152139619472511281760379628149572112861781638151620223424232212504544387240314026

19011425318227925313925030018118421728217526032218218512919624461281268144217202138318351761002766928716865324171222347040414954516513730211028210159168838812989

19011425318227925313925030018118421728217526032218218512919624461281268144217202138318351761002766928716865324171222347040414954516513730211028210159168838812989

116298199131114479110973881301058395135577347941152212611756102595413722681716034137721911858100152120171433301834188483950267841394614

116298199131114479110973881301058395135577347941152212611756102595413722681716034137721911858100152120171433301834188483950267841394614

7965668968629108675426141002533785768993667809919674585544772605312103998447670950334170516260851180225473946032369054474714835116215623918522519623012476375403309239451326277309261

7965668968629108675426141002533785768993667809919674585544772605312103998447670950334170516260851180225473946032369054474714835116215623918522519623012476375403309239451326277309261

7965668968629108675426141002533785768993667809919674585544772605312103998447670950334170516260851180225473946032369054474714835116215623918522519623012476375403309239451326277309261

845388799180375496465373795757826158504540268497465548326321453971267554245054522128615141117728817292328233832202813

45534863496130395034413845494953412930462419476843402823291129296111293910453236719106181210101373243122141620141723

793489696284414599506486666893906269455442121019142755429621161438615653831435279192917241813131511113464021213736142619

918011491761088161945876739475961047658851065942103100446964568015728467269339749751751949181123242525271110474335295437393127

412436285638273661385837604455534328294136185051163527153311423333173814123433449297713131481544131524152927142010

675011074637749791094410168103786984595752747435112634851452059224251723080382960475013342117241726171885324429194426262833

55296765736150499041767796466660644637845519877936823130581349337315534325684775142818181918302720119253126283424202319

5251713665443134733851695651646132353541422253712655261950632406115462113393474112267813111310125192213112718191817

725075103957557629061837310541748558734091575398935780583786126953572368432862537192813172718182621144434532255830223726

452752477152283766294942112354958394338573616101652739281754105615541843321651385032058191016181595181614142522212016

1038497154155135738711457100811158697124876972908131138127678665439822816611742966345997792114624194528312337195556342235838543747

6231495354523831603733516237406552203143591965792442292033830255016533616422649121917711814715106243023173116142411

55226160651961857326556664838147450476048752964558740942344452816169054126851637120548983406287579171554356159549365518892231081061471291341101839751249272252145352176183221129

55226160651961857326556664838147450476048752964558740942344452816169054126851637120548983406287579171554356159549365518892231081061471291341101839751249272252145352176183221129

55226160651961857326556664838147450476048752964558740942344452816169054126851637120548983406287579171554356159549365518892231081061471291341101839751249272252145352176183221129

793593112118100381051087295971158084101126736568931711475587882421011466408525100552087407915301718242428193098494935166829353517

47322651340750047322746154030937940764540744554446133635837643514457646621043828916338869340247494146454301139462325439741939188123105106911538843200223217129284147148186112

1240596140712411257122675297714406609501098142995010991301933823869100697738017881311692964756414103220392358311864001055614414954684103323952224323930429630625134320180588539459392686425350455321

1240596140712411257122675297714406609501098142995010991301933823869100697738017881311692964756414103220392358311864001055614414954684103323952224323930429630625134320180588539459392686425350455321

1240596140712411257122675297714406609501098142995010991301933823869100697738017881311692964756414103220392358311864001055614414954684103323952224323930429630625134320180588539459392686425350455321

62629977662665960441550981435850961180153254866649042443251052420197366737753735923059410751031864621855531019149339262612426612213014016215013717311442324280242210345224171232161

614297631615598622337468626302441487628418551635443399437496453179815644315427397184438964132655401825003042234612924071152561211091641341561141708738264259217182341201179223160

126611292021421156911817659971111401211271131058181109100471321527711811538153229266134231207868126921141157242124232738322011586179357650566335

126611292021421156911817659971111401211271131058181109100471321527711811538153229266134231207868126921141157242124232738322011586179357650566335

126611292021421156911817659971111401211271131058181109100471321527711811538153229266134231207868126921141157242124232738322011586179357650566335

126611292021421156911817659971111401211271131058181109100471321527711811538153229266134231207868126921141157242124232738322011586179357650566335

20151416225319191958219210751739249912241922174224691682185820651627146816431864163881325982044100517451273770163433814691109188559817751161873169114291743356903440473597436541493638350170102610278336271061696622895506

20151416225319191958219210751739249912241922174224691682185820651627146816431864163881325982044100517451273770163433814691109188559817751161873169114291743356903440473597436541493638350170102610278336271061696622895506

14981110154913641341157277712471803889139512301746126413131470121010751264137311376591774140168712128615521130234102486012404421157831687117210471190280671317343453289395376466250118745729560407702506446664383

1458110015171350129615367621218177686613771205171812261286143312001051124813511114652172313686721187841545108322898485711974341108803682114110221159275662307336452286387369453244112732721537399683498427643382

40103214453615292723182528382737102416222375133152520747640343849285312531591071387136613823819819211

5173067045556176202984926963355275127234185455954173933794915011548246433185334122185041044452496451566183301865193825537623212313014414714611717210052281298273220359190176231123

5173067045556176202984926963355275127234185455954173933794915011548246433185334122185041044452496451566183301865193825537623212313014414714611717210052281298273220359190176231123

63674252659165157040678938635070778738815352556272575302586165095552454348275391481522897752678032465183412425425347115846183792594819605450385524935426430054221263266713401364190014951790156417231205546307330752422186737412344201125341967

2289519924824015811316722112315519221514617322817916312916317056244231113211122105160321228320472203114461691261953468564737425353613721115136874613387707745

2289519924824015811316722112315519221514617322817916312916317056244231113211122105160321228320472203114461691261953468564737425353613721115136874613387707745

2289519924824015811316722112315519221514617322817916312916317056244231113211122105160321228320472203114461691261953468564737425353613721115136874613387707745

28391760326530023531314117342446375518592634259834712460264930892498208522632521231597236733117159724401918113924745412281168128319272651172311642511208926606071311674637906742887719796577236144614941203989176110618381243848

3801593822874543612102794012252613054122682954122692152553213109744733920629420213832359291176408113316203852952623835911582536789955211167311501541241002221029514460

3801593822874543612102794012252613054122682954122692152553213109744733920629420213832359291176408113316203852952623835911582536789955211167311501541241002221029514460

2459160128832715307727801524216733541634237322933059219223542677222918702008220020058753226277813912146171610012151482199015052423814233515201079221618272277548119659258483965379266768551020512961340107988915399597431099788

3922604555105044642693465642843963995264454325184003323023943181405034982523403171824038736324043712836226117840128539985173791251231021431111248736214212179153258182159167141

1671232071462001741151792321231521552071171511591361251321121343921820212813789661462614190198531961046313312916938763629444652635021157075814611566377856

108441591741451326311415577102115131103133152128898595111311721287995132361082096631523913377691479798254730213838402831229617054367846395526

17921174206218852228201010771528240311501723162421951527163818481565132414891599144266523331950932157411787171494349139011121636594164410787691535131616114009004474096344675574654803801459519837656541088665508799565

33002397312732653269349020162457381118992563277235712696303931922875229524352707233012613835343215362532208412982713585221520282913961259620181283274620852567622128861068095771185079286659128915121445113283218471196110312141074

84862486979286289762470898249664168687275179980565955763263261027598788539365957934571915258252873927771452534265458366917730215517623818920020620216474385422275205498306286289278

3783063623904414482733314542192282694033213233662762302572652761124994261972982231593466926218529511032322912225225130475132637410476758381803418022012289213118115125109

47031850740242144935137752827741341746943047643938332737536733416348845919636135618637383320343444167391296220402332365102170921021341131251231218440205202153116285188171164169

43131944148349244729943557229038745057841846547349739436045332819661051123040031720445877339274427133401291203406351408911951101251471241421481387853221235210136280197191212197

24916725328029026616627535017524428032024725125928423820428320511738331014025720011929342198164247872391691062402222415112761608269817890513012615712094174120122137115

18215218820320218113316022211514317025817121421421315615617012379227201901431178516535141110180461621229716612916740684965655561704827239578904210677697582

488356491465503462262327509272350403535373383461423322323361380165575441209371274184325883122274271373833011464002533858218410198136971051001207940199168166135238157141179135

488356491465503462262327509272350403535373383461423322323361380165575441209371274184325883122274271373833011464002533858218410198136971051001207940199168166135238157141179135

15331098132615251412168483198717488411185123315861154139214531296102211201261101262516631595704110291456512112689829991320414109890159212868981105272607244281436301403338406270122707620481356831536485534464

15331098132615251412168483198717488411185123315861154139214531296102211201261101262516631595704110291456512112689829991320414109890159212868981105272607244281436301403338406270122707620481356831536485534464

672530813822869762528614902478643655912638708868648523560672580275111487938965239330270711354836071221076349628362550168314332416113221718824317625714873357399328247420290274324250

672530813822869762528614902478643655912638708868648523560672580275111487938965239330270711354836071221076349628362550168314332416113221718824317625714873357399328247420290274324250

2501442553063192491781962681572072393322101783122041531452272071013782911342171271042053717811021375266165942191632224810549456947616082422310813010984154978810268

207120213242260217151156222134166200264175147247161123118190165853102381051761058417031143931775922413080181136188428544416337515167301690109916812175747757

3130203065374238242924203916202926121835181629371524101122317192393219361225258148554436421081271566913

5632735280584245583035667351437230433279632011968315428266210472262166931145329421114915211017151765303522164222232613

12515166241014121216191825121914810686271612531412212118615151137735332332525957101062

2712284218311616351519152522203617122315224193511241771661892742923931222081254941067427161371515141010

28303828323523213319193038262443311718282014644314271412213272223113322928243382495133681144201810161320121312

531139745932182260295350713528484331172734255239224233143772220241346201144296141610913131117211031823291526129137

43244264593227404623413968353165433027374216685329412220356351736164235143827346205461010915127182118163322142511

43244264593227404623413968353165433027374216685329412220356351736164235143827346205461010915127182118163322142511

15162061513713201413172225201610761512631114165101310324322115119191565245356312571711695

15162061513713201413172225201610761512631114165101310324322115119191565245356312571711695

15162061513713201413172225201610761512631114165101310324322115119191565245356312571711695

23215330426529629317222234517321821933325827934023322121023322810143436615424414010229441210124294812851681081971922645113465459083796091562313916510596141108107109115

139971951662041991181572241081491372201651842351481421391311365728025210916291741822814472203571931077313211418534964129534957496241139211372689769667189

1592218241612173011101324172326171617142173316112296171186214151339122351072725874279941766106

102125135387107714831269647216851135729114513811412912212323422683772452

76711301021281195895134661029113511910213297948890824116013868996247871989441173412561358871112215422203335422841254656853445547424541

3815314139594537532130264721566528232823267719025301716716282151144025342119417301061198101010514287131814141140

93561099992945465121656982113939510585797110292441541144582492811213665291249261356578791738241637342211291510475233284439413826

93561099992945465121656982113939510585797110292441541144582492811213665291249261356578791738241637342211291510475233284439413826

1752172342452392071711832691342051802251452312002011421991971336727121197175121861953515012318151190152761981371784380413756549853794424989910766118747310462

1752172342452392071711832691342051802251452312002011421991971336727121197175121861953515012318151190152761981371784380413756549853794424989910766118747310462

40365165515635425731384843347144432952452019584222502625467302943632241651423271468169151110105211823191710202418

13518118318018815113614121210316713218211116015615811314715211348213169751259561149281209413845158128601479514636663529404583426934197781844710164538044

72625671105804790975675717652581017548454654248410654663918468654171197345206853649341711272471418107243436194328251918

72625671105804790975675717652581017548454654248410654663918468654171197345206853649341711272471418107243436194328251918

72625671105804790975675717652581017548454654248410654663918468654171197345206853649341711272471418107243436194328251918

72625671105804790975675717652581017548454654248410654663918468654171197345206853649341711272471418107243436194328251918

72625671105804790975675717652581017548454654248410654663918468654171197345206853649341711272471418107243436194328251918

805478679563493580364755105595883564747657125103703955483063167543632241481458664592923192022202023135243437246025262530

805478679563493580364755105595883564747657125103703955483063167543632241481458664592923192022202023135243437246025262530

47285552644733175428333867383655412832474515644924313121401045313811322883645266171411121716111881132422153411151718

47285552644733175428333867383655412832474515644924313121401045313811322883645266171411121716111881132422153411151718

47285552644733175428333867383655412832474515644924313121401045313811322883645266171411121716111881132422153411151718

3326231531161618268141738212228151915182610392115241792363012251192062221193129885495541110159261411812

3326231531161618268141738212228151915182610392115241792363012251192062221193129885495541110159261411812

3326231531161618268141738212228151915182610392115241792363012251192062221193129885495541110159261411812

53014061673842936457645944916295833343574160402560873658480946674005374640633761334927386452483228883655293728683843233845753590417721554734327421493499329056652452436610441027242519552340164215881109797477023102037219838932649201020611703

15658517363143358387586113977569278497010450533412114921804028652047354922504335635968302381213191613191112392923274021312726

15658517363143358387586113977569278497010450533412114921804028652047354922504335635968302381213191613191112392923274021312726

15658517363143358387586113977569278497010450533412114921804028652047354922504335635968302381213191613191112392923274021312726

1515039704713433647655581307054857647659449522810714621763923641741304422433634635056252271011181313161012342817243317302525

1515039704713433647655581307054857647659449522810714621763923641741304422433634635056252271011181313161012342817243317302525

58123169219113397272251011614341513655771912511221331516374121

58123169219113397272251011614341513655771912511221331516374121

31925637829542335622030540518325424834519823427824021920522722411132927814918120015820792255162264110290187992081872931112046857113113113921246668220162114111199146119148138

1391221711371821718314915571111891507680112939374105734512992637774699527877010124114995296771333675241937383824372019744141386954314243

133120152131176161781341486710384143717010488857210269441218760716961892379649722108944891721203466221936343424371818693640346452273343

1834403917232029239171211121013151719169814613715107313171321118922820584236434121225291161610

1834403917232029239171211121013151719169814613715107313171321118922820584236434121225291161610

46234119553226384322272650142230262025222674026181916232761817329312810231738112468117751385241314121491277

46234119553226384322272650142230262025222674026181916232761817329312810231738112468117751385241314121491277

191522236227122642171519321417221321626127282214191651561815276341581717301216621011106104314710111485310

191522236227122642171519321417221321626127282214191651561815276341581717301216621011106104314710111485310

50484950427920414019442750312139342722382222393315262223408301525532332129303261867121013101058191411927244716

50484950427920414019442750312139342722382222393315262223408301525532332129303261867121013101058191411927244716

6219661051574857510858234185365864864265455132921442155145249

6219661051574857510858234185365864864265455132921442155145249

6219661051574857510858234185365864864265455132921442155145249

86659590130111787211752778211774777173477469672711311343495830581893487431773722555672185723173634221426144594334415747403728

86659590130111787211752778211774777173477469672711311343495830581893487431773722555672185723173634221426144594334415747403728

86659590130111787211752778211774777173477469672711311343495830581893487431773722555672185723173634221426144594334415747403728

86659590130111787211752778211774777173477469672711311343495830581893487431773722555672185723173634221426144594334415747403728

64407344895245629836515752286060484844365130605530374734291550265125643913353862133412927241911271410392623175433382225

64407344895245629836515752286060484844365130605530374734291550265125643913353862133412927241911271410392623175433382225

64407344895245629836515752286060484844365130605530374734291550265125643913353862133412927241911271410392623175433382225

64407344895245629836515752286060484844365130605530374734291550265125643913353862133412927241911271410392623175433382225

3029392422221422352415202620173526311317339271813182125253225183830351212221626443891213173443341835485216151912104742

3029392422221422352415202620173526311317339271813182125253225183830351212221626443891213173443341835485216151912104742

3029392422221422352415202620173526311317339271813182125253225183830351212221626443891213173443341835485216151912104742

3029392422221422352415202620173526311317339271813182125253225183830351212221626443891213173443341835485216151912104742

18601358276315012853303619412281417518651479143422051257223418701251146516641236124895323451943109813621132933149751616371209145664817251181741125312522282999216740837370553947232554629719517239056567801157820517735544

491646454737275050275052535450485424426243287543154129232742829521137101734242812391414271613151086292417133022132112

23722252021162623111631243222302112212715133520619131410111161751651020111741496141081055417159813106116

23722252021162623111631243222302112212715133520619131410111161751651020111741496141081055417159813106116

23722252021162623111631243222302112212715133520619131410111161751651020111741496141081055417159813106116

269242027161124271634212922281833122135281540239221691731713356215714131182558136555321298517127106

269242027161124271634212922281833122135281540239221691731713356215714131182558136555321298517127106

22517112111816189251222161412258132720111913416135931111264152511876143683322229455128664

4479653897997614684888421105634862926323542112253233135354142

1164343135334544224541564112143211351321431622111

1164343135334544224541564112143211351321431622111

1164343135334544224541564112143211351321431622111

1164343135334544224541564112143211351321431622111

631861375121491216171091559121033151254871527105571155101037353254163452571

631861375121491216171091559121033151254871527105571155101037353254163452571

631861375121491216171091559121033151254871527105571155101037353254163452571

631861375121491216171091559121033151254871527105571155101037353254163452571

17203022292116183421302236322115321539331081523151817121531712254202472021161161213304738111113109172261612

17203022292116183421302236322115321539331081523151817121531712254202472021161161213304738111113109172261612

1114413410557141671131243312142451123133221

1114413410557141671131243312142451123133221

17192921251715183117201731252114281433261071423151517111531710214172171919151121092546371191096142061411

17192921251715183117201731252114281433261071423151517111531710214172171919151121092546371191096142061411

1301885626013110858139127898963231726474768580725619513984686344956144101629433127813852905444363221335461321832096751244379273311054

53852842463522967564337261073229343428473116100632738242651252850243820563313194325321281229238113085351515192912104320

53852842463522967564337261073229343428473116100632738242651252850243820563313194325321281229238113085351515192912104320

53852842463522967564337261073229343428473116100632738242651252850243820563313194325321281229238113085351515192912104320

7710327836685629727146523712440354042573341409576573039184436165138561371482533472912235101126235105312432369245015236734

7710327836685629727146523712440354042573341409576573039184436165138561371482533472912235101126235105312432369245015236734

7710327836685629727146523712440354042573341409576573039184436165138561371482533472912235101126235105312432369245015236734

56346631897655104112594650693753573238373838397266384530373129634563246444285629672644616222117132249352727344331282715

56346631897655104112594650693753573238373838397266384530373129634563246444285629672644616222117132249352727344331282715

56346631897655104112594650693753573238373838397266384530373129634563246444285629672644616222117132249352727344331282715

56346631897655104112594650693753573238373838397266384530373129634563246444285629672644616222117132249352727344331282715

160110962035133325412783177719573835165512491228179510472033165710501292145010161094679202417099531191100375813464331417104812155701470101064610861075161291319953543145844494192684152601701580778573677981715431553450

16686248230269131126320213284801471402391391903231293771591381281013193818125817664125492088113472183911001141121745217048474244592667292210195599194118377448

12872194173223119924616712674241121101811131522649935813511190822763496822015854904717653108631627687948914636130363628334620562516828348788076316239

12872194173223119924616712674241121101811131522649935813511190822763496822015854904717653108631627687948914636130363628334620562516828348788076316239

1272042322812266612212224114117312111121141214211

1272042322812266612212224114117312111121141214211

3813525026108153259563322572436532413232536194230113614934232282292114122016281339101013101241046181271212284118

3813525026108153259563322572436532413232536194230113614934232282292114122016281339101013101241046181271212284118

11611476421264911432551312524222323241135122

11611476421264911432551312524222323241135122

11611476421264911432551312524222323241135122

157912301552111941021532491081201381931261241621319412713510366257149911091019010335145126150581601105396121185671124022565752244226191166556779054555336

157912301552111941021532491081201381931261241621319412713510366257149911091019010335145126150581601105396121185671124022565752244226191166556779054555336

875513296119111558212872717811261639685546877684116810050635155692683801014498773469671034264211132382916271512604337485035312621

7036985992834771121364960816561664640595835258949414650353496246491462331927548225481911241923815117562219294019242715

6052895694827075886270577355445957815544546481636356359350109846962749057414851933042122145462116313111472646504242242428

6052895694827075886270577355445957815544546481636356359350109846962749057414851933042122145462116313111472646504242242428

6052895694827075886270577355445957815544546481636356359350109846962749057414851933042122145462116313111472646504242242428

20207219484225564337332865192435323312243626844817361826161533362721492448367411279141493174428181729252625173

20207219484225564337332865192435323312243626844817361826161533362721492448367411279141493174428181729252625173

20207219484225564337332865192435323312243626844817361826161533362721492448367411279141493174428181729252625173

10850149631828562205156958368145556379635754455429129575580435746169260953313794345364129316916133417291622209684838635636254121

10850149631828562205156958368145556379635754455429129575580435746169260953313794345364129316916133417291622209684838635636254121

10850149631828562205156958368145556379635754455429129575580435746169260953313794345364129316916133417291622209684838635636254121

87763289162014417201101952157766959357374746614018034584729024865472868027984794674863168571525555075762305994453376284916676271385177157268188198122171104661054414251259544281186239267

87763289162014417201101952157766959357374746614018034584729024865472868027984794674863168571525555075762305994453376284916676271385177157268188198122171104661054414251259544281186239267

87763289162014417201101952157766959357374746614018034584729024865472868027984794674863168571525555075762305994453376284916676271385177157268188198122171104661054414251259544281186239267

5787911518155111512301885881384552281434251063766526146434412234242725481733

5787911518155111512301885881384552281434251063766526146434412234242725481733

5787911518155111512301885881384552281434251063766526146434412234242725481733

20715834717728632414229037519719120731515316018717516912912816210334220316415413010814357270157161772431816813217127489182485084615061624237162107941061241086210044

20715834717728632414229037519719120731515316018717516912912816210334220316415413010814357270157161772431816813217127489182485084615061624237162107941061241086210044

20715834717728632414229037519719120731515316018717516912912816210334220316415413010814357270157161772431816813217127489182485084615061624237162107941061241086210044

1331391351151741486317018410811310215412976110105668610990652231205911060578226136711403817010341101921834991342269565223322917897065718061645839

91998579969732951015658518880426269434156502914376406336264018714393219749245561983348221141322310191612474233385037283617

53447243531175984241153443311333251322424222231114143

53447243531175984241153443311333251322424222231114143

53447243531175984241153443311333251322424222231114143

869982759290309198515550877337536139395249281387336593323391768409019924821535994314420113932218161511474129384937283214

869982759290309198515550877337536139395249281387336593323391768409019924821535994314420113932218161511474129384937283214

869982759290309198515550877337536139395249281387336593323391768409019924821535994314420113932218161511474129384937283214

4240503678513175835255516649344836234553403680441947243142865284717735417463185164312112824291313135422832333024362222

4240503678513175835255516649344836234553403680441947243142865284717735417463185164312112824291313135422832333024362222

4240503678513175835255516649344836234553403680441947243142865284717735417463185164312112824291313135422832333024362222

4240503678513175835255516649344836234553403680441947243142865284717735417463185164312112824291313135422832333024362222

238179292315231818182616161518261111111011342611810611212121210181721013321741765224212955129274

238179292315231818182616161518261111111011342611810611212121210181721013321741765224212955129274

238179292315231818182616161518261111111011342611810611212121210181721013321741765224212955129274

238179292315231818182616161518261111111011342611810611212121210181721013321741765224212955129274

238179292315231818182616161518261111111011342611810611212121210181721013321741765224212955129274

765558407157367168333532493138436819435231215534222740223814282661144733232631602632415211719111386431919183619292221

765558407157367168333532493138436819435231215534222740223814282661144733232631602632415211719111386431919183619292221

765558407157367168333532493138436819435231215534222740223814282661144733232631602632415211719111386431919183619292221

765558407157367168333532493138436819435231215534222740223814282661144733232631602632415211719111386431919183619292221

765558407157367168333532493138436819435231215534222740223814282661144733232631602632415211719111386431919183619292221

43327954436547043435942855830144135057535437835736730730036024927854035729238122328232519944033837825042528323336829446211323788822681751391151256546249164179184279267214149146

53742644768544610214332121663434441713668441122323114222353

141321442311321213211321411236222212111111

141321442311321213211321411236222212111111

141321442311321213211321411236222212111111

523413246243135713311914623234132432241111113112342

523413246243135713311914623234132432241111113112342

523413246243135713311914623234132432241111113112342

22317728619421623923722831616125020928518220816222518916321311419129018019820612520418316325421019318822813815718815823255110404816410778596835171417195851411581366690

22317728619421623923722831616125020928518220816222518916321311419129018019820612520418316325421019318822813815718815823255110404816410778596835171417195851411581366690

8777108113961027210215067130891707310374127767190567017079551075351793710582795310251618780111355317288553402533168693536346274643437

8777108113961027210215067130891707310374127767190567017079551075351793710582795310251618780111355317288553402533168693536346274643437

1361001788112013716512616694120120115109105889811392123581211201011439972153104126149128114135126879610178121205723207954383435199723659517984723253

1361001788112013716512616694120120115109105889811392123581211201011439972153104126149128114135126879610178121205723207954383435199723659517984723253

2059925116725218911819623513418313628616816418514011713314713284248165931699275138331821241816119014473174130222541234733102665854542928104918297135104788056

2059925116725218911819623513418313628616816418514011713314713284248165931699275138331821241816119014473174130222541234733102665854542928104918297135104788056

2059925116725218911819623513418313628616816418514011713314713284248165931699275138331821241816119014473174130222541234733102665854542928104918297135104788056

64278442817540677437594878564244484031473221944926572933237573953185047163838671736149372320101410933272841382726317

141721671251711147812916197124882081121221419277102100100631541166711263421152612585128431409757136921553787332465433844401919716454569777524949

27823152820635128331438938443221025032519018026623132232518115628758216520317111529737878440222128029231920717117014341418025472691501331471221829996215156168204224159101168149

1871463431552212112622772691981271592301471221731622131101171062124621141751078323216429031717518722719214215613211026710315852621181041046887584512792141166158120819190

323552414942195033262119423620242422132014839221120152621229151673022220224612237310511310101175141421138126

14132634151882019141181730711167866323155141016111152731071511246643826373173116128366

14132634151882019141181730711167866323155141016111152731071511246643826373173116128366

18222673424113014121011256131381551485167665101011413942015251122617323537102389556

18222673424113014121011256131381551485167665101011413942015251122617323537102389556

1551112911141721692432272361721061401881111021491381919797922044239216487682061432882881601712201621201541128822191135455910899936577484411087127152137107737984

1551112911141721692432272361721061401881111021491381919797922044239216487682061432882881601712201621201541128822191135455910899936577484411087127152137107737984

1551112911141721692432272361721061401881111021491381919797922044239216487682061432882881601712201621201541128822191135455910899936577484411087127152137107737984

91851855113072521121152348391954358936910921564507512051286432652144948546936512765153833147779620732294354954151886427386639207759

91851855113072521121152348391954358936910921564507512051286432652144948546936512765153833147779620732294354954151886427386639207759

40547635503224414142372640142646433731242528352412221835433040175339653182318554861762011344845354963528112915105749

40547635503224414142372640142646433731242528352412221835433040175339653182318554861762011344845354963528112915105749

51311091680402871741924665552932472672184402547852716421430171464452940266234715159229351311218965062251219273724102010

51311091680402871741924665552932472672184402547852716421430171464452940266234715159229351311218965062251219273724102010

1801153019911467180517591235226121891152138112802099128114281486149010941206142311597592017160681111749788071096403137813001339482145310816831204108716488161189300340947742128586447548630520417116706281716977799643557

72276540647167625246364373345848333240503431846439391834322444564337434331515362133113101717141513157362335324629252925

72276540647167625246364373345848333240503431846439391834322444564337434331515362133113101717141513157362335324629252925

72276540647167625246364373345848333240503431846439391834322444564337434331515362133113101717141513157362335324629252925

31132416232225171219201725132618145182517113525111591413102025211720211921192551155968944516136142214898

411441244149424540271626482132301927222517204939282492019142431222023221230343782085811669112201029182415172017

557471556421464508326438592327440385576361447437522314346417329245510456254350277258315124379330354138399332175350277442122248799721013016710410910056278202175144311190208165134

68549971105765362115517176715650705053456539269184527437424725643561226039174352102243918233926291591511523228364936272815

11151611139610128162924111113111310148613181219781441741481011361117662612346413646948951

11151611139610128162924111113111310148613181219781441741481011361117662612346413646948951

1391323192071123913161210102395111311519161213678312101513951013465565222236688105347

1391323192071123913161210102395111311519161213678312101513951013465565222236688105347

1212185156413181111714146914212615121165693543465382123852514432215456725

1212185156413181111714146914212615121165693543465382123852514432215456725

3218523258413628622331242121232516331232191047392237181822183017291031146192049622972014194310325221014291613147

301541295039302155232719181821191225103013936342131141619162314257231441518415207619121742103231791126912126

231138267745332648226111516423273438242812211221251337121

48941745735035943227337647727636930950530539736747226130135229021941937220227624021626899315295293116339293158307225340982096174171104138891008545226170147108262154181137119

48941745735035943227337647727636930950530539736747226130135229021941937220227624021626899315295293116339293158307225340982096174171104138891008545226170147108262154181137119

73744539577233426235614663395938543949553334625821453326411634385613393615392948102315919191718918526102892626281418

20517716015911714611017418190130971891131461541558210713899881571348095838796271091341064012812155113921303980162668316438492514986255367943575646

41417431484235366236413783334539503133333426564927282533322446362619523226322145112887269105812623271215301613169

10917171716818271514211699194619131416218198211112103107165181655131691312436422181699136465

1601161611041201568710614510012310815411113811716790991121086912611266878858892911680893910288571187010129652130544241283028186155433911463794541

67577749687135488043574395314236574143483625826126393826372151485713683725372874114410112725112615104431729252731242315

67577749687135488043574395314236574143483625826126393826372151485713683725372874114410112725112615104431729252731242315

1619167163691126819928920618791597118891051271213143233710920141377363211813699733

1619167163691126819928920618791597118891051271213143233710920141377363211813699733

5138614252352637543538346722223039343433271871531830282125143935431045341827195411309820188201210232916191822172012

5138614252352637543538346722223039343433271871531830282125143935431045341827195411309820188201210232916191822172012

10157150911531068913116484587698628594575940725830143108495940406820100598828117705154661094666192135262513311511894747495930403528

10157150911531068913116484587698628594575940725830143108495940406820100598828117705154661094666192135262513311511894747495930403528

10157150911531068913116484587698628594575940725830143108495940406820100598828117705154661094666192135262513311511894747495930403528

10157150911531068913116484587698628594575940725830143108495940406820100598828117705154661094666192135262513311511894747495930403528

1613202225281321271319202619101581713141312172010142213104111315616107814192745610435218116101561066

1613202225281321271319202619101581713141312172010142213104111315616107814192745610435218116101561066

1613202225281321271319202619101581713141312172010142213104111315616107814192745610435218116101561066

1613202225281321271319202619101581713141312172010142213104111315616107814192745610435218116101561066

4173874713043493242493764782523242525333023043023282452643322831523673491652222421532328130730028010631423918331024233768166496216112111283827543200159131109239162127157115

9212683665770445991597043996554605837376767216775304746216415696252257150436259591134883817191114176292929255423132525

9212683665770445991597043996554605837376767216775304746216415696252257150436259591134883817191114176292929255423132525

9212683665770445991597043996554605837376767216775304746216415696252257150436259591134883817191114176292929255423132525

147831801051199179142154809876186991029214377104101815214411552777449682911983843583625486691192668141953432724262119735542476946326327

4414152615155401771710123115213022301619814911161712122242217571212141512494851455212762884115

4414152615155401771710123115213022301619814911161712122242217571212141512494851455212762884115

805012749775460789752645713748694596435865513385713149452943155339472153333347408822398832271615211210443630354025173315

805012749775460789752645713748694596435865513385713149452943155339472153333347408822398832271615211210443630354025173315

231938302722142440211793720182617121620111145351012128131242222092317925141920671311105471712610211311197

231938302722142440211793720182617121620111145351012128131242222092317925141920671311105471712610211311197

150136170117158134111133209101138117211125129135120119112151120661411437294112659031100130129401401177814010113825522531675559473434159066533010587796662

150136170117158134111133209101138117211125129135120119112151120661411437294112659031100130129401401177814010113825522531675559473434159066533010587796662

4024342025261018351722235321222924152215271232167181016226191512527191020142959621314111175716121291812997

1101121369713310810111517484116941581041071069610490136935410912765761024968258111511735113986812087109204319295441483627298745441218775705755

2842381615291542241218163713191571211131513151611410181061925156201082213216122436718338977116331

2842381615291542241218163713191571211131513151611410181061925156201082213216122436718338977116331

2842381615291542241218163713191571211131513151611410181061925156201082213216122436718338977116331

571518652540682651456118579638744746169847248255448538646049040626481454826845134128340212948649450215449635021139440760555462712613449141395262022026918313872522472591019529365228234

571518652540682651456118579638744746169847248255448538646049040626481454826845134128340212948649450215449635021139440760555462712613449141395262022026918313872522472591019529365228234

4925756483674364109534666594247805258504755246466215238292413594249197236224437791327148261818917107493130264725182116

4925756483674364109534666594247805258504755246466215238292413594249197236224437791327148261818917107493130264725182116

325374365271350348245886409193226232381248247298256217253272215150500272136258189154240712743362897924918111521721829946148772683873458485511392151541159137147148828419266121148

143131428161935231216122513199819141312812151129197916621423451412295853539453210689147775

432835335636258942313327363339323624314421227141243922162113282631935181426323641599735335937201615931717226236281912

35303746374334767221303340353734382729262210672623232514237303836932279383327313676273167417239822819176733231411

927212278939781156814153569658546671597275553910962296155345514867858174943245354698597181061601528826313420926393813486382131

34161219161066133101310221611492991123159211514173143241016175211061512101318442012281768643383372816820

63158945778101433061085450551117267745454577466411408834715135711764109991964594744497919516420311531413752425594575034041403471651163651

281724131621185427101122281271991416613634156179814118141651011511121626454433174739613681114650171286

1650281126241910923142017239131511112311111546105181514176224911111595161433656053534811183142725138611131174726812

45355547644834677432445085556249423033473818814932362331365462955134735123437572128917241427182396551921213523102619

45355547644834677432445085556249423033473818814932362331365462955134735123437572128917241427182396551921213523102619

3351715234321302520291832391517171130710152126188101013618121961914911233151429117967411597131191586

3351715234321302520291832391517171130710152126188101013618121961914911233151429117967411597131191586

1197914014316214511313817989102951418811111011870941178857148135619781598934897590371098453889213954712932432950363431151095642519853565245

111232291412122332289142523171924712259121833131318131459131192417926151991676621036631071013131510116

281026334846322925919272317112213142016221031231524141019816613725175131528920529486422281661016131168

30819223327213035182822361528273222193320172818211414102572921231017131223293818831297127108517136112912111110

50496359676048568734463257335542492743433718716112463526311435354311433727263354182714121916202014155542020174013242421

22214727322221822027328426520716816424214613416117817311911212921920615422216113927814428224021619828923713912196132223127162585613215592757044501398513817015017013410479

22214727322221822027328426520716816424214613416117817311911212921920615422216113927814428224021619828923713912196132223127162585613215592757044501398513817015017013410479

4425783664771437361983771554231404389201626155161614949361925324311612150237104488020378377701827891265532332529543472985684763722

4425783664771437361983771554231404389201626155161614949361925324311612150237104488020378377701827891265532332529543472985684763722

351949274555102554778265733302034356216122011911121003630136381708276281617227551628645351111753903316201523382454703953492716

96299192241181420111422121168274463654491365615733445227632212549192419710363622161310616101828173127106

1781221951861541431302112041091319318710410312113584999610364190138731121038691391249514852133914176951405092402943293743371921855166729486586757

1781221951861541431302112041091319318710410312113584999610364190138731121038691391249514852133914176951405092402943293743371921855166729486586757

1781221951861541431302112041091319318710410312113584999610364190138731121038691391249514852133914176951405092402943293743371921855166729486586757

4826159405611836265374507984329253043037032555893357632941688652071049323333831075944321040384188737912153664099386105931732223667108398936728727751996318300638341347081854813305221932716411641651339047010137922389850275109428384309176122170211255312424249183806041279610124087077627261339466662528999400152184779112811695191134097108623975323637002317137371193086118969873878421719891573539177408115000431734906

510028566625643266125366407236137489337345864667702441484995565044533876361741743892174591296735302543583200180837418304840310750371823497635611965377135784994118026591440104314431209141311791430883511278832741870195132881595148219991526

510028566625643266125366407236137489337345864667702441484995565044533876361741743892174591296735302543583200180837418304840310750371823497635611965377135784994118026591440104314431209141311791430883511278832741870195132881595148219991526

510028566625643266125366407236137489337345864667702441484995565044533876361741743892174591296735302543583200180837418304840310750371823497635611965377135784994118026591440104314431209141311791430883511278832741870195132881595148219991526

510028566625643266125366407236137489337345864667702441484995565044533876361741743892174591296735302543583200180837418304840310750371823497635611965377135784994118026591440104314431209141311791430883511278832741870195132881595148219991526

1326930328617616212681186851191192221121111451011051109686562571478710570561052022283133491827761837416336794135455131292415168710155608136345031

18868652443215128402111167312493061123416371736264215431889224017111505120215341588535373128631201168710757131331313171210841929713185914016191298134819154701050563330404470484383523322178110513196457641396592559688514

115064113971359127612108728381632783103810061435895112112389929148079387914271543135858594079141084817597872610983171059783473994821109525460327324134626130226734319294570642392403712374314422381

2031032661902551911201172491181671582811472032241301431181171643841122611613979591003120911218668199217731241332213710765385844553440342296136889911860538463

2711443333383342361991983742052582713802113003032711942112391847864134023624320110425553254172260108283180105179186247671119070896066858449431381931581301639110312677

693397934904844741547556103648765060993360767277550948250257552125711049013846325032255389767745267726562343132451149561716134715614621516222116820513377348380242243396241196289217

76563794912048877155355749514617177681131633699725739533667675558354144290041661248124156414178847875430377147231058252173615536225218328616125421321113881444503290252422201223340243

3029261327703080270135352271200837302135221224463586249324743431276421122664242226231132374339901335230925511041197255822881749246387320301743113320871975231269620689647091416774114375399956225816032435129211081737118991514731311

3029261327703080270135352271200837302135221224463586249324743431276421122664242226231132374339901335230925511041197255822881749246387320301743113320871975231269620689647091416774114375399956225816032435129211081737118991514731311

122614011078127212061452905764152392293610021452967104913529487711020966107142816461612566962923438802233976711967366841665448833809862242763350259538305403297370230103660973488442698439386617480

122614011078127212061452905764152392293610021452967104913529487711020966107142816461612566962923438802233976711967366841665448833809862242763350259538305403297370230103660973488442698439386617480

196118152209208239150101254220153163237150152257169141178190203872403431151751836512533176991887015885781621511415714587531285084636637141191959762120824813686

10301283926106399812137556631269702783839121581789710957796308427768683411406126945178774037367720080061277929668358037067165872118561826320641025531923430419389541778391380578357338481394

1576105214611588127618241199109819411026111612391893133212361836162211951372126213765991845211366611751414531103228811418831264447105195658310591007127339811465493927724116714005442891348411284675591879644463760720

1576105214611588127618241199109819411026111612391893133212361836162211951372126213765991845211366611751414531103228811418831264447105195658310591007127339811465493927724116714005442891348411284675591879644463760720

201144149191141193167154241116146162225152154185192138166177164116190210621601375813538151131160481451198216012114636105523510046724862342010311680899864678589

1375908131213971135163110329441700910970107716681180108216511430105712061085121248316551903604101512774738972509907521104399906837501899886112736210414973576723655993524822551147381168595502781580396675631

227160231220219259167146266187160205241194189243194146272194176105252265103172214721383717115523260138122102195159177561596558106586956854321102178129751601066696111

227160231220219259167146266187160205241194189243194146272194176105252265103172214721383717115523260138122102195159177561596558106586956854321102178129751601066696111

227160231220219259167146266187160205241194189243194146272194176105252265103172214721383717115523260138122102195159177561596558106586956854321102178129751601066696111

804060228207962478728880540154779612453972356771993875968416817878425626668970965323372611016942340956249530626036184135862465334682727375837452338797353574367141445374519811808351819962481207423031529670386649262849254841243205325529992764

113270911631280107510666836251244646959911139688310421028790746926106676748616081465582832726375107516684565282932579258947699666196022055829124769137932628236124591562723380392639461536431408

7844618368707317254604018564396486509995796917045405126737265283111078103542858250226181610257847955321354039332369943464013536322418955329324218128518966394544269266429331395302281

7844618368707317254604018564396486509995796917045405126737265283111078103542858250226181610257847955321354039332369943464013536322418955329324218128518966394544269266429331395302281

7844618368707317254604018564396486509995796917045405126737265283111078103542858250226181610257847955321354039332369943464013536322418955329324218128518966394544269266429331395302281

348248327410344341223224388207311261397304351324250234253340239175530430154250224114259642671732761122521961532972273208519567581388684101765625168179111126210130141129127

348248327410344341223224388207311261397304351324250234253340239175530430154250224114259642671732761122521961532972273208519567581388684101765625168179111126210130141129127

348248327410344341223224388207311261397304351324250234253340239175530430154250224114259642671732761122521961532972273208519567581388684101765625168179111126210130141129127

376425573359406834274442238923544077201232303069419634374053399939042520289928822256165244084146180627162026999266958326472382333213372453196117393555257528225591541933898143467111109181055708283148520371292103317411173127113251187

376425573359406834274442238923544077201232303069419634374053399939042520289928822256165244084146180627162026999266958326472382333213372453196117393555257528225591541933898143467111109181055708283148520371292103317411173127113251187

376425573359406834274442238923544077201232303069419634374053399939042520289928822256165244084146180627162026999266958326472382333213372453196117393555257528225591541933898143467111109181055708283148520371292103317411173127113251187

314338465455311332215321379176345389349313267253330200340287188165362333176226230162308582062102561032772042193242402784114574981437913810889633217117815693135156230143138

345022192894361331164110217420333698183628852680384731243786374635742320255925952068148740463813163024901796837236152524412172307612342176175715203231233525445181396859800129159297281096664525113141859113694016061017104111821049

29482629346740803155315422012378404917552905261841193151315729883008225227193027216115334527361016152559246711642327588252821752476993239518621574259923642745635153271363113108681001852847544277172920191094104316471484138611731117

51650154565052852833543265829948143270452247047351939748957439027674957626740744317740710848834946116038431724745543447010233411912024815021519614510535361575200228295273244235190

51650154565052852833543265829948143270452247047351939748957439027674957626740744317740710848834946116038431724745543447010233411912024815021519614510535361575200228295273244235190

51650154565052852833543265829948143270452247047351939748957439027674957626740744317740710848834946116038431724745543447010233411912024815021519614510535361575200228295273244235190

24322128292234302627262618661946339114562424218634152629268725152489185522302453177112573778303413482152202498719204802040182620158332011154513272144193022755331198594511106271878665670243924213681444894815135212111142938927

7044106111108866359143487881121969485796065666730172117478867274210845061307339436480923038341858401913181210338729344961363320

7044106111108866359143487881121969485796065666730172117478867274210845061307339436480923038341858401913181210338729344961363320

23622084281633192519254018031887324814082346210532942533259324302410179521652387170412273606291713012064195796018784701956177619548031938150612842080185021835031160560493100467876764368442723213351357865781130311501106905907

8758719861164908895569650110752485473311368988748889076997519156014581177104245670669134870217969261865629767752543173468275218345722716637221925425025317276461455308235471379372324332

14871213183021551611164512341237214188414921372215816351719154215031096141414721103769242918758451358126661211762911264115812985061261981853134611681431320703333327632459513393431255156874902557546832771734581575

19612721819621521812812024212614117322712516416314010814512113955473202921428765113212261251908219711190203143187311144432837844224032199014783809787627052

19612721819621521812812024212614117322712516416314010814512113955473202921428765113212261251908219711190203143187311144432837844224032199014783809787627052

19612721819621521812812024212614117322712516416314010814512113955473202921428765113212261251908219711190203143187311144432837844224032199014783809787627052

19612721819621521812812024212614117322712516416314010814512113955473202921428765113212261251908219711190203143187311144432837844224032199014783809787627052

57294073618761785613648238914161710534655051458370125052501653125014411547094763400922958064638927414527398819574187100344003445477315854600351322424611391647061123278414261342208212061612125415581048453272029301998175031472134171021792100

57294073618761785613648238914161710534655051458370125052501653125014411547094763400922958064638927414527398819574187100344003445477315854600351322424611391647061123278414261342208212061612125415581048453272029301998175031472134171021792100

57294073618761785613648238914161710534655051458370125052501653125014411547094763400922958064638927414527398819574187100344003445477315854600351322424611391647061123278414261342208212061612125415581048453272029301998175031472134171021792100

5553397959846010541962753769407369143360489444576800491848415120489940174602463738822228766461812667439838951885407596942363335463915304431338721914496378845331092269813661312203411601588122615111022436261028121905165530382081166221222065

2357178126982713238828221650183030691434214319402874217521712177216118761956199616569743049276011721910190780617554231831138520646531957145310181931169119754631123575566863466682529688442200110211268057041295886714882799

790536817940795872515559968470725634977794651731785572745676547321118887841962650226664112960446762221457946432263157063216039419217228317623020822218356390426255233431310245318286

104167111221123102011146466971238619854831135891785010309057118388788023511467117552390570233773718682062285230579761636082567187320953825625542523330821326018488501532357365566408312405317

486360496453465614309326644280423398576372444466371321371361320186757575204353298206335873813373961374023461803892944131012541271271681011321031188429230261183144282156142172160

8796318517817518536496619955577496541015660725716677537692726557396120379334960448627060714460052470522169650831172056264015938921619229518423617322312963387467305209464321249345503

176942031681942071228819110515712621213417519211598107126127674002087412993721123416411013455169126511151281733186603048462428472617110118939510953485735

176942031681942071228819110515712621213417519211598107126127674002087412993721123416411013455169126511151281733186603048462428472617110118939510953485735

29633118311433143042299822692796371716072350213633292663254826932874207922222612192012983420262012982016221611312173499213621462182732203719351562260420172472570112256658311868178367057654962781279110088075715701340137710901109

74859088884770772060863989943563454589269469462167151057765950432098570632756946228149212261050658520954950832477854763713931513816527720323617220214069359295240223387300352278253

74859088884770772060863989943563454589269469462167151057765950432098570632756946228149212261050658520954950832477854763713931513816527720323617220214069359295240223387300352278253

74859088884770772060863989943563454589269469462167151057765950432098570632756946228149212261050658520954950832477854763713931513816527720323617220214069359295240223387300352278253

74859088884770772060863989943563454589269469462167151057765950432098570632756946228149212261050658520954950832477854763713931513816527720323617220214069359295240223387300352278253

15682004154617161605163711211572186578711171054156114131230138316121096115713789916691561130266692212476211213256997114611213449711058944138110421230266533294298644414390365386240151559525425340819739773587610

15682004154617161605163711211572186578711171054156114131230138316121096115713789916691561130266692212476211213256997114611213449711058944138110421230266533294298644414390365386240151559525425340819739773587610

15682004154617161605163711211572186578711171054156114131230138316121096115713789916691561130266692212476211213256997114611213449711058944138110421230266533294298644414390365386240151559525425340819739773587610

15682004154617161605163711211572186578711171054156114131230138316121096115713789916691561130266692212476211213256997114611213449711058944138110421230266533294298644414390365386240151559525425340819739773587610

64752468075173064154058595338559953787655662468959147348857542530987461230552550722946812152949447617951736929444542860516527413412026520021016817711658361280215194364301252225246

64752468075173064154058595338559953787655662468959147348857542530987461230552550722946812152949447617951736929444542860516527413412026520021016817711658361280215194364301252225246

64752468075173064154058595338559953787655662468959147348857542530987461230552550722946812152949447617951736929444542860516527413412026520021016817711658361280215194364301252225246

228219240223282213233213414166257183300247239273226173172231171132257226139184215931764419319318057155132101166154197651144551916489656543191368380581491258180103

4193054405284484283073725392193423545763093854163653003163442541776173861663412921362927733630129612236223719327927440810016089691741361211031127339225197135136215176171145143

16328571960849148346015239951221267171403411852411494207139141771564810726421198863107234912602171311930111610214245651142703131851913103878553547574698285501045843597815798569153987675061112795623222397716509919401075104289570716676109539411888481446124985350872865313605350519161388401371560795283274434903435492363204228717157458516045260801415935252724826292633339690378499746761522

1221511374119441146110982102948188118141278352967725999612617768583339141772873158809846966374127202967549425964858220504574301922794772787992301373806924596691987107877022943935267921663147266727172266236114041048437732102848393058793869412040343834

1221511374119441146110982102948188118141278352967725999612617768583339141772873158809846966374127202967549425964858220504574301922794772787992301373806924596691987107877022943935267921663147266727172266236114041048437732102848393058793869412040343834

357432743192303729652983224834973454139419962716316322842313240522031941238824461789117046742042121717432422137221555392099211221817861884205917902649192723165971013647664990740671647633396308112479875992516321156124710681092

357432743192303729652983224834973454139419962716316322842313240522031941238824461789117046742042121717432422137221555392099211221817861884205917902649192723165971013647664990740671647633396308112479875992516321156124710681092

73476836725168096675596350367052769032164618610277194306481454304421433252974856401524141339142952428378448723103430611604809445047041834445040233590542443445210145524211674123117261591168012981395779617267318901675252835642178236124672265

5383496851455114493245343895523756812379331544555529335037294090332132254045375029581839958431711824274338982241327684734403294341112873151302228544027328137431068172712059511278115512529631045573457197013571243183026191697177918181712

863107610447838996345579619924356507091028441534588657615649567507287136752023750046148051916361058657521357849740577057070218032718014522119319618718311570352247222281483261318319339

110179210629128447955848541017402653938116251555175244349260353955028824406043675415133825111507595707183347215043316274937652073672891352272432321481679190351286210417462220264330214

1294126415011615134213489041265163968611111178173510951206130611041042112411678335432231121261495892657096922310397161107393104684258611258361244242501358271431336366321333229123580522414477683535512499477

7898088829767947915287929724396476971014631675804648644674703477298132171537557959333458313362141569722560850733678251175214332321116723619422319720314165359320247291438329315296279

50545661963954855737647366724746448172146453150245639845046435624591049723937933323638690418301410168438335250343325492991781471041951421431241308858221202167186245206197203198

20451589252620092162166113052211258190914092095273810341317166411421204155413091250526512313028701257112314821158290178411221493744174811107771388105920854739976852943755424843234202221548899214979351103529478786399

20451589252620092162166113052211258190914092095273810341317166411421204155413091250526512313028701257112314821158290178411221493744174811107771388105920854739976852943755424843234202221548899214979351103529478786399

20451589252620092162166113052211258190914092095273810341317166411421204155413091250526512313028701257112314821158290178411221493744174811107771388105920854739976852943755424843234202221548899214979351103529478786399

178913942159174419221360113720122204754122818152369818105714129461025127811061074429434010777281026933137097225015359801230622151994361811579241809404868566229294457392247348186132748811413806981423391655315

2561953672652403011681993771551812803692162602521961792762031769778322514223119011218640249142263122229167159231135276691291196581859276723622141110841291221068713184

2647824450268472464122913227141689124210281961170617026220232684817403176981942817492163381969318685149409300436121569995241380018783115851645242021634616095184036780161451625014387195861478319097536596215874517372155900667752995699380122559551730269298006130328566936891679218

2566623712258742380922058219871632023420271321130516421212422562216799171681877516890158411904518075143558981408951517491891330718254112191595540881565915571177096493155281580213958188051420718313518592515603497069635657644151335531368021759266702766457657125448260909688398931

6375287255546935824436587742724176548403763715292753695104033722281591417301380370294365814743944822456073242224363725181283122061152181851499615312455249223244302336211174252190

5785076725166485544246257252553825977503453424992613394783833432201501403278349351283343774403734372234943092144163384831192871851082081761379013812350236198233280311201164231174

5921533845281933491735579031293014303220298901423311911224342145221131582034359252171091261515132511222510102116

2490823116250062313521234213071582122664261921097615927204262460516333167091812616563153861844917591138908712389211467988171283117818108531552639891505915109171266195147921540213689182841375217672501788615333484067255439625150205350353621128966674163617281121437995887785208712

1356126315741304126310877681252164558792613421678906898104690183410309147934462812828540804856596892220974768104242492873955310297661174270569374217334336351317299196107560465316528683382515455425

87372586069776666754380688139059972891559259561166849760567448228413345583314255723635301224995005652575215283895644666071722951971722601872111882169064344236203267530236292318294

1840190619901818162918591328186619598901176133516261432132712911535130813571643102672525891155623964169793112843311029128912944431082127111171564132912413545853444055013785144134502561656833865014721006628736613753

22001783206119841892175011451901245599813851878227113711400166114151252170215961201811373213227861229154591013063681455132616266091462127612461604117116355129305304086815305844665034011887567166226631015776678881769

76167900690466345854628248736748725931044872537561204924498951984955473655055214405026518331423024743360556531264950113836624509472915983602490245975780402247091349223012341636196014711852156614921047615265815911745160036212487288824072909

13211047133712961201112076012281442593802111513877638209037847529838688114882028730511725842599698194976713922353837785510966732967250505323228385318344262289227117492364405401608506443508438

67645972665862656839059184029641158681345351656549144250337942720114434052764065583043801154894084821844833784484693395251422821991401501711791061316962240225175258403214183239210

181819261662171215751893118616671775830120314441714113913961327132411731379141898567126891213638100216457341195310103711691280385109212531238138010261240360622348379446360428419399253163611530510529790650835594707

353204287339247255185294328146161308385169208257190179217202160104616206105151182145174612181462259822719816423717825471106995612492665276382110213779113139939310584

7125286155375414973415736802904464986523373744443284084293754051651204398200297287258299122394379389154378410331368361432128239152961561371251011407572258209137242287178203206204

591550724590556514433599776259391544687414457465366362492449407237116538225236747931437411143136248819645937626741433951713029414612018012115014215810567241220217230304214220212167

9749131219892916960599102911354807089541259803762873732641826804534313194064239563379047660413780559182429576462958274359585523044126118526826124721222617371371381318358500338365350292

3042843192152742271503363021471663003801671842141151631961481689667718310015412710913636221165207932071887816911922349194794668100605661641393107103110149858610850

3515305038363654322431242653307838381644219632633789242822632680228422442644245019801274653619861269188122241625236161923382309254284921772026184623961888264585111798306501005774938598734410318125988784612321708930113612811199

759578892805670504467696877322485756929435520591475395581457461246182544131743344936334310553147551125757344332360142164814939021710220720320212217613269298287184278400278204243211

121681431201319856981665777162177908812052868681934138378719666726418126681015312976478583123407864152033411728208516340746554456729

121681431201319856981665777162177908812052868681934138378719666726418126681015312976478583123407864152033411728208516340746554456729

81273897383285572757179010644016057811226604530653602497648610585319271752533549352936649711468752469428761744842978157678418037027120325224323616616812180285275284349488306272328287

81273897383285572757179010644016057811226604530653602497648610585319271752533549352936649711468752469428761744842978157678418037027120325224323616616812180285275284349488306272328287

81273897383285572757179010644016057811226604530653602497648610585319271752533549352936649711468752469428761744842978157678418037027120325224323616616812180285275284349488306272328287

15793981911224142958014744731174674166883511509271444550133494869242110387271154893101771012257141276290107676613896401110093127948512736688257257392937409921013682578787770645150316172712512469673138977379209600471039020275897684161106349611613041406692955392834023303064332198149548391528546359271294422099425143352109221734153006496928246421402794236454800074616175672145481432743810

15793981911224142958014744731174674166883511509271444550133494869242110387271154893101771012257141276290107676613896401110093127948512736688257257392937409921013682578787770645150316172712512469673138977379209600471039020275897684161106349611613041406692955392834023303064332198149548391528546359271294422099425143352109221734153006496928246421402794236454800074616175672145481432743810

5642775590685050944350264183164406723281354969744421992140873139453569123732812948563096083108683189312913403552533303842905311949773617082854801572542253593365502454193404148414932999130081833886612402927403833140228474840885129138131914288698126950579858679113498587438112695950359769165124482801722699999710730383782247487141729168421140998182729

5642775590685050944350264183164406723281354969744421992140873139453569123732812948563096083108683189312913403552533303842905311949773617082854801572542253593365502454193404148414932999130081833886612402927403833140228474840885129138131914288698126950579858679113498587438112695950359769165124482801722699999710730383782247487141729168421140998182729

94058012746038456409689266809721162566769226859442820657445600676515733225573144890023918937718062102967077145686937789228249398651661328112368454339220550722911178274381058576672168923566796163676433561320583594836863018350879360436187124608122002451828287780429169139317016766629213231638823894514715698140299975127356277930132385516376453465478207316405535512

94058012746038456409689266809721162566769226859442820657445600676515733225573144890023918937718062102967077145686937789228249398651661328112368454339220550722911178274381058576672168923566796163676433561320583594836863018350879360436187124608122002451828287780429169139317016766629213231638823894514715698140299975127356277930132385516376453465478207316405535512

1330115317451231145210861102139617306779061341174379090710807028041102989915544264188061180889675194625311627191099421119988872510717601264346578421229289373378264290147136545604352572766417418550404

1330115317451231145210861102139617306779061341174379090710807028041102989915544264188061180889675194625311627191099421119988872510717601264346578421229289373378264290147136545604352572766417418550404

6847172926713866283168620598014934582354644713008344029585286212337128433554287338354441035051146806374232618778659391372662333746450664113545768119904620939596515291794345799423513931758071422414836612639195101111212076170471451915761127071413687896071222511649315799172353336219248234632166423953

1603715946172791366614619130261053215069164928269104561347416064802696321050088738634112431010885615585224718874566278259245774394612717110459316119914136111961009572971254994071230632575310309922513458404836612376307618011343557549993428504481773959474150745109

437274706044787414154611939624329955698139080181082777037983377452391928168263452466329850327773099223733174454572525422178842201630178272233084878802915824901337731160128890266742680938517277032931176901159065818211113228636996986849421597039471364892461052998602074212467155661380315962

8707992093207750788271515818103048899370658037071831451835555602848185619649157065129315710463484130773905564361695459139360065379576522065713558252117005513167491692261014321614226718352131164716391018781302822481842233144432822315627872882

474034745715645953144710311943845891197433324887741929173483388319832390324232072870972168613642209435032822171521726133879254741701446364225541427265622984439113623322226741868129811337491047518379188819711410248020831316163618151212

474034745715645953144710311943845891197433324887741929173483388319832390324232072870972168613642209435032822171521726133879254741701446364225541427265622984439113623322226741868129811337491047518379188819711410248020831316163618151212

127211221212563114111053610530793011422129095316775598561243683818292910385637753897882566802422318527761143756382858953747555192876537398819631367242761464149260700988902409376826022210369928712926246126151556995430029472867339962044200426743274261

127211221212563114111053610530793011422129095316775598561243683818292910385637753897882566802422318527761143756382858953747555192876537398819631367242761464149260700988902409376826022210369928712926246126151556995430029472867339962044200426743274261

1130011177111681020492499512709810133112624740686585871056875677438811478567033797674106015379215441687938365592788948416910173166276547728326836230696758798414625177602059322621742028322224332643225323291387888376224302528286055443642383837773926

2456248624982136209818291485229425961106166620812657159516331937163516031683154213846974364150091513381690106712923651552156916296431508142610541881147018634798115864215665575754204852992129205955717591251818789915697

380944743814367030943173251834243749154922782520326924612347263525792381268925622018142236842185113817272612161926915611997218024988081889265721742922212124016349265686961203752888736803529303121471580978419491336148711901562

2557226123982281218922201468234025251054154921132271165617041633186414621855152113988203486153986512351669110114343871606144516315911447152312601931152118125177664834736955885396505053071968465485606271122779796790855

58748165753251451244552967325733046870434835041130932345740833917012023582403174172493708444229440920945236529831027244913920914480185160156821297142218205153229298199154236173

18911475180115851354177811821546171977410421405166715071404149814691264129213778766832705129767897515018051123334103010591116432934996109313708671235290514393358573376485365407181135564367435461924510612646639

89466088379277168654879810233816038121213542586635536491710618508289193448537048846233545115461753259530064943144360345969425536726713523725620715917510675336351235344446292242401238

89466088379277168654879810233816038121213542586635536491710618508289193448537048846233545115461753259530064943144360345969425536726713523725620715917510675336351235344446292242401238

5273755124155163322844916241952874576552722683541712292922282791421152247169302238198194434093193181533632169224329943695175161472401827649111633220216610419521426618714997

5273755124155163322844916241952874576552722683541712292922282791421152247169302238198194434093193181533632169224329943695175161472401827649111633220216610419521426618714997

72597393661759475559665141815765696332526243452270394985101031424847681526350264822422527665268502629084040440526355014105838014454639017723690383828665170428881191179227110121367225411531716139316381080576273821751750119232192127207121492472

72597393661759475559665141815765696332526243452270394985101031424847681526350264822422527665268502629084040440526355014105838014454639017723690383828665170428881191179227110121367225411531716139316381080576273821751750119232192127207121492472

72597393661759475559665141815765696332526243452270394985101031424847681526350264822422527665268502629084040440526355014105838014454639017723690383828665170428881191179227110121367225411531716139316381080576273821751750119232192127207121492472

72597393661759475559665141815765696332526243452270394985101031424847681526350264822422527665268502629084040440526355014105838014454639017723690383828665170428881191179227110121367225411531716139316381080576273821751750119232192127207121492472

72597393661759475559665141815765696332526243452270394985101031424847681526350264822422527665268502629084040440526355014105838014454639017723690383828665170428881191179227110121367225411531716139316381080576273821751750119232192127207121492472

264425162960331529232943175721253586150424202163340523212504270124812002210325931878115541323012142420951886992215645720751809227579021191617121123281914215950313126055541039689740635725482243142114039678931502107211021112940

264425162960331529232943175721253586150424202163340523212504270124812002210325931878115541323012142420951886992215645720751809227579021191617121123281914215950313126055541039689740635725482243142114039678931502107211021112940

1263104714241630134913767689801667711111710181635113311991261120492710031239869606204013926549909104491018208982886107535810447405601141955101924260627926349732535831236924099639630431457725483554553447

219184238238261246118169280134162174296167201203174146151261149107445227108165140822083920814919271200132113176162189371114845906051676357911610360701268411110665

219184238238261246118169280134162174296167201203174146151261149107445227108165140822083920814919271200132113176162189371114845906051676357911610360701268411110665

1044863118613921088113065081113875779558441339966998105810307818529787204991595116554682577036781016977473788328784460844796579383020549523121840726530724530618390523527371387599399443447382

1044863118613921088113065081113875779558441339966998105810307818529787204991595116554682577036781016977473788328784460844796579383020549523121840726530724530618390523527371387599399443447382

1381146915361685157415679891145191979313031145177011881305144012771075110013541009549209216207701105976543113824910939231200432107587765111879591140261706326291542364382323356242144782773536436777589548559493

1381146915361685157415679891145191979313031145177011881305144012771075110013541009549209216207701105976543113824910939231200432107587765111879591140261706326291542364382323356242144782773536436777589548559493

58548964569563958242548381231556651874951459561857042543756045623290270630248141523740712245440353319044837227744339349210131614812323415817514716811357306324235194302260219237173

62777165779874181444752786436256347278452152861854950850863942024890672436248141925158210249641452518748337928562144749912728613213723316716313515610173391341229185361257250237263

16920923419219417111713524311617415523715318220415814215515513369284190106143142551492514310614255144126891231191493310446317539444132281485108725711472798557

28915301770921170376323025181630187208947711026971837876222333011321161876394160553028653102266608214310220739162059947172611420843432323165179324910016151777322252469510158421919749141268973846617877165588941260104149071314419014698131149426121678110240721985719143580612155053648938064534136955050827988413973016428065500966061624539091734689767647657796285763608811164406750531793941836575799816

9767272649632162375127624284542613251175486685942995381740564222444431001084971684603678737784786375672642041646714804164562259224426411104763318359695661642277044215520512645267147323593340083139621461161111432012282106614282663150311489686958510536111086313052498554967051186901095651566371076503531323423311385914693956335347270128478226504191820142794

66115450677163080258605836883841993440598296639564597045136479277605865867961720625364892355763606304796029418801987656733059522784656921805496231180746655408735126618840467743734526838516984359451248145553645915792154222636015206215301766620472137465773343343596124494197893543425394219212833124645

2268161226542988203924081546154431071407228518572992225422422201228817412132221017181030274227651203185917787681707379164114001901683158813798781754168219094681357511517948466757600745452165107412058116891208906721992966

20721484245328221806220714081402286412382127168927122082204619552132156219012065149596824652515110116861659697153533914601261171462214471259823163215291709412123044347485843668255166839915695110626936071101838677893911

196128201166233201138142243169158168280172196246156179231145223622772501021731197117240181139187611411205512215320056127684390307549775391231431188210768449955

231517673077337925332670154716463249151525192173339824232455248326111990205225481762116137462938135921591869798210141818981703204372920311451102221681633226050113676005389205257876567565812021328145993276613949608021045927

132711141839178414441523886963183584714041218191213691355139114381134111514651050651220216687741170972443116024810909391232459120187958811789481331294803356297528303436340400312126750834555443832532488595566

9886531238159510891147661683141466811159551486105411001092117385693710837125101544127058598989735594117080876481127083057243499068592920756424424139222235131635626976578625377323562428314450361

39674269124448049283367864141625162265135098023258365123107147697368443526536895385692929733919364422874518027472774556919821311982859912827299376793282542499831219114322798822160163453187926616314438317216699164894615591856512788103871193680683162202392079514119113012060914801123011655714634

12538871811188512341542904850188485513991110183614161272141213911249120613169876661836182178611421064427107124510188281061444978767686113010091065372953335343592361489444496300118810751532388732560467600549

134310201436152512311384776838156274410681041153512021042119211578809721253878541160114756539467914329382059968059974089307235668869291003234719306280458215393329386255102617699466425702425367501453

67684738848193296473716542764603883139346633539783166468611963997035512758366631502134058567781634845448527322845215121647764027528519824746377428385831457954581416366816441609274613852191190119021341484357035562475192634282518213528852665

1651999183821451491177511591123220996114571285203315751476153617281185143515221217757196019538311336129554112863121201114212894471109963670134410931346337994350338639369556453494372148795770598504903612493669576

2865919268309143439926357295501804719099364941676425955222383397726183253562635627258208562447025720206421265833313325041406722326201769143214274815202631819622587815120225159331158522688190062257159581533565296376111566235915972608658580023101444715019100488058148441068688391190210391

570032514375565843925440327036665766315841413764574243874679486343683885434849943706201457046019236939633171166435371059337829373612133130562844201138163284330310102488112511441613991142211971576951434259323241651128026631733164024291715

570032514375565843925440327036665766315841413764574243874679486343683885434849943706201457046019236939633171166435371059337829373612133130562844201138163284330310102488112511441613991142211971576951434259323241651128026631733164024291715

10824779312115128671010711435684566661363064379939828213664964893889704959378988490925274754668146851305452608452737534857565166980736648833132248344621940748311713886432089537423592327358521353119246628462064981539660883563334657423832331740063788

9259682897871079984979722580557011138654728237697911483800179328138782366497065757362453782124561116043847065605929496376141668295625703426946969530333717007599172361767445219841936300217602602204923851749833449551952964284948973210281533513140

156596523282068161017131040965224496517021303218116471456156617701249142516791230886222918948761387131653611892531244102312975301375916703130411471407322922375391583375517417461315148901893599497845622502655648

53343732492960834726546936234024623437894308421757845030465055745107411248225184455425186044622230474647377722634776148934113187416014413767329225083770324136902170420420331950370325242657236026131630829370440903418240738183162314033022615

48283441431755284238498033363738563334613879384251524510416251254604370843114751425422745486563828054225342821154322138330552923377512993413306223333453290733232060390219251816346423782494223724131527774345738023212225635352964300930592404

50629161255548848928728660132842937563252048844950340451143330024455858424242234914845410635626438514235423017531733436711030210813423914616312320010355247288206151283198131243211

910612219896144607432504367871192487755556245352165721344923625403917361005694624017620058723066313136593118590951743534514299195008330906686751326636609364230475193715463022255340276938299210883552727664369163883183823376568271437977205503068902950711151027219481499971609189913616593904295401998997789812244536546311836103084204583163489118149

320475168523319323212226443248183922411035273415313050783065588111314359995125112654061131793573484503682524042078499865440352161791580242944211491391722102568976193751944027816684115622281841271127698192601782913695364548072143622973327390472493788010532776209302112306122969864177163859311454111164906752081

254021751846331422462433155518722761162933203446717943444605362827143815470041221763161123873586150318761746908310610641686166016965401298153711432035151215035691414563671102156995592614299742611193975809628141297993217421371

295074951021473290072001840815168372223832512399023017583031128039524316555079062617781104653535334456682482821902198254416482125931429941068194031300921791967912176891778026120630114324266471156825663177481632613126350407509136912871226821462943695410389875235276011113113219055170883718110475101844732550710

59873449508451355037529887492799624532894517392877194957465345475524376552793869419123376303542320014062341917714057137835952897400715813531323615693258357940491081242811381184181711431588119114491045412264224741782136028461873130022051894

138531852712181771489023010311716917910316020211612412812713751290241102130121611274815383143451651003694951323789444353343824503115104108716610945417250

3041993652863802795914196354469505376142973781648134173592040523965042334919022417812389562734121523083232177124200210235712017362104136126923408822162208135168242115100284145

5545319745344578443948422687251356612717389533836111411736773864506729064231333738151636559048331709370831201587303570331012599363414533134295914092964327436829732138102110791660973142410751059926375237621581576112624951713115918491699

8518851468731030953792693270401247353730357825315583478936397416723816372016565184821437091180377217501117679469109477774853962660394456153005085505791973831706762239241798182434282660814531414841361621219881626023328102377946636737098221528265695739347924957743493480992351913007246961776215621410523013812261855844501867097773855399

8487731458221019153769003251621232213641057626415357678049387276610616177316387183003435197178641215822116068465146476171847532647134438082989635494011961841695832218371791472415192646444453414565354961208641619753310472365576554836837215818221095198342064926543081475772299512727245541768535559710461513519260858839441859347663654665

31121051118023691878151489319892258887101411321947178018181894173616791611396316036431326180715451178119910932087671190914377802766661124627176312378192615714465415863124125225242801427686176152939975067751102734

72935485381182379756450595554669577511769268499067107168628196083301168110138862757728061023165342859625658640723447753565019341223014426917726118927919968412482291271445263206386241

28012031228028828823321133819722326740227024029623223826932021311837834710019718095199972421542129321413390158172218751418160896190621027426126149101861471075713377

4492345415315355093312946173494725087746566096104784785934993952127907542884303971854111344112743841633722741443193634321182711498418011617112717712542286333190185298156149253164

1573315711104201145010081158198936162781503486911039510806175871324614240120471072211080135091251585495996109331538057099137600952971036739027571830491313236729083675700990983747255249964242355298447232694385537804684290511295650538743742826658142424170327997455

642316271652649198861424302811153474172011254261615561233113351222115733114211010245652948141422

1566915688104041142310065158148910162291501586831038710800175731322214210120191071111065134751244185325976109221535556679131599352821031138907538829391283236728783625688988783637250249264212352297347192673384537704660290011235645538543652822657342414166327857433

4231182418363518262625901611259032311499211424314361323829843069262427083254481823931084242830532228201519381774303910352316206014915211178170110072416189515704641032620689860518714760907539214126810459045611516802108212941079

213881161941771221551542127715921635221928323317418730528814876160208214119896122887163125813472106701271031032365343849405435905412687073489850687884

24461123116522561588172798616331911965134315042893214318411812167417342061304514177001450190112901203127113561904637140813729303207161099620156312401006292660370455532323445526566338117831644545339877468670860642

15726135551068861741470803110845761271111168768601024776787888148582830881894472469357835790731174556348016739049631772655246114930721619627915521519925114785369331286174541284344356353

47137323123412348146371666394732476325236078133823308692941742205390683943939984406132635849665356923310415793401856087621465314972878114367275287610263072881343574116182671123973239923423238692304411473856463108461289719702816514783141151325674033510193221899412900123622245418316110501608119859

47137323123412348146371666394732476325236078133823308692941742205390683943939984406132635849665356923310415793401856087621465314972878114367275287610263072881343574116182671123973239923423238692304411473856463108461289719702816514783141151325674033510193221899412900123622245418316110501608119859

47137323123412348146371666394732476325236078133823308692941742205390683943939984406132635849665356923310415793401856087621465314972878114367275287610263072881343574116182671123973239923423238692304411473856463108461289719702816514783141151325674033510193221899412900123622245418316110501608119859

25715530033634328924519643624724026139720925037128722426823222910248735613622331185201652621642639019516169171239261971601035698727065876319151237122901681046416197

864101489481291411577044351135649689130611956417519677746307236316673091460107235281559322658215375448678726674856519848366574826255921717843625930214925712069543620406392521319210388295

3249242430761062844541453629011560790748232580277637663721468039164894265231723518219214133205152605208379924522276298557723281785278518492815216717422496304023615758365301268414121341029383742722763840303215316971376349018365146112016555700

142049515761253127818091356366232724781721153719261531192027242321114528962076664749140311304661097299723510451602126108218243886165223571108138918077056843882664442816337186632511065648626943261007427378615434

1407785139523891302155310269682230130210991056160812641077135514059391254120698652114711652794118910486009252641006837113152910367744891000105410604129713113404853024754845472921748587504713287255083896551181

230851384213791192041641826887157381806628383135251254011683173631912017505170651762610257270581537717922673416151239248898134671153854271036437969905150512316947121153010411146341669521461137944206100875629512910831349655285260531031451673807178715413456697446666397173856781

16855135971309113524124571771610506109321836310799120001079815950125821325613586133061051114294126521044459651600817482561110907984255181142625959926940813615378497719373650312279108441041032987472293027875274272639383167362926921166698269574418317084535146491852225371

691433774523306647230211914677299811091219242260516163822192314217917356118213418135114735228295615523311676220930868453101494628231707211507301464172479905568616368121939618471033845369528479704932610967086130738856331013173528474186379444603168638267456917081468816973363912829779245653944013256313934161287422806831580880

691433774523306647230211914677299811091219242260516163822192314217917356118213418135114735228295615523311676220930868453101494628231707211507301464172479905568616368121939618471033845369528479704932610967086130738856331013173528474186379444603168638267456917081468816973363912829779245653944013256313934161287422806831580880

1982614861350506163431400274532452434695399091364642098273763087031994398993511350181248162543234616213741492618437364993979929680454391197130415555814987274371857362671650812800178403158317076173706071137538679109821466464291145414134206545399227811781149011137869321346210582106041475113987

1982614861350506163431400274532452434695399091364642098273763087031994398993511350181248162543234616213741492618437364993979929680454391197130415555814987274371857362671650812800178403158317076173706071137538679109821466464291145414134206545399227811781149011137869321346210582106041475113987

49317228841980164106688774649846565858722918614247992179825114803142691150140141452112239232775130417913301746924707986568443861342221109311167372025514371513326616381468607594735122222125398513309492469915568557839472921360433292653504953974203165771667334610432824010551661434163832635186992587950705316765356466893

99853067792592703714044189497090794333861730140247208794868137426193961168436152231952229231294104252502913907141321325620968291733364117341541165965574112649579209793423100848808976335211254186585276101031148565711141566561000663931086704111349111155074394410133240075305175

1728120974272277336271985235326801039612144019498526607351493865722413841214872807720313754212516124345612959114232559903521291637292416478011560104115182608397219352426772414022225735657364918709614214961106109881447139695013127221323071862599

179742432112205456123806361611318487542511867522442033307138092739144123293153957124561182854785714253690690932017514707160363840715752416735780800044859575316542554171318864370511248569243381420839735116746537077530859612273471923425988347061511070106291071611524133277593

69711034273198269182296111380725812749891970255688909737663581955720651036452091065346230730790620058817406597848650746764350915100612145345109036451085552507686624019729197475093316232180269271729832370152021268519930603925643368372498411882457134600182147754798927781510688311756744782

186393762472914305828942102276750001374419324674361350829293301247125792733362223862038288629831701266626928972282470244821242182103425211341196622821642341863014694838771097776907521870631189101420161484101614441102674873929

10796489720665342011360085957910175871979757831784915502149851725819030161582390913654123582183791678182852715734101681319320244660917452281673241521665902904664238179305152448515953229376004292940436641265057306015676831191257651251424735260057726427501770815815

179852314359011220337126930810454211686906871564101483516369608625991201358990834156457313608031243575110179413607069024821271202127400111294336599021263210152978048395210801938588744528931083862262198848357101843312050113836009410799654717223331392483996450934779245286570443254042315630611947265686440163344948354572305217121816645285576387424724266553755341542450514107560359556283

367523145216603739234121248624835295228042123376527739753830393541773051307338302687191456544748230833553242122630376103159238632621241326421441625298826633568766211210689091432864134710201149775305203223791434128220331442113215561321

276817843835450329433076182218463922167530412493387229602836300731412249230528162009140143743541173024792378933226445123461826244092023851616118423241948256958015388126641066642981757863570236153417541035992155810698671147984

86859011171337842976586558120055585875111409108318499156517147836613731260104950374768226872415269252869229369250937970463474219848425218933818528222323518772470522304277436320224347304

1900119427183166210121001236128827221120218317422732205020052158222615981591203313481028311424921227173216966651540299165412981748627169311078051620131418273821054560475728457699534628383164106412327317151122749643800680

9075301381153498010456646371373605117188314051015994928103680276810146785131280120757887686429377315981356082232187952844166471599918657425624536622236626328620569498625399290475373265409337

9075301381153498010456646371373605117188314051015994928103680276810146785131280120757887686429377315981356082232187952844166471599918657425624536622236626328620569498625399290475373265409337

1755277140492711681071213451100115316394158440149875441581204837076115874295413915082031319312120236910591911317332869777123565412335461098485640560119704514769044607671043762827921439393105029225503681398999081911673213705019057789396697050291357341967001897419236422547305243011306284596608256291426823334394342136296256118211622367549773409061252403731406527079500795543106541193

5338361376016394547857253342334172093196572945867488534949765094524342674782527036392845764861022751448037261767426794542743485456218004430316222414792379252031084296013191267184212291761132816091182433273632961989168829701895167121701719

4302118084507045513462896993265154837964924845854264394095473861721224690292521272168468955384065172155043221513543576231293791838015515116111513110849285499219253332166103225112

3909272550294280362840122293236148452182381931545103369233823376360029363379361625382015476440751794298025471224291765728322350308912222942223816563570266735187181949859925127684012289071129832305183521431360110420431336127114651214

99967717641664114611627036911665688139594915891165111011331217892994110771565816601337665979907375882193904729956363984602434868768106223763227726241123837230634924279616654410331595393297480393

2534641573441606281748891470102489471143851240942442331150581626581278282367952103171873761415242265121103871883611564621825976568817121123662971610157246137339540081250552806011238411299415022955794112557122319914201442701419531273113350472865345024237779028335295661939116466943659417531956888393950197323028215271126512087494466816

363018353547318736523474229919424072206128082679434127582687320926562416255822832585998534938651305280519641132216343729691923282412232727225182621662232289066617317786229707097656147516272681799208011361206193610727511341858

42022183317942153155343024611573424420943192276644493675341533975045282732803395289213023714371414162773346910682272589263523322830110123412240105124202360257862315027106791186622119212638706582901596168310898031830122394113181341

46224332363151432925324013760922187198143990720584309912611341509343732980326273403142110132269288023094414669345143579811399289952190510392237785679228932238628651128302384024896141482974224701270276540150716445815614752801810942803084827240308316001145811085363022067612757109961261313763

634333264651441543965837262729595468304942963549582742404003423742893077411835064544169855245526171538282906146134508873799324938771604335329651513329435203619942222991310681665882138110101081951668251022871944108126721607143718661514

2670814824180031812318267196811159413126210351402018356137922548916846166991597416441107921557613465151797393203782334058151502997726759129372869125881322313570660814875124186651136601414416723355378643280331861933338506834264001380919381084679615529361498735852533360496826

60984264598857066377650439373351744036249958459812485796914742135101943267269524850543162329788566752735488437022137418796645283403519325284554357319994225508548771133340229023743570320655558162834261199502316445043460161031832233169633592435

837085506743826656003690458583267232289572078259794756138976714628323530914316149316534277733855065768853194693902941304126234703061967124933910583863626529937455941580631042361942224347948453523584984451748475421179229712937918387126321302812128578623819172211219870962155723235157832279920078

2955616623192362191820267359531982617211267692180124926153943058124714444251709624605134952471317326266557586242543204173271875119919876215509341614028161051760365491371718009806918288149271544341619757421347479536425477554988527742581838128479193565336351015974345587127477307

189107188217251227121125272121197201324246233245177168198236154764113421221781296518978207991406717312349120139170311175445595461347146121061538499138805810473

46806258793049618583213407764922610410986294821725203731976040328322614045525969203881550822740282872647510168301538402427153329731160697392146547531247220337299477478159351965034871224072949318135741013708766582079252420855105491970756783146230002427682516856101281563386261274812621

29722353398248233361347421792086435818603317274141783115314431743519257727103374220016014767396420122779276010092541535245320982805994240018051398255223302681656151878668612096441004870954632286158118881170107517011168111013181133

29722353398248233361347421792086435818603317274141783115314431743519257727103374220016014767396420122779276010092541535245320982805994240018051398255223302681656151878668612096441004870954632286158118881170107517011168111013181133

249811952750217326192688148712763088154719072054313818342205233819651587174517031641711448529511236197812757191560374214014172200935200514927621549166724014931469613427641570697467578468192136117799199491432808550972636

87438483563790083050545010804926196691018621685798594568532503513188135699040567240522250313772247674429663951125455255778015144519313421017217816321315660484595305282446226169323235

16248111915153617191858982826200810551288138521201213152015401371101912131200112852331291961831130687049710572371418941145663913669815089971110162134210244202934313985193043653121328771184614667986582381649401

4408582846413421923382203117394328562406402528884435952428083008172651594174863123963085603083483109062383892908352779592859101311564026414162021385702845982019601176352590176205823803524907029711710754624923224370513672929028824809526653776569175693721707578313133477276984327261387818689893072719161217743711474995022198484122361105651137078120551

4234942740943246483219932958974150202307202425794240562334152870402525583987542990102943932940562977622276402786152653642750681252033805773986351311062723281931341127162472485940322571823848428322110249523670823395013060427709423764025307773481167566688147268312628174075941946938983792661012944918345916866610945290217189664116984101208131172115484

173641054717544162271584217836992010309195399393137771260118732133861416714292131441074912220125951084259532206417567746412270882649191176926551231710586138965051125249755612513194104551346030888127335631005053320142383224402628881278815387715297480588205377444359065067

4416628106326643250332020372302367227780413532950830368245684396630542310443090130883233132947126193288671310438398412741393628032198581297125868585523357264652913811396283232512211755278442595527749709817042675772901351175149522716996336827326919077162831181678802060013279117591342712971

2757416896191262019218504217981356315870236371984718305142142711718590182611777517707129531795514702178407910219232436187471687212347762815674348213811168241679070901749115169658215755158701692241641020139694233823143615503439961454101200411238941771024477121858523726684758296

158381084912749116141277114636965311458168199132114849806158281136812183124821263699091099910995104434943153061598648671062171395128975322818827918211760405010154948750331157895881014828116481261329125080301638142648332226071203742663774424315079484508432347224474

754361789697745796456452897529579548102158460064454045151749658425111699273225393722154419271945958825667846614051149767912336017514520013720512216611962413489290253467248170230201

3982266971517376424244762938314759372559548939156314515044234527525939214047509230632710525350792668417440601478354170732192879338715223442219819463525289442038652082108311641797988156913041532998317218926071694126121991762136017091558

3982266971517376424244762938314759372559548939156314515044234527525939214047509230632710525350792668417440601478354170732192879338715223442219819463525289442038652082108311641797988156913041532998317218926071694126121991762136017091558

6634637347363846503544403125546023253180834263965156262971274394493333874916334351074590643697264037183214594450184975603414603152903439485195861407213397572385901831004537211099132538254371304601051183933509083797923246146390713522932824067330316877676356110069250606832981660931405121199421177474281919568215709315430065131298607201968246741188989245289

2638141126702356273328941640165229811547215819763446207119192395198818701795172217758124493315810591991127365415793842160145121888462207159976216651729223254213346854698926096834465804862271437177410129741538782585940645

24503236162122619748198322496414907125702485712805190991657324143197591921019784175131584017972210161539510371267132420390961649810716779617941392515108145352048154591503515603889620512143831491537009248451340426943413261954780500044951838110531046673255126133447764861281317705

2734651681621283741264411191952766131265662064532334381327851519561173721843721318101618321476621450831234661382631085011226975580612931023329553434129723762204858612675836574968872014381577316577413140613042010835820838315671512311129102777893898842562100615295895061335990475984068819409675457234058127279837788467433472086473560087

6621417339045853405842648368392788366369965446387724313945688693877580458223500238235155739674799509339722541603464658765244674351851741616315647622275442603748537738741629173437513075742102385494648599772684974769964259931143517753195051309318020583233643211462525962183921729567383732460127824

433491845637326233523849539046266723804051452262572248727801353762137723929345531598522050157731933023628572353926439241411124441116901151023983584425443169153107310671275382191213154293072016622933737715433711739864489707859554279727445152322181701707211160967024473843412458118786485

106506609120039153110261164583656144130877160869687971317586608392109906643788968898006787831401758613199473088464131322778501715858462751006132787875760227877414653084162087510827331825215324362435178323991871937631566353531333675763055287838672443

2426443430921371981328069479017877161097978381243657780119191411518016173417562618555910431913415594605196847243892130365185278772741430423382411474082819496962191343919661383159031200833249491376741589051599003296881142216431420518330151484447221109521280198245973729439984767212254575192766047886118241345758493313662774837140100

101156756102089122950610612848662291266876538192840912339836992319765961870398543794069814013140741118951657416527533057684369874706054827032487729625840418004763979052047531822612141359425382860223226371908802474857794010373758393297286138703332

303320653149281126713079182016913329203422352335343523732511286229601975244823352025128442053253162023511616106020301017219616352433985215417411259215921412292676180964869011277338976767585332311470172213269291822103488111281121

4623454424835245262543335202883334195443193424644212843633403801429495002423692101482719142724242614044032813933832542380231133881541011251161159434246330208217258161126224152

45683339451641714134485151203072617539943998387958854189464243264600336843273970311019885428496721483168244515114081206930882959362713843447286319634055374834168312165971100616481120135110971277936379204722651576178025721541140217911610

20521007210116572177215612921133264413371626177624751488173621131637141214051295146659934922469115515281004586130252117591218178473916881326680145214251774460111350935766558448734348734515898514629008111187561452727449

32686228504484945690323013562122198229334419019009347962843544844333543186732742346632637428449332422450516370429523969018057283592633711120265485307241652139827235994725448190971382927093223362844867431636477987743121907345105928632101116810268817444188871143397171766912286101351286911688

20511267421048752448226814581561353413393188219235922661238124832853210020012713161914383140294715242151230977219923181905147617268062043111810381881158923905241383627623963557921788838515207134014899787551177865622926956

899260061354313595934510212605968471300951961041478721302810274936490821129474658528974570435181118641079455028336823929597548151469256205779329007232521940857988642682131947470221352287370021173260261431312009743495453923462274347503649295136943444

127210201900222315171386885941195280215971328198015191384145714591101114614489647501954176783212411128471113821311498961199427118379359912369711313292695353350525314485370490288129750897529493723505429572457

333824975707627439423626260127245463202045703625530741923847378644783561339443272425209448574527248334003761150833136002928243831021145296921501709316127203635780184688110281548856131011551261770334201421271301104219181541121515051453

17033120601948918723150491812911195108602023296521502713418209371470814891159341457912147133801500912454690721137196557716132311090054101255726621125810383134154669120219817639812827106301289732007738380234555454350146163705439132281275838689825163468491015726491861725378

2948211599591701882365221392842544791059451167182576241161821649121522132385392728151595771501062841001297342308682177682169058657615993619301263496184103186060519311396613741714177712722118134758610118455134040115900207579157418141593338508270039137571331005254112777357599126036553928190568970280150564273326799139967616740610537375191

213385922301646216224161196114623551162154217102415151215801893154911831385131613126093551210290914759255591298296180811801811680168511944691171135818694161110475334481466461320442340165115911707627671199622450687404

290886158386166188233010135335250340103511114634252845114072161757148440234052269071155871146006276925127323228322214529214213851531538031884856160318103218266050838136701366441386201249501780185739711546113187611488520519815483613832833126807483824356435995764025776329588685945753235187248759878014550353201796774948516637110397474196

18027141770186617871723123893824249481613206320722232212622075626122811611923138081425822425984159624755341662477134910911518533130997054612101224139630884241936446840456772446635316794596663048311661288585712591

9147051244129910399826596561323569110884414839649029469467308259837174961732122654584068135082916789060892631684967939477462998221051822920433123331723926317391547635357374614368343387309

9147051244129910399826596561323569110884414839649029469467308259837174961732122654584068135082916789060892631684967939477462998221051822920433123331723926317391547635357374614368343387309

173211831966207416061744113912332109911158414702234153915391643157712661423153111627342296188891013831147550124629912211161133752413241019706144511741439342799392324547347464396463308136875969599502843568508599610

173211831966207416061744113912332109911158414702234153915391643157712661423153111627342296188891013831147550124629912211161133752413241019706144511741439342799392324547347464396463308136875969599502843568508599610

173211831966207416061744113912332109911158414702234153915391643157712661423153111627342296188891013831147550124629912211161133752413241019706144511741439342799392324547347464396463308136875969599502843568508599610

3783927477450484774638739416262392523575483522233236820318494885935977358373702537620283883105235094270991669458215462401996731693265641172429287625329988240673309111334307132263914973307092561232353775620227957181131336081841152991381082478783164200112326613630123662105913361116721509813159

1620112187172031746715373169719657928818766917114170127011937914240142251457314346114051255513462109426485235191868475681251597704668115182512122199878133654603123389611585612435101881293531318254383729955374334545583447414730611278819092815431504988515456473362045199

2108144131322853221522111333124727721348226217423004196019492085192415681650190913851007354526111183174816076291582319173813171895669179812597571694147321764441132539474685474601532622400182118414858197561200713577878700

10846821110801105641018211261646260971187060278995814412469931891489538887271978299865573764110156121227245498126585030797534168879896443889329848065653838768277664382012011528925081913346321752917212326312028816539959063572331959803618324340843305

32472535327040502976349918621944412417962913281539062962312829503550264026062898218113684362380118362641231396024025052492211825779502475181412232464207225586761833790608122669610407928946332801607189010409741671112591312421194

366222423515389464248251543242298367476356327353336242304304306135536404182299267125238553092393071193432011422752903047021910210813672107104997423182217142119191120101136125

366222423515389464248251543242298367476356327353336242304304306135536404182299267125238553092393071193432011422752903047021910210813672107104997423182217142119191120101136125

2127215068274222976422977241911402014036290431291922352187812900421381212852209922938167411819321328158511007434160271521221718879165276931175313686174601395019419661218032128278975179991513419114455511754563250107850476768645587657847431863116391376880577198120177785683887587835

337224744957564938283996244824895010209740663288522439743752391743053029306139712701182953194747235135333300126332556763021257930721142311320151630322626153281754192610138801312800125410891211788356195423121363119419911415120415201476

6913791094129299674147248211994868757311231836787836915674643776571431120994449481576226369513967641770225367642729560761674018644023217228417826622326217254465499278262437273225338260

714250368266838673037976442543269162418066185870890366226604688268155050576863725075290611879853936725764469421275271115356344254647020435764431026935622486660631442370217481520240715012114158419841386558371946602529235839992458222227782369

658947677327779868567343418042088290395764115504854458096165648364644761555158954604294010627811133435295423721005051109252734237590920325408396626175312442556691345351016441391233513741944151819241567552345140642398217636392329207825282261

347824125778663939944135249525315382219943823388510241403977398144393227317043142900196851264811235734723534117832596262856246332661142307121091740323226123361828217699510471512914128611731197830343205022331489120819511310110915941469

119982212951397107713226658041554705111610081517110911411177121788710681097794562148313986088978213801029201882755102632993080542811397591026222637282273425250358263328228103554692424384611420423472415

119982212951397107713226658041554705111610081517110911411177121788710681097794562148313986088978213801029201882755102632993080542811397591026222637282273425250358263328228103554692424384611420423472415

119982212951397107713226658041554705111610081517110911411177121788710681097794562148313986088978213801029201882755102632993080542811397591026222637282273425250358263328228103554692424384611420423472415

119982212951397107713226658041554705111610081517110911411177121788710681097794562148313986088978213801029201882755102632993080542811397591026222637282273425250358263328228103554692424384611420423472415

119982212951397107713226658041554705111610081517110911411177121788710681097794562148313986088978213801029201882755102632993080542811397591026222637282273425250358263328228103554692424384611420423472415

179461192617494178301591119282112031274921242102601391713325190491435615543160031535012238139421446112423665719827194838040132601139657911334930731202411605137714987123049745746813512116491275734668487362935175968348348293890489531851495825383585545456192576055506867775667

179461192617494178301591119282112031274921242102601391713325190491435615543160031535012238139421446112423665719827194838040132601139657911334930731202411605137714987123049745746813512116491275734668487362935175968348348293890489531851495825383585545456192576055506867775667

179461192617494178301591119282112031274921242102601391713325190491435615543160031535012238139421446112423665719827194838040132601139657911334930731202411605137714987123049745746813512116491275734668487362935175968348348293890489531851495825383585545456192576055506867775667

179461192617494178301591119282112031274921242102601391713325190491435615543160031535012238139421446112423665719827194838040132601139657911334930731202411605137714987123049745746813512116491275734668487362935175968348348293890489531851495825383585545456192576055506867775667

263120772536280921982697154518703048134622662169302724702513238024361964242723681822118323892726109119401774902197846917281582188371417261455112821041649174251112805135849495428356626804922231166107674557312849407961034943

257415752203240219682661166714392995151719091765270221072131220524061801190320681736838287629591042192416347521741393179917501968769171114949501915172317614721134476447761428627490582411209112611877136161262780638898724

8717237999307291080574669111555064166011588238738376815807697708493781109110049364560626489519050553971823370948249262851259415940618614942718829122331515768400439272219473380358391355

2867209427643102233533181784203033631634237020452851221424502406257719052244246319051179277930671191208218459202119501178918552276706178315311145221618601927522127555357310594927687058185312401301113582362713929928601055983

9003545791928587868195265633674110721521367316686931167427576817572505988659967926111307910674963142236669553729536616152062035879692625656375478337536649590567331802439219011764277218332308181025001594755426045212992252648462963241633992662

1751842532592141811302293401211961892872412442052111481781961209634425910621120570138392401912015613813184170230219441074347280200534958281110610761801022582087781

1751842532592141811302293401211961892872412442052111481781961209634425910621120570138392401912015613813184170230219441074347280200534958281110610761801022582087781

1751842532592141811302293401211961892872412442052111481781961209634425910621120570138392401912015613813184170230219441074347280200534958281110610761801022582087781

1751842532592141811302293401211961892872412442052111481781961209634425910621120570138392401912015613813184170230219441074347280200534958281110610761801022582087781

1751842532592141811302293401211961892872412442052111481781961209634425910621120570138392401912015613813184170230219441074347280200534958281110610761801022582087781

26021030240030624218820541616028124839927126626833123822728721814445033116527326512821355268164266952551601352332332945614579761086395761066627162158114106131102919697

26021030240030624218820541616028124839927126626833123822728721814445033116527326512821355268164266952551601352332332945614579761086395761066627162158114106131102919697

26021030240030624218820541616028124839927126626833123822728721814445033116527326512821355268164266952551601352332332945614579761086395761066627162158114106131102919697

26021030240030624218820541616028124839927126626833123822728721814445033116527326512821355268164266952551601352332332945614579761086395761066627162158114106131102919697

26021030240030624218820541616028124839927126626833123822728721814445033116527326512821355268164266952551601352332332945614579761086395761066627162158114106131102919697

794591889818930901478495100760582476611716997358456325957317245833191661100845666551726459416477150083129278856730271658380822156129218331224223120325417076443627359312498287230372248

29721330127129827019818532920827424433524726326624622328123021310337133912521519697222522152052708624620810127020923469153836110879797080562713914211259159757511989

29721330127129827019818532920827424433524726326624622328123021310337133912521519697222522152052708624620810127020923469153836110879797080562713914211259159757511989

29721330127129827019818532920827424433524726326624622328123021310337133912521519697222522152052708624620810127020923469153836110879797080562713914211259159757511989

29721330127129827019818532920827424433524726326624622328123021310337133912521519697222522152052708624620810127020923469153836110879797080562713914211259159757511989

497378588547632631280310678397550522836452472579386372450494370216129066933145032116737211255629556120654235920144637457415240820912220416315213317411449304485247253339212155253159

497378588547632631280310678397550522836452472579386372450494370216129066933145032116737211255629556120654235920144637457415240820912220416315213317411449304485247253339212155253159

497378588547632631280310678397550522836452472579386372450494370216129066933145032116737211255629556120654235920144637457415240820912220416315213317411449304485247253339212155253159

497378588547632631280310678397550522836452472579386372450494370216129066933145032116737211255629556120654235920144637457415240820912220416315213317411449304485247253339212155253159

144511061315135144561395517378942311920174631002511205105541829510922130551308011519107421023511515993252182657516140732510757824464449899263812180884814025433313842840158311234297741293229728000323827044332360736392950376125781130640572365634511481644971472860144544

144511061315135144561395517378942311920174631002511205105541829510922130551308011519107421023511515993252182657516140732510757824464449899263812180884814025433313842840158311234297741293229728000323827044332360736392950376125781130640572365634511481644971472860144544

144511061315135144561395517378942311920174631002511205105541829510922130551308011519107421023511515993252182657516140732510757824464449899263812180884814025433313842840158311234297741293229728000323827044332360736392950376125781130640572365634511481644971472860144544

59305437599961764917622238524303670834024833418461084840539252335181393748025455378728035716582325774201355620954696114339164066461314894150344426035044373143051119273510741174208211821654140914211094413265123751844137130601999207321362322

59305437599961764917622238524303670834024833418461084840539252335181393748025455378728035716582325774201355620954696114339164066461314894150344426035044373143051119273510741174208211821654140914211094413265123751844137130601999207321362322

79304739827776658359105245221716799466202579558291132055517140721058226224493654835727219119797958843456071420141144763139277104374887926209157461629646753558779621729490019781383203922601788138421531390658344645193553346247622736242035951954

3589188932773168331346172265352341062792240724354340229227042805236226961964212425108776342387514112377169218211942586312020003630110737361928116430192305344063016027535287608176764617735512431356162711791114199510878411341722

39142660447241504539530027253216533130223069307962623013401339183169307427113134289012071244651722662343223341986259972340832149463113434759249716743322295338541025309010767671165130010198741240767374192826162161219224621494144520251150

42719052834750760723142850938831931571824642348729145426122532710710095412722621753072228350722561817066219112641232966874208149881141439349140724116227621315630515513422982

59143785961567963235045080942157754186753152363751658149757741822410627294034854872354401035544085332245353412645454566651243651861472111651971571879459308342237281342236235283268

59143785961567963235045080942157754186753152363751658149757741822410627294034854872354401035544085332245353412645454566651243651861472111651971571879459308342237281342236235283268

569421728612688640376441982957515079948235545436055527215165524472391062653366505424875462491752941959823252338829353348368313534115615926817016713917811054280409223238359268222265220

569421728612688640376441982957515079948235545436055527215165524472391062653366505424875462491752941959823252338829353348368313534115615926817016713917811054280409223238359268222265220

569421728612688640376441982957515079948235545436055527215165524472391062653366505424875462491752941959823252338829353348368313534115615926817016713917811054280409223238359268222265220

246188325329308253355115234655172247303482382562912724932172451701334282961561991947432024862228209287962171591122262122826311989791007370519045261141917810114711790130105

135109198193189163161105218175148142220163163196135135149166100932902241011131246011289149122174651061047014912917741736455615444424529178011750579084518675

111791271361199033904712853427658812875939513735868797040138725586706839047737987113311115542778310522462524391926945169347428445733394430

220146264188260257158197331159207182324205202218209169209191199674332391342071588615840212155213772101611332171962635013545461296467575950231071527889153104849174

24176116493525284731243073222329322437312514934318302219205451834135127933335892255202018101055172719212617151910

1961292031722112221331692841281831522511831791891771451721601745334019611617713667138351671371796415913412418416320541113404110944494749451890125596812787697264

10387139951201305570152757682151111859671599011678392011187699724610215895598599668489075138228722343933303129155596667485947484441

10387139951201305570152757682151111859671599011678392011187699724610215895598599668489075138228722343933303129155596667485947484441

3815424135402429491523294121192726283433262338222522302119740144411331611223739102111813161191243202217202127172315

3815424135402429491523294121192726283433262338222522302119740144411331611223739102111813161191243202217202127172315

3815424135402429491523294121192726283433262338222522302119740144411331611223739102111813161191243202217202127172315

3815424135402429491523294121192726283433262338222522302119740144411331611223739102111813161191243202217202127172315

3815424135402429491523294121192726283433262338222522302119740144411331611223739102111813161191243202217202127172315

114339841861758441489361228631531287324293649154690720691179931044231505501048771090011104841053058167695767102561101393886531651151205948067711418191913462499244942571212344126171114410516911090367374495291128073968631517883834069755283503522865067477323816532658415392360213085876376090971773560267020270495693864690342125

63342977770864861647045977330452148768153149656757948648048636826275965131742242222248911047241143318249929933449536654814031213116123414818414215612156369303211182293267201245211

63342977770864861647045977330452148768153149656757948648048636826275965131742242222248911047241143318249929933449536654814031213116123414818414215612156369303211182293267201245211

63342977770864861647045977330452148768153149656757948648048636826275965131742242222248911047241143318249929933449536654814031213116123414818414215612156369303211182293267201245211

63342977770864861647045977330452148768153149656757948648048636826275965131742242222248911047241143318249929933449536654814031213116123414818414215612156369303211182293267201245211

219881665243101338303111225264149161706934623134672509822271403792321424938243092303218228190002223817910107764413025903152462294617857874718826415125943156762790910849301631370310725190211993328853544212739679157829215631172825967839444492117129161556910901110401237995348906101518257

346255431408325371223279444182301267514315318318334289276302191160589362190294250128239723171942891213032091272682763076520074781497910876855233144190148128200161146123116

346255431408325371223279444182301267514315318318334289276302191160589362190294250128239723171942891213032091272682763076520074781497910876855233144190148128200161146123116

346255431408325371223279444182301267514315318318334289276302191160589362190294250128239723171942891213032091272682763076520074781497910876855233144190148128200161146123116

29926644242036635817621743619937928447429926829430925625632824313747234716427924711323449272203280119292192146258214304671587169108831078681782817619313510417711810610599

29926644242036635817621743619937928447429926829430925625632824313747234716427924711323449272203280119292192146258214304671587169108831078681782817619313510417711810610599

29926644242036635817621743619937928447429926829430925625632824313747234716427924711323449272203280119292192146258214304671587169108831078681782817619313510417711810610599

7750619411839116178671887954926080112764805877469881153685358393824787426551703883915727427911220922249286993685830086843150370315577718127036926472538606972607881861760424420492047329021112699220625901677681417145703141268742863239315032863083

4902393871077274537354683419396070123002535343337050517253825238544340194393519336022588702958903077437342411957424397943633605450816364239287723924426380849981072259012711245206913921734135516221038437259929631952167927292034202720991976

4902393871077274537354683419396070123002535343337050517253825238544340194393519336022588702958903077437342411957424397943633605450816364239287723924426380849981072259012711245206913921734135516221038437259929631952167927292034202720991976

2306181139963690270928281705173534911443283821443707277424922434265521182178263816941400334227131513214221708692174439215716122182873220314931218208019172663570132965267110095867877037944981941283130894981412661001906960907

2306181139963690270928281705173534911443283821443707277424922434265521182178263816941400334227131513214221708692174439215716122182873220314931218208019172663570132965267110095867877037944981941283130894981412661001906960907

5424457366535895833683857733605835117795895195756444144675604312918496193384784471824268551136049119448435525046635352511832512613121213317814817414150289299240194291204217227200

5424457366535895833683857733605835117795895195756444144675604312918496193384784471824268551136049119448435525046635352511832512613121213317814817414150289299240194291204217227200

2783220640343653332731531902245441741616280725455915272330752918285024572258279719031364514833041810247823111204225756337872088265010652672176212762513227333546231501113267315071008901723889554278151820251050137916071546143913261071

1431092461841711347016120677130131378136142126128103831201026740814490139128567823321126127671468255100135247317591301178739194220139213346109801101134338

1431092461841711347016120677130131378136142126128103831201026740814490139128567823321126127671468255100135247317591301178739194220139213346109801101134338

1878146326622483221420131238156528531021193617013494179219982027188716591578193712309383238220011551627151581315523792222134117416881688116780717501482208642799467646398361359250458837618697512626918001055945920901746

12659891825170314361420880944194371112811172221712161385139313111166105013488136422206152981111341059555107224713639031145465110076253311979901367293680420332624401397336406276131684822457536730600593594486

6134748377807785933586219103106555291277576613634576493528589417296103267134449345625848013285943859622358840527455349271913431425613135921219516818210055291440234264325345327307260

335335557503461480259337566254366345772423444375434333307352301191584457239340336154329724973033591433512301693402904218320714587201164136941278640214232122170214245192178137

335335557503461480259337566254366345772423444375434333307352301191584457239340336154329724973033591433512301693402904218320714587201164136941278640214232122170214245192178137

319227444394319336200295387199284289904274339289283295211283212123643359203277210143241715262223211313542101232582594276017616170145879773986334156287134202195174144154102

319227444394319336200295387199284289904274339289283295211283212123643359203277210143241715262223211313542101232582594276017616170145879773986334156287134202195174144154102

10872125891621901359616265917936798152101118677910558452751441239512238571822196102361337312265107173224959236157373334958111157986372705048

10872125891621901359616265917936798152101118677910558452751441239512238571822196102361337312265107173224959236157373334958111157986372705048

1821118523382581219719291235127326481084170515512628166017781912181314681462168312667542987219111331677141572514373331762124416856481709109769714411455192138696751741269850353547754834716797811836876591032723622804620

97869011481293106799164065414315269258091267910906100898074277585168541315061100586907760403725162889676838349929558360772756965197511252226380250289273276197101488589352327539356318423354

97869011481293106799164065414315269258091267910906100898074277585168541315061100586907760403725162889676838349929558360772756965197511252226380250289273276197101488589352327539356318423354

843495119012881130938595619121755878074213617508729048337266878325813411481109154777065532271217187356884729978053933766969995618945626518631825324620427215066490594335332493367304381266

843495119012881130938595619121755878074213617508729048337266878325813411481109154777065532271217187356884729978053933766969995618945626518631825324620427215066490594335332493367304381266

31152299385035893573349119572341407817352930273141942746301630782711243825752790211212865168346917452427219711312473575269720042897109826521916147524842209307963915088086781154723995786926527271154716421156110817291251105612871047

1926142223732311226321451221141726361101183417552698168719721985173016271619177513308043278217811061546144870315883781679126618506711709117489015961390193941096348844774447262147156434518397610587407061136768642800664

674447813802856755437465981410677638979604710699591539592614460278108575938760852925455612661144266025263947028555847471713338718416027818322217920813277369381286255418260209296228

678594825764802688433510911347628591965585672674613614548581464332132774138451646925355913760743864924360042630453848666213931519316122717322913517712357344366254292410277238302231

5743817357456057023514427443445295267544985906125264744795804061948666783354224501964731154613865411764702783015004305601382611111262391161701571799049263311200159308231195202205

1189877147712781310134673692414426341096976149610591044109398181195610157824821890129163988174942888519710187381047427943742585888819114022954532023141025137431536218288571584416402593483414487383

2462733902753223001802283951602872424102372322762262192602561931285372971462081781102184428918026810526717715524021630861141886796729265625721140164112117154104115121123

94360410871003988104655669610474748097341086822812817755592696759589354135399449367357131866715372955877932267656543064860383216840423216431417928225030012567431420304285439379299366260

56124083197711125612277681137594222112123724798676871467667297840726160874576490757596291270418015673151258608441723345164101097834154126854998153153641302749117240113801852400720571773221017121884156832161168636424556044491486832042382230331182149

565492831694674590365425762318599502852528530522575453461505463293897618310445418206474944464105652085243472534964166261132881451292041361851331849150291316207234312215174257195

565492831694674590365425762318599502852528530522575453461505463293897618310445418206474944464105652085243472534964166261132881451292041361851331849150291316207234312215174257195

45993318183411014411164581031513537993031956954677713171579268546331517837894096488354772237164765691459078463693196943868358969346911727462414410300425874103651010324163335521792154518401451155713372906997539370350654107446026712028198726921807

424730981747895411056654052875317293352901651563581254454206418594248433494376745685078203715753510842407391339718394108784859632661110843881380727922380376461859736153432801653144216771335143112242675907487340347603867422024591785180624161591

35222086360359840527636559529443941962737243638933529532931539920072358335045529613027851373203619236603212207339325588992721391031631161261132319052300305240240212243181276216

4482735994184394112432605202114334086534094564083343343503713511746424222253173061593048136827539316638129018731231443010616712099166125142981268047251223177174221139142169147

3522084773283183121992073961763513145003383713352642772703112701454483471782612311272426026822131112229124315924425231387105968114080113791086737189173138114172106109131117

9665122901219944531243582941537185737057806081291947547567532622110054824490472868621171962241826452919181310625039604933333830

26216439630637627217220335512221621844220725028118619322818817792531277151190162104179462942122429729416111717418832250154835299925345594623137162931071441148410272

26216439630637627217220335512221621844220725028118619322818817792531277151190162104179462942122429729416111717418832250154835299925345594623137162931071441148410272

26216439630637627217220335512221621844220725028118619322818817792531277151190162104179462942122429729416111717418832250154835299925345594623137162931071441148410272

44901351167169463046421947714127978359545657825429527034013354879390373911140574411283138440681422255013128541651494892423066454233859920865401129288139337332394480321261376192981556958734673428171051202043217811369164472395212870165511520816239102486148458932235640220230953403130336359101897218679

1784156127582208195925261359153324091030197315732469168417521732183814711665171916879042663193010311764174570115803461537158519048011560122310361554142524376241091454544969633636540611375185143599589877210861032890735696

686671107389673494552659298540678260799965566365266956167066363835796773737870864727862814859060668928459245542257855392821840916120342425026223025114867573400366306409419319275257

196219338241202283115196220114189151268173198169198155220176185922742121102052248918836154227194821991261091231602886511250391366867525739231819611077134128905963

49045273565553266241139676529259345673148246548347140645048745326569352526850342318944011243637949520239332931345539364015329711116428818219517819410944392304256229275291229216194

11693208140115159638415678153119164107108131989394129129642191306311111147114251121061395411884661019218762113275057324250352061297078605759685058

11693208140115159638415678153119164107108131989394129129642191306311111147114251121061395411884661019218762113275057324250352061297078605759685058

982797147711721110142277085712685461038847130692298194910718179019279204831477106359094598737683817383587310764638506845488757801322344569266291488351332260325207112733525454406620554503410381

227134338276232264134168308131212208269202224258226193191208168116360246159211199741853317727524810618712113322218626671120626394737260852820147103110901251221439486

294249461343333367246270337165256245390325307267413266284325241152374288183270383131215612512362961422421911482482123589414769941408794721047538182156132125183164139126137

143105176164204170169161178661521221661261551441211071271071597121512959116130589820951121294513616672961091773573343151394535423314967849448881594353

3183095023893416212212584451844182724812692952803112512992873521445284001893482751133405931225040317028520619530927352114422910110320315212193947140308188163147224187162147105

70584099810477661144512591918430764597895636759739759579690709645378113080046472969734165316255057470730861144650962358694627746619525934721825021725817063613373380364438358314327285

58071970175354881939143168031757344064048451752655644952652344329075055532654850927549212239243147320944332639647443464318534013417425816418815417612554441239259241339259237230223

58071970175354881939143168031757344064048451752655644952652344329075055532654850927549212239243147320944332639647443464318534013417425816418815417612554441239259241339259237230223

1251212972942183251211602381131911572551522422132031301641862028838024513818118866161401581432349916812011314915230392126618589546263824591721341211239999779762

1211152682752033071181522281081801482321452342011941231581801808333423112615218061151381481292208915411610914414126385119577682525857794071481231011039091739057

4629191518381051192378129766225461412298510210141410144451140774972463522411202098475

630646501026169304809979635904679694634306781495864004771514854606193417450534965641728097361640629185568479650385045165339204816503725784313323336764666463777202109381412961991451512951801185919491227581593827523292270232792425181324542306

630646501026169304809979635904679694634306781495864004771514854606193417450534965641728097361640629185568479650385045165339204816503725784313323336764666463777202109381412961991451512951801185919491227581593827523292270232792425181324542306

35530652244622552816318738717837235234827032733538121827433331215845639619226623220429410725524325815422418520730228041710621979108227671081141126829318180187117181152122127104

14909942560149312002491781101815666591533114713781008109812121195916117910271618615197713835961322113910471175352937119112276079967018241030105916935029302924729783434333914272711461401608849785680589431576512

3722706964953227222422644162204943003912893363213682403023104932135233862093833533712951172702283542093042052173022666851562679414535076981101378025481181263206234160115163131

222218563645237915883148137016382461131423501795234517981869204126211528181718052090972229122839431778160217861766529129116281642773147111761348171217412511672122944869814214576426906674132011864101510008121202831634858794

40018495452637689925037952823260034648727430538141034835436460424463447922754538161040215629032541329037220727630232180422835691178516921241431749847624219305301228153114183129

791574984800579105442969786243977753275056170765074454255358764629066580837467362748758318044260762427757541646062458884323942515622249212322121723215466615273331215385277214280323

67646690079151995435549672638865548670157150652047438257453965431781567137760146253353021243559451926837134334439438276720638813616853113717519420014367635276357266369263183267313

207815303209254320873161129114742603111824391550265117521801199918981454182616882040907321622791053221016567741688375164915022025942187011211079170315902923755140452662510135996395766204362122002101511059291186900766826727

458403713822531720290367672279520369633439452535471365441460398258683556265496401199460943943444622185022682734054086171742921221512281391431411398641429222241191276192192200196

135112200180192211951171728512511920410313614612112412211011351146156731329161123261219112370148678710010920561953338623840333832121548068627140603751

32329151364233950919525050019439525042933631638935024131935028520753740019236431013833768273253339148354201186305299412113197891131661011031081015429275142173129205152132163145

369351761448395761292309495204560298590407338395364271393339514170783452224532375148355613253074691973672272113663087752003601241402571221461521149936557219263246242218158181163

14186272185150276111121162812091292061051441501098812913317760249149781911285510925121114165761528679114107295911384346835553534842151839280678968545656

22826548926324548518118833312335116938430219424525518326420633711053430314634124793246362041933041212151411322522014801092228194174679399665721374127183179153150104125107

4512777045254636512862665672575553675383693614404333364443414291917085212414623941393339136135844320435923726231635455013627311513421013712411814310248347224217180258195169168136

4512777045254636512862665672575553675383693614404333364443414291917085212414623941393339136135844320435923726231635455013627311513421013712411814310248347224217180258195169168136

80049910317486981029423532869378804516890537650629630482548548699288104275032372048628854012956949365132364238933361652098124547916520031820122616522414987669350384312410295247277232

66244778462756576235243773633463243170846356053256543046647453825381363326256640224644711645139951725551533927352743066417737013415326015319114217311769464284278225349235208231211

6127129577011641537522855197364651383345408019985936904919587545170355527426043151326519243426101326168893145473128192013

772511864631513042582287348538444627193734811613158256435233566443643372231829471663644122324222510251610116356140303220268

306262407435296406218293409158325305418313270334402241302367333127422374149321298129315542492512401272501681432892463781322458994141841071211336231266139151157161137121153133

306262407435296406218293409158325305418313270334402241302367333127422374149321298129315542492512401272501681432892463781322458994141841071211336231266139151157161137121153133

23022429832420629715619832012923323032124421226033018424928326095296289101244237952444817819517790170134117222191286981886374110578598984721211100110105124112101125117

763810911190109629589299275976958747257538473321268548776134716715663378034266755923457262031272223351510553941523725202816

384350743569470565255359618247463374695437441421414350430397401203798555223472376171380744533724062294242782153123486781492891301413112101501241437868304273226232257328272209146

384350743569470565255359618247463374695437441421414350430397401203798555223472376171380744533724062294242782153123486781492891301413112101501241437868304273226232257328272209146

321315584513409467221307534216408307565376384367374303357355320176651483196399313145328643763153361863642451882842935591192421121162471641271091157041242232182188229261222180119

633515956619834528431556713061575440477342812714772277363265210775770436033272855119304718256446231528827624144442867502927

8746281613112810931513590800129950811397481335858864897768663761800995394167310545081005763313833157834720994444816552415746752163539173224427055234928423729820981940523538476569542419396305

8746281613112810931513590800129950811397481335858864897768663761800995394167310545081005763313833157834720994444816552415746752163539173224427055234928423729820981940523538476569542419396305

5183681033699660995359474739299695459809517518492409382454473644255106362428860745919051194505469598279506329250442481106626749214816836924116614417413146611296357311349321273229200

118982101681612037813716987163111181117130152143113116112104462161569215010039114281281081914410377671231161944994294576484639422891069055647189675530

2324381835231417421320224414253613142013275442618372363092622131933961816281413758116512622616131714126142

215138332243237292139172349109261156301210191217203154171202220883502481102111817817826175121192102174137921631393476113360529949664970442419712111384135120739873

240226391324313327139219375138295213328222216219245202236276199143319259148243200101215451961892059024416911919315236766164656912586799580372420314710211913912513589120

240226391324313327139219375138295213328222216219245202236276199143319259148243200101215451961892059024416911919315236766164656912586799580372420314710211913912513589120

240226391324313327139219375138295213328222216219245202236276199143319259148243200101215451961892059024416911919315236766164656912586799580372420314710211913912513589120

120596144322102129561388131079160462232629532127104712781362120185015361161341267234102110787260617685161498239106988918961242165184696211391334577114312164399935875346563623708454164373773019331482886711486673571

120596144322102129561388131079160462232629532127104712781362120185015361161341267234102110787260617685161498239106988918961242165184696211391334577114312164399935875346563623708454164373773019331482886711486673571

36233715107353752425232305480207117428068028540642635025950332712211611223628224922600161451653122756634775762482933194132237539797118373288102185198226136481422244670524271223130232178

8436242922136792037135817741124415208867314477628729368515911033834219151121871482563168411683551047174757614123376510755986698209213534892136728156258724437842548231811623154861263958615488356441393

50764011749865725101694432784277678828775578424063664511490950275172403342764810453524766757555729364900452220214197106946214300476920324411314930174470400265141754296011841580243816111789159116961082590383325012576260028122563235220922170

36362849529647553614489423343174481119783952290744413261338534843749287230353377315117934778400219863388324514062929734335730353319141930182225212231512876446412092063827111717611194131711581185779433271917461771138319731882176414261533

561571702636448773354482586237424335529492433461581430455426435264528454213333508235411933533984091602853983124163825941442167513422413018515014310043359177192150309251223232213

2292012963322332341191883009623615124815316520719215415519417487242240120225172841674214518518790187112114156172235971344655103576761613928181101108831181088468102

2846207742983787293338871861250439251645329224213664261627872816297622882425275725421442400833081653283025651087235159928592452272311692546171516962579232236359681713706928143410071065947981640362217914681471115015461523145711261218

144011622202181714872050944110319778991626133319251250152415431423116112411433138468319791555950151212776151268335126412651450613139392489513191126205054589735746367741747243351130315711147558051217839681588666637

597698435256343667335449615347463271405468198236285041293812425649233738285441933252122228262113131124627423682713143017

4914057396045167473813907543455554457024655064965053824285004792317005972874594322044081154004625332214173383224483816001652951361332501361641371778564329276263351306235177210209

101571419478132586011145956710579106819256717099391088864107724398168185115377650375455973058223849282724361711795358674743294244

789624122410768411115471617104547692277210576538659207946527028097383941089834571896732339724192741662753332863498508763649126031849218727035022726025928519080660399442431459390368384367

749968811309186797173144184587637187883768958557099436644959986596623650526653614893233222113767811352779806748294959701351552450557457368666564722415362576044136943325579617112760352319212302215625401630741855934104834380642163323279730892966

1661814171981735957815620567312108244142134159132841711423167039125081270153591393314511321014819899111133150539122204431001014961707549153248919615210374607877

1661814171981735957815620567312108244142134159132841711423167039125081270153591393314511321014819899111133150539122204431001014961707549153248919615210374607877

78843051177431642678034184551154055855036905421831210113371671064075117660121951154436659142110113734626321372438282323159136905636343433

78843051177431642678034184551154055855036905421831210113371671064075117660121951154436659142110113734626321372438282323159136905636343433

33292691518037193129562919392887378117123864259338992671261628492720219928482600371514004986332415313220285512422571561250321183214145428702024181127542570523012902291791116014648079729041074664282326814991912154118211370110613331234

33292691518037193129562919392887378117123864259338992671261628492720219928482600371514004986332415313220285512422571561250321183214145428702024181127542570523012902291791116014648079729041074664282326814991912154118211370110613331234

56236912186865181329369364705291878390835520489512502388535529888268100863828773049423347090331380602357538313313468448129333353115623131015420917021914264813275476344322271177246227

56236912186865181329369364705291878390835520489512502388535529888268100863828773049423347090331380602357538313313468448129333353115623131015420917021914264813275476344322271177246227

303227584339297533179269391135455263408323242251257235291303381162459298172357294162257662492822901822871832472472566281402507213216479107108976948383110214138173141149144114

303227584339297533179269391135455263408323242251257235291303381162459298172357294162257662492822901822871832472472566281402507213216479107108976948383110214138173141149144114

319256748426328942190282486158603253613326319352288264357294747174737465201475410145390792832464002163652492173073228842414191141842441301521351558337650231301266229221122156182

319256748426328942190282486158603253613326319352288264357294747174737465201475410145390792832464002163652492173073228842414191141842441301521351558337650231301266229221122156182

2783524603592615281492012931223201723011822221862041701971873479541026911528424110217052208169290112219167110279211571132216639611963887783643331413418116015812698122116

2443202543082032591271652491091931432321531961421721481561671978126918993171184831284216114121157154149762581743486111451478542695550492816089104781251028898107

3432206515826922364413127296929264432224120150141418022113571942104728795565183421372237110212493421192233155154457782332410249

83989718219598131932555698101544012876951147709613769769529808686117435712449344179497982987091686505148934688795664406707581853436793171311399257262266321214821229428652504486395334351336

83989718219598131932555698101544012876951147709613769769529808686117435712449344179497982987091686505148934688795664406707581853436793171311399257262266321214821229428652504486395334351336

1625182423581876158026141086144718328091682118018741536130814331314114713561353153766519311520686152813976681189291107911731437654118510778681334123822755309552674846593614144024783171571347585766611868689717625647

4763509406836339083774027032636074297034704435264823575034445962497976652766194042264109144938654327046631429537843992320339910217028014614315217510450524258298229246228224207163

2141604922632176371331892871163571473021951662441481461651723538349028811033425683167551741702091582541201181742225581322395795100625769724822389123197163105977411969

9351314926930730106957685684243071860486987169966368464468873758833364456730057573735961214545661768522646564345578257779419531710821927915321418123116585434204271219517364419299415

116397694155762225310204185537222888415599753112262135321280110821965395271042476801110013503124171317415538125525720105969129502612363247111356683991235856348840971438008458557890168556116806534074787593836035757508948903014267710860595420100594415263148212300649965675

116397694155762225310204185537222888415599753112262135321280110821965395271042476801110013503124171317415538125525720105969129502612363247111356683991235856348840971438008458557890168556116806534074787593836035757508948903014267710860595420100594415263148212300649965675

3261263181171262424180224510306230819278415258226237164489754376946424324195209183167616611109022548917320652334014044163440157166961401649621320714583992581431500111108510332014128197

326203663493388766214316427202500285469274337316331232305251519144592432178430293168299682822613892003572202142863437651793138913912087111941318834515187292217209153133161160

5264346558424698013243606883295154995664704934264563504365014753476865892754703692014701121832369496185373327712101732662418530113914122811416813318110764379248410185424305459200239

5303788056474187522953885692405774096724404904294533964184714382435955392464804852154089337334143019943527224250538560517632211016221810615813716011440422238289205276190198201181

41218275589253256928603552314548513645326320139683157401243735574234533023293310937561321281919112197643232302726191716

12612230416311838087881687217810627212211814510185123152171764361756517411849124261291001596912178113791102656711140476337506254313119391106918257466641

48732379869251591330639668031266439964951355251351937843542854526479161528557552919242512476735644632049234639360641786319838312917723013417815316612055493263335235304222247208209

494263641767360780342348601289561351612404457398526363387454467204639532247494462184418644673454252134262492194023386201873691151612051041721721759148527210290231246217176185201

67946310171082618107747457387133680560291960964557569451157471761729592077334869061430061513471456161722552644842459253088325747215326328314619324224515364587292369295345276246307296

489273803582448874232323481247588313500288359348335301399392475180650488253491398171359753672834492243652382593633457822063501081781989914312912610883549221290232270186155196173

2802124714222714971912393751803812343243052532762822252543022871744283451782972511152456251423125513422319527035022145010418281981166511089946526289130210136200152160138146

4523795346673456052223664702194133124443223533363713083403634102055144431913403601553568268228338317933122628851229554013024994125170841601321517238400164268187247168189188149

7656067641011634110846259093641872957583260065760878550466975363143481575036063358831760810022055666522705454348931377529781762442177297311142321223397148232508473610247493474543280307

2630154523118513204434031644180644042485190854881968306615761689197713023240453230907620303624461133166114141407426379085497204137971590169640492661030756132535852004182511601542194814172540215215991034133519411474120961747864592971614113081884

14585387188194431921532407428514825514714817511893162120290643742289825316675141302121302121231771001061251514571242045377876136575852512871071591379169606656

26176614126682424124873311182118321134381062782719715416938302411161516941728614143776636415342617244108

34019967446932974620632439515844628438423832134329624427932244113653241618341629614629353287232334174280181201295266634155264931241447589881575136351166257163188127129127133

421782153815135726771621521081542411611545343123625221232117673321

719428874819500852343418763291623478718478460475615414508583567340706578278505487203479110152837456725244232270783443475819438114818325513418718321513068516272448296406336352218248

12898254181142313911041698324113217912912413313810610713620173257171761931554611226134961216912879145138137328591114762784852406738292385912611710075676950

231232121459920159291231151418101318103010382272316832320101813121051617581114420847486428111714951176

2522995136694149572479495339386148373740682212644265738224654432462631302729338323331814181312161415555274325261882525

104180416271755113916666828541352602131799313831067100610511199883102210851135584140013165831077991445101420615837691035468812616853122578716634337322383905202794313673572421191113534772516634547547460470

8780206141833906110196521877115076741078675106691892823413949202124469414948411473105655671583188510535605334294030341622171119835062325831

95059911741488805147466078712385588859261051760851803771656720102187475510901084426837729373884221469363883338667666018282185628113836364626135047430036935033121418885151910224118867991070371448

401931127402516139408644241031945024219354403271578734973686396235732851390933605558182769174749228547503940171434177413202258644052097384824992190349033127981232635131062167619741065133812321534935454538221202941250024011882145518471597

398930907339512839098577239931664991217253973255573934813662393535402833386333395516181668664730227447093915170433937363172257343752078381224852170347232867908231134911055166819651059133012251523931451534921012910248523801866144618371584

23961860436131252403510414471947307713393248202534852111233224452167180924092046325911344067290414792870243710162118455192115692635125322781528134820621899476313332079657101312366737587489275532683189127517271509147211529621158976

51342419250284277680331625312134547256415440179171929146122723570154044106679734847278308941402943351227333616723232106723735122819

611537103176056612483674327042917384628504864915585203805365207942851029670286647542229466104477439621286498374275527522110732345413823427314420917222015667688255391324334235184253231

9316591706115189019415587111130509124971512838507858607976038647331284380157810654631070864424739162734521101347293954946381178717305618182313784212303362723402061091240539686580537444288398358

30226333316711283321431648162427331846214211511911412510245301330193614201826731522789687114333193115211691013

30226333316711283321431648162427331846214211511911412510245301330193614201826731522789687114333193115211691013

1094927139012911074126164591912716119598621325841915976751733922937958479136410985249377903957672038298541008434849633559856821120133463223128951133035829033721696829552509407551494348479396

75656775877765473040152775036555853278350953157241942753251953825784167329553044922844712050644857425949838931149649065515940015017127715720615918613044461314285221363296223274241

75656775877765473040152775036555853278350953157241942753251953825784167329553044922844712050644857425949838931149649065515940015017127715720615918613044461314285221363296223274241

57509410974100798911438784191501027381595684764710454616764455516639666527138897265934137162579424041402221675850393554265443

57509410974100798911438784191501027381595684764710454616764455516639666527138897265934137162579424041402221675850393554265443

28131053840534643116530340720832328945128228233125124733433434417541937116834027712226567260310368123280206159288266453134195659315513111290111643130118017414715314499151112

254297437355311357150271358180279264377250250293238226300306273162340333144280250117240622172713171082371941402632353481041535677132118100759153272311371461261341178413096

2713101503574153249284425743232381321342871137938246027525543395115431219253110530429162313121520114704328211927152116

169214831925180416141745106912821927768143812481846119814211323125410511322138512118262205139079313421171676119134811381147139261712761062877131411421951415843376427720520498458442323181992872635605787695736607586

169214831925180416141745106912821927768143812481846119814211323125410511322138512118262205139079313421171676119134811381147139261712761062877131411421951415843376427720520498458442323181992872635605787695736607586

277288375404299345189257421153270251344215258276247182299240218170499274200266253139224672492472741452151861502842104781112378884159133121119966949211330151130124143145135147

116598211741088107111316848501179457879782117877092285276065878187077850712928514577807014207392206877468523577967065807977241083244457205260446304306267241196110597412363354522428485362344

25021337631224426919617532715828921532421324119524721124227521514941426513629621711722861202154266115265170147233208390601498383115837172105582218413012112114112410611095

12531719112910122751429012534787175671566914583830613814134719421984812626936673017647792677553432110697100982444710045669742208134226511525439201973566281271410947696484061302210939298370592446414645651011035092781681325131681882660626745968464764382295643463605

68943368674874166453650092053252862486156361065462347955653649976274559461158247724845846664390656124658647333552865855914936715619925026321117424910884363372249337374224187268245

68943368674874166453650092053252862486156361065462347955653649976274559461158247724845846664390656124658647333552865855914936715619925026321117424910884363372249337374224187268245

3392583393743903142462545033043043394472973353423912642652802664573913013552962741302472753764952891283032391672903592867419091116149141971021516544179199131165191124112158134

3501753473743513502902464172282242854142662753122322152912562333053542932562862031182111912674112721182832341682382992737517765831011221147298434018417311817218310075110111

187141272213229260125103287147177166263196178248191146176162168179244181156192118811501072042421949815812479192156228511375966946760806851181041209212412983539665

187141272213229260125103287147177166263196178248191146176162168179244181156192118811501072042421949815812479192156228511375966946760806851181041209212412983539665

1541252121581701951047422112614015020815914718015811414013714116718814513815110164122931572181527112110861145123161391014250765145735741157294718910569427558

331660555965212966213716553731683332362527125636184117172814472442273716184733671236171618161571110332262135241411217

1165566171195211314133201161072106964144621390476011302412347866290601172485526676691572287088333809708932323680927161023891752622078144073805389806284119701035065507686122081015227836555223138814221978032382527649623541579835955706404922359734075271639823295

6071307353434353518137354189278348251020926238521523530254138408831183040259829403159283733194336417870260228072008254019376882431990445326775152691735133873685635051229307996721311713757210849994030783983453419263960486923511894119516631198

124921761452021861059720611113818421714111815914011811716412915819911012411713954138781291581236714285669513413329772655815644255632299256618810456435364

4302834503774214371912784322462823464673533973583592963442622953943963503202962761352762433384893371723392781792733143727318384130145155122821668436258142120182227169100169160

4303135424867214073112206465553145426973685385164303213033412621334404350136029630218325988666313633603585723542833421052432812249719313941491835578355263154260400179223125117131

508723854175334538372705358221883541929718897449385421633085305521892305183421732473264872195255416066189320901636186718169769429980363320804099620029853163535625681046259576017531348694782780932515848633921157435194199184114469271324843

55233514653569188060781830014159956536634939444970315596487375865386358442654259389049916444592557686626322818584944269155376028448635816776341030213781532965841546343412551739249221972141152224561550593379136222425431635932163150622932087

46922811540259377088690125543631851430774218372459814769416267484599302435793526331941755622501850655906265715154277223247595104382832216004293125673095461257531355298811081492221119221898132621341384505327732792070388431361841124219701767

253173367364345257124158343191253242308258199256274174198233170259256362260260161911981512372741901122531481272052462126612347929595705811050331441331121781581177010288

57853076661762766032337070839546848374256951258251338648850040155756654544346041025246930854165046824851933132748147161912532310015518618017313821211655370210243254299205194221232

61307443795720227232395481414950485252293916703442436725421146354126422316322363842911161113610213342219382918152610

61307443795720227232395481414950485252293916703442436725421146354126422316322363842911161113610213342219382918152610

170511387620939205941783419602124511518323925975515869139212135216324170351661416578128571366115708128937675231601941588091623614114654712996328615796135901434762001350010332804214043144061852940148749397539691199976645071452449093180164296718613666859009447117691005067216174

1377124316421800156116351073141920257791434115217521622153713201454109111131335997607207617147281502136554611043041433123511584651085847739122813531451315656308303140791746536442121811173666151745179213461105551521

10779801207139812821279866113515746171152895140213131200100911218378771065835495162113125721206105343587524611391020902356811657607995107411082435182512431130723386290349185815755304063446281074891426382

2392713262862433141752123681412532243693172952412901761902182121043992771173042349520658273207228961791549724325227160120656128220787607968201511491207916028323010583

16620312517112132121414372720141616136113652015292051222912138177109342891453342812674171251210311991

21416624524932119514916526914033116026824620117619920017924817210323127412819218980201612142161806315011112117318518244109444011873101697336121199163561141101087786

21019117428418629316125832612318216425824924618726314423618818110326127410523218611019144215208176581661421652062141783910951632221347357612722969576591132231698986

287274306419353342294349391136245238323366317261271208192313189123438348152290317113188592882652298321417217723027429453975755319182806786401713613180951642802389886

11172136129154118761301886512795147108121130829367927059227119551591073277221201127648857137134115155386929211559933314310966526243671471274840

300263435402279356207284451162282257350309337311333254236270162112455402156296312111229582942152561092741901322332793437213857602771947974723330161131111107164272214125139

300263435402279356207284451162282257350309337311333254236270162112455402156296312111229582942152561092741901322332793437213857602771947974723330161131111107164272214125139

22321680342328922381258615821823314912342395196030632172228423502207174720202274170412143109253412572188189478917684292066167820329711898142710831889183725505621212564585131985576665067545820113211179956817121913081198908830

8816161186988963104165667112564839417531183848869934869675794862638475133498550078670728864016787264783434071755344467579293722346224920454437929924425818376459480367326458561552365316

44326636625749671063248387262494338263333381363592687502741674582834342721337166203412119280101612763420223016103672511

59431447273132535586488611810660817966715410547431007440644622576136595639524122605675164533183534271023187425141325126332222

2331972872472462711742213441222772102952433052612441892562292051673213001292051806619758231174233781881491531702162704812066591748685747845151291291007912415913995100

21911625829021522214312527410418914529717015922524715214817711892374201114153158731154214713123070178142103184162201631146637715563545749209911658701098710610353

165131211176184190137108225841901222251761531741621362002051369722919411114415956140251621311585514110585136145166378343389073635151401795888063871141346274

16197220167183169100952219315112018813712215211210110311394632471578013311444903012294129641248960921421593966294182515139372411607666527172735856

1351106422371904141815459261152189375114541207188013241415141613381072122614121066739177515497571402118750111282621194103111986311181874639121410451613339750315381775476467406417275125862699589491761747646543514

239136309286228259138211295117209183294238202221165145196217184122284284882201446314737204156193892161389422116825350118496013066646763471913113479641341231056886

57643286284462173438245277231855154783859266269953353353558248528381469233558551521650312250546153227648637129051147568418233014620233221819316417811439408296285226324311272218214

536496106677456955240648982631669447774849455149664039449561339733467757333459752822247810348541447326647936525548240267610730212011931319221017517611467323269225201303313269257214

3463245440974036369639902517302649832022307728454378328634183480329225032747302427311439492641771826356427561327248869135272670301413492901208514252806308141348571902791788278217709568851044654353206918811436128319842716211213551291

64148965668058960935945076532153050873451552456352437345850941323070261528749741120936313959942449620448533321347239357313128310511432216013816617111744304254199183297264275254188

173130183200183206941272581011591531981511531651371101281571265820216565127135409637144127165531429852143109187368440468451424260339908367608687827764

109599712475865260135458191157687911798598676593717896477690276527147687938833737896474205221153836202529147454631443834544526

22416124620725323114619725511818819226619818819416715717219713392208255102188114871415122212917785179131871611462294989302912551586555452011490685212296999154

13513913014978866766117571027211398104871224772799543114997310672556124861007528816737797483265814247522183427258553533275147404144

274619133293328330163192211925164123166624262278353027232832285427292098221524612197117440983484149829722280109120785342873218624581092232217101168228226253394672154466963823981571801697853520298164015901157104216402405181210681079

237210215309233256240170338181216168277203227281337192182187198123293299107226206941445020019420095187156105184222246661144640155104734473381911014378701351551117875

182539303218720451020433313201916121319209411712301661922721172138118162323621831914437411318131810211784

1267227830615013810814520271115114164143168142128113104134974521732086135128351082914911211049111845610312216235732429969142513518209469595383102885345

24204146465416265318203049282441302026422218674318343292633517299381512223139425431510597104272711121533191312

2819412535341835352021273721402225162226271144371534229235282739919211225243810229111718676521712101516293058

35113322332810244717343639313025131919202484543163117161752726191919716232325121863322011274326111217181513159

643810352871071155812742698098617767614979535318109893672502346107167772474503046617721441925775416102368464760423174452019

413080455462243769395935775242345337393649338210822604320371058444219614828463312314301717301814914175392926203438223220

1014155646348445110226546377765254484961494437736724575820571573524839593853405697122881042172421363355111227253738453218

132126173120150173981432437914011517412611912913610310711712865177190821591044812022145109125659910658132132175318243291529444284635119177546288124925550

15132215262113382915101835132128212112221432818101918141241781981575341625353212497377176111413669

35820545839339944931029756722933231644234839239136429628633728915057743922239630316631872373268330154313223149280354471952439596286203122951235649226182189146247316267119143

71488990897852751214670679083719068694868611816695507450316616746058354953296364761348171645262018241312495129294645442730

20411212816121826131018301819172061991643410816124820121581314971325672114114453121313411161242

56941263070060560838556379533645443871657758154060443245549643821182161530267252024839110861747248120740530521846465572812328513612468738616315518310654331274189181309695457230220

6135585446563744803443454962475242554451442910673385142214110524345254536145165521325171647371491074392025173244422013

9185991011001286879132396861921129788886792957544981086497684677228970783171573787671121735253380482431521711515136444872464244

13417243131111530814101920132721101513165272582089113178134229861823510541413452531091268101384

2274617423126273919233628242539171419152174234173416617221242972416815302912266822216474314174141221171213

123831121171171168010715973981071411141161301148988978953174137571119144872110980121299677421231061232348332575633931291512567338367577834848

131891681921161329111519372119111238118154138136961011271035217114958122875410233152961195511610162122128150296235261017432313124205267594385111804064

542566648874466885326048796345803945405341359573247456164998144512257302536416818331314574117161781484641294247322420

503170365363254374367637814040415029544246251215824533020359644757225523927577015321214292215131896333119212832152319

362280389440393471283318532211301255465377412379298259290353282171490437198395303132267743742853361363362231753302844378922886892941829794967442198194147127216277216154190

76521487391189396095351215911448626339327454121351267841956527471855606053944244526316754751736623917222017111253780584747253324

76521487391189396095351215911448626339327454121351267841956527471855606053944244526316754751736623917222017111253780584747253324

124216001277136611581310110311171624621100789213471313114410601416950885111682658512961112534879100446897219690593394036586086772211389501183244465229282643410326314312202147546448386319765678651464477

124216001277136611581310110311171624621100789213471313114410601416950885111682658512961112534879100446897219690593394036586086772211389501183244465229282643410326314312202147546448386319765678651464477

133170150170131161951071736511210417718114212018810298125111571891667010310257113271418214142101103791281111673854404010162334338201854755349100103715351

6978887608076088056246529533455665137228046936178156185026514873546836493285465592725239449951150820147447243466957264212125612517539222920417618312078301248225181403389374266268

151126116117132113937717164871071671151081291268211897715421014050105131468317121106103541077571969115126643420686428282623197663464878831025156

26141625127228723129128132714724216828121320119428714816724315712021415786125212932535814423418868178217138245176223599130478255616765393211562624118410310494102

11910119811915027313910721167104109161146164121102101939994522011747117716532106341271011396311889556898118315428261047938333024186170537074112804336

11910119811915027313910721167104109161146164121102101939994522011747117716532106341271011396311889556898118315428261047938333024186170537074112804336

436794485551514170355351876564434647373743318872325344113815455371235148283741451720149342922161389324021263236272216

7634104719522288661413251587481100785654566251211131023912412121681982486840674127315773143414177050161717169293032444276532120

7722608593049465792488515362701910900451271086140959070177720739474305776608171345829336510272874639787213630630205936146869316375626526845934445935656173642983141781398817881800533234202262209022011464712439138612948265040775241452330332714

4643067157375585803605427473004494416795245525045254124004953872287056292935475091944058650245445122540428730243843658413824814511841727215217116410551259260210161281400343225179

10772222111102151761011785892941569912010298887791816017216462110113428120114838252906759107841342961232091592435322716697257446996793526

17815025632524624915222431914020218928224124723523719117321317598304251110253236731773322317820579166117120184176242531148050171120746363401289110867211417915410780

179842373012101801322172501021551582411841851671901331501911317022921412118416079147331651931649414810312314717620856734248155935473693823101786745981251108373

65955273060362563242361487235060551490563160755158843449060842229888966326161552629647411756652651321147135632849554563714727514811448429917816718511354326255234196363518441241207

65955273060362563242361487235060551490563160755158843449060842229888966326161552629647411756652651321147135632849554563714727514811448429917816718511354326255234196363518441241207

1611732722571871931401662849818415122416415415316111714316312382297199871461876413445172142161661558498122169193381004150149786342473418979570721041151417160

701051278892766277133481006810876696378416583624212372376410438721880727129623826526986144415268044272319238514725304251843332

916814516995117788915150848311688859083767880614017412750828326622792709037934672701001072456262469343619281110464845426264573828

271648293435191119420392850776830241842184198935111057513195117553413221314151089512131292191049610711021222214207125105

271648293435191119420392850776830241842184198935111057513195117553413221314151089512131292191049610711021222214207125105

54436364963459563842643488235053044669844551056555037643651341523574364128247736417943014146440945919247833525948144565810627210912928118217314516010655321350254188326287272208176

2931563393763113552371933941782972273572252763022761992152552051334423491572582148822192260207230116205180126258236403661355271153104988380572619618816010517216313412692

25120731025828428318924148817223321934122023426327417722125821010230129212521915091209492042022297627315513322320925540137575812878756280492912516294831541241388284

1451431772251701861201571887113411317511014616415280971139030215150811359952102281281031375492895896113148326334348268303639206928549637183656639

1101181312011321378713114961949213384121134112667888702116612167997234782010087110336964427582102235230276749252533143756238505563474732

35254624384933263910402142262530401419252094929143627182482816272123251621314691147151951166317231113162018197

27817229233230524116420933818521916927817020321422218217419630688377251117206161951575818541116592181115921651592946113661669644595975452618911879921201466780106

27817229233230524116420933818521916927817020321422218217419630688377251117206161951575818541116592181115921651592946113661669644595975452618911879921201466780106

2101176826582626215626101332195929831206196417552589201022752023214016261734198817039962644240011692127183184417033811937173217807221647128710371745174424205231163508560157310386566146534391851187104687175799515301283877761

322296426387309438253290459194327316383355343285324267266320203170462300181316324163292702652842631022712042523102493758517572652231399992886131145153140100169224209117116

692713897862807857410683103638366454183265977563771254360459455032983377542566957625257811867961356823855142930757962589415837517019162037219421321413162415340240262335538462313240

1362316181971321614723121616231227221863323831199102201291120119121333610671818432538151568191353

982695121312559321193588889136554388981712529101041971937755768992849463117211835051020825386758175905753855337728584418786761102524655523327066047632728231322081562488438359444694537413368

9258991069010374841028070749974100114144496960832814411950918734651668708534775951589693284827275233322436228575038303955622934

96766812041177108111616648361391600903853122790596110578707417238307014001419117054183777035072519990679280133977556439580284198321946423020963943028320628016491556531329362546592592386317

2222362448261641451840194327382631253132259613113352792544327261831151623353562477432465114231322141523151610

36523338435033036023528845717928530041630932433527423026024921210940935615024421610827261237262258108258208131262238263511466457159106107659450281741788910820716017012995

356241439457427442237299523216321299397311318403308260221302300143536472181330281132269613602992301172981861503103483918715087902541989072995735222162110136171233264122121

224172345346276333176208366187257235371258281293257226211247164139413311197228246101159732662042879618815598207220294751447255183102806476572413717810810415317614311991

220224255286183206140251336134228171261258231205294155188194169852592201172422117618457215236182521431349118920721640945560251145605865532513911185811322622028787

120149150142991158414217277140104165162123117198741301059451168146661411394210132134141111338080621191281232156333215588393637301480644854721721295156

1007510514484915610916457886796961088896815889753491745110172348325819571196354297079931938222896572122282311594737276090733631

2156170023042559203023691574174027851198185314992504172320131928190416351654201614729142635238810191776157385716033511739151515827181566119589116251662208646511604454511141760599586523378200121598974565611251101992782777

4511261530361124181226263010221823121424261647311526175214231322102412621164571261410583541171691516157136

66257278475761269841754690735255045876558661461455050447461448125484476831160649821851610150741249219852038428654054862414732513114039326616417016112161317319223210356342354250233

5414556187395337024274448113275323926514435695035173975065823972467726033074944102344451034624384302224023242444434856001403361111172772121831471449761362266230190308275267188214

90866287610488559337197261049507745623105868480879381472266079656839897298638665064840062114374765263828862047535562161381717148719718046127724426621315677519388283241445469364331324

135961339595981007614168851211391148911785868711577492109560100854595241146710036766240129671092152262676284129312110614451505934544353

135961339595981007614168851211391148911785868711577492109560100854595241146710036766240129671092152262676284129312110614451505934544353

135961339595981007614168851211391148911785868711577492109560100854595241146710036766240129671092152262676284129312110614451505934544353

761617865821869859575596892452659702922654679772592603635611635364107086335561353932052714069353169926762849641361259167020342024115933618521715919513990486469321260477334327324252

3583623623224284053222844142192803184192643003132552752862863161614453661622832601642645529822534212529625521631325031710019410579127819656956638220204149129231134135160116

3521563342612534633234425632372555332321171660372533363030933324120362460523132911131019108128101162519121912141911

79649384878652581034568811095968926469796663221001295162433962156650723676463863596930492818371815925227625138305615304231

262649455040132648213432924440342239242936275554124314133264519531030281530324444311128152112107318351825262227218

2182511641602492182321662001211441631621291551621141341601702009623014674145167821402515412417659154157103168128172579153396338522352272712493746213085647866

906010980881158259916174779589102100776471728941129112418246295323101866341874644787994336126156025221816188656537294454512421

7758947571107744679546666818094886158606080351149838724025482085765532704042727282295123135117191715178555632263946432221

132155178813127811149812166111296151431064531610891762671241032983111109535882

313195394419353339171253387172305307408301277359260264278253230162496385152248233127210622942202941012451951532212622597016511065149799985845544201200135102202146141140115

2481613163282992641281883061432432443002232362982152212272081841233963071292141919316656235177248801951471081782022166014391531115870756747271531651138715512112812091

653478915475436581296263108784161454351454639100782334423444659434621504845436043102219123821291017817483522154725132024

6131905894856755854050811236086784783515148301376439753839441177446734725919315493274026152923241220129465236364533232820

6131905894856755854050811236086784783515148301376439753839441177446734725919315493274026152923241220129465236364533232820

6131905894856755854050811236086784783515148301376439753839441177446734725919315493274026152923241220129465236364533232820

6131905894856755854050811236086784783515148301376439753839441177446734725919315493274026152923241220129465236364533232820

171741089126333184251669117886948917362230378491154461371619665162901748715716145751133714247139271228870482108315539875319034141865609118483074154652401017116653714469858912249126101480121775553086783612470815073106065544402450083079143299167954699260897531141741134064405179

5763479895615706513144657613295724797134866144985293914845145172898084802786774182054388652793455124044027834539641784923231511617737328118312617312447380272271262277364278180169

316502020201081612171324162121117151120749168321881131943131312961518361718491555276520131318914654

316502020201081612171324162121117151120749168321881131943131312961518361718491555276520131318914654

5453419395415506313044577453175554666894705934775183844695034972827594642706454001974278350889153822742826933938139981321529711216835827617812416611842360259258244268350272175165

12280227117141142751001601041121091451061208717380951001117217611258147884690191352201225282646788861834862233679534026351910736768576573713220

26615943126925830014522135413128422434622830324920419322626121413034621011730220494210382143602521072101211691801653919714349741531268662835820151112107931241681298090

15710228115515118984136231821591331981361701411411111481421728023714295196108571272615931116468136841031131482397092405812697523648411213680839479109726355

21351482300923052014220312591669241810081942180522651604194818691639135618811611159510422657182911631778146087915766681550219020669821730111013171510140222877461100455550105866171652461736017714299849189771092909687863717

21351482300923052014220312591669241810081942180522651604194818691639135618811611159510422657182911631778146087915766681550219020669821730111013171510140222877461100455550105866171652461736017714299849189771092909687863717

4521604747461251562041485336414428305925421358432448211140743624315361132282956303714921131314226329212328291720197

814713667116902768108327074100638071814055516926120834597702339126414974437338584473152355124225545321818136813836433357302712

12268451593137010611251817905130156610619851160814999101089982510178888626211532965699918820552916499862106610936619336897038037761139420591250308579375400291360217100823577511637677462317538452

1399214293581092642922581839311313455503212678107120107772868532495145297125326357978475113234715358221433419159542354265163644267

3819955847912950693358418244707434515161521387533161382339733905926483138352677206016926178192010555272631301992010

41435367250049743127238359923943438854237444843540225335235933517654039725039834317131095360464437145357205260374296499147200901211831401531131166941256191178131202205173156131

192105311170188185761701939319718623516017618014512522114912873213211861881157513734136262235602207912914212725171114464611250673562301313110790817086746138

112026815165411216510565115716100125881560856161005090001318811461113781071596567325926991507888448213901103595644128979659344375461811108391257711311412598345615855684101064814994300656502378318011835834935482632328020648686239532945373764470811055880241713266

45122966631753974273493728094554603423524339373951554193444741934114314937753862315419755328433123774772386315323274715381548994279158137832429311035623684512112162087922120733172337139311441310808372256020141742142220663153272716631284

22010233627423126412932239511219816730127127319819214421120019263324209121363222631603429742025191216129164193284399811145276395340494559372213512210010610638533210161

7620156435887238074346247696159723137634143198250309564173677815978347127863268128253952676551461614449224135196272267

92559413251153841840575950125747288574810448739608838476747338336204161049895495957851282668137756973902315761489645750818102325942619324273751532726124815671529417320250388710594342241

2674189235433191257931271721251035471427251722503130241224932478246319002238218618281192327826111440265322319902008445212926222368910219314831713207020522860684121654469815981105801671771496227154311671041838125715131349992811

6173589577365646193616927613076775276115766625625813945306024712855955662917044951804029255572568023154230150251746271116729212816551132220216121610548304286240193296483380202164

2558165437953004252526741570386647261367233820873226353733072465243318042125219018951090364324261379387429358531754532349736622411936215714061677195237354063782125253671256253794817615833496198140811481030822117650753812935903

3732135464063313701964946331793452564114644193503422462792732991124263052065143991262078144646534416330022319622845463710215473102679467106791107428258129156124195603512130106

51736673058652551233786510122834224286957807234784253294103773732435814852738386521643171027897734651704182833223638778341452561081151434100814312217476462582282101762581237947194168

182116376266238268125266349108190189410246237243221140185186166999412161113292256916347293309201862011251511933023348911446684302647148773411135968165983482488369

1051162151481391921163063239115413024029230918918313012212493511541578932219050129282292531404413782110952822776887212846631246347338591706752484163095734

1075111897668450162149387265721411178381568072442795644715210430621511516971237445616612614024421719197134341613139223735243417913832117

149872271261141586022525068120125160169177138119891131171025215712761194142301153422718514543114751091321882124464344527919653433824674646248712611834951

81942112721013796795480121215024277366779321126101473479060469679061540288177843111899692805641571073115479830468638656366811451282234413180267171311722862062631686742938233526034816111161261259

306284311362316295206336508173299217306319311250272210240251203104408294161336254104197683253542471032271871652073613477612257684272417867856926141142847312442031412999

17377292358105212831424584201118396801131129912061339118712078628341459113010704451380107460613781121332921177125715771883564156767820881098128025253447462585361193939413230372224808955326463824761071955443355

776015096101121421091335895861061149710199641191038835147696011110322761810513891591005197641161433043283310583492236226533764313798894036

23317436622919726012934141810320217724726927018421713017618017310725919989297279691923529333522810320798166152314419649755834313196654674120146103106721133893357569

2522213372722352581403684731392001862532842882161811721952051478725019712337034183154413173812296419913518216332241260126597048837582637135201318698741104384129770

1175274150545575078527311938153806348506006725327063654689696426622167246093346003981584998354272313353381061394164371952815511904801163501691622169119812634565306378205216146119231180

1244854244614801411151561111441642664124797824611354139618681108814947920101048823621479692176680638181819613701326154261913955967999531084189935710924613998847395553334293371237581166731785567961666693353

18412129718416916972181181761791602572062621991471201601441478421618592240126769839169279215721781081511341642935810845541731497242683917971049380591741738549

2131866023842682461432563491381972213702702372972221712242011761013273351533011705715324242273295135244131167211208353791787468228153994686552315818912212613218314213055

265128434248261311127275371114209185371296249327225212162173190924412621493471808220035345278278119254107182172230368691615892259201955986602216717513811812127119711564

58241911136647137892694327413366624121463582648104551431140140249721113786972988783301663679861449675429371925029943648288515164528418522423628918618918361336698378461255333154363185

11516121625123210731021526101313675539958971140103810419821139724963104875948411881049590110793434577919190011131196425932506882845865138630747320132681654037031033619995618469388353423795642437371

858479129710468067474347381069415813710934841799763947562763837594387904829476830738282609146636793931339727385694672656105322535416425659239128925326515866470380298259328601447347305

29313332818626727492275298138182187206197242219192162200211165972842201142771966317045264320265862051211881732093338211937702241498157714129148899094951941959066

760638115681868481144661099233084059592364767664867955865864551234290869138076059124258112358983565225563535751258859384023545819022043028625517727117178440396355284303421373350321

760638115681868481144661099233084059592364767664867955865864551234290869138076059124258112358983565225563535751258859384023545819022043028625517727117178440396355284303421373350321

23418343728925025912317334911131818330624121721925621219320915013126723713129521774216432072861928321810213317320128466155516814710098568239301351251168790157112111101

5264557195294345523234376432195224126174064594294233464654363622116414542494653741683658038254946017241725537941539255616930313915228318615712118913248305271239197213264261239220

269220547337336314146299372208256229371300288249342237236290302119306282132360272104210492535663101252701541932382343581151396167187131103639350321841291289615620015010989

269220547337336314146299372208256229371300288249342237236290302119306282132360272104210492535663101252701541932382343581151396167187131103639350321841291289615620015010989

12212230719515817079178176129132961751391351172111181101381765115513570178130431142611933316449122661011011161565775323287755335532813926263468390844643

1479824014217814467121196791241331961611531321311191261521266815114762182142619623134233146761488892137118202586429351005650284022199267655073110666346

16039983098161119161677880125521657131302114515471253190212671221980123211681026561170413338491900122748610322481241574016475811082729919105810681718922711290366834659539354410232164873604573472724845765551445

16039983098161119161677880125521657131302114515471253190212671221980123211681026561170413338491900122748610322481241574016475811082729919105810681718922711290366834659539354410232164873604573472724845765551445

205993361472452391141632438812414019816225516516512816214012560257172105255163611392715668621364123991251171502251049232581311027349512216127836660701181157844

35322765743241649617428647015936729738533837232328624130531228012940737223940633611327266285810393113288175222275280434130193818919914213191945930232158159132160219197150107

53732070952644150135542152523537534242538444137741230440137430817450339120141639717030686354729413165330234281298283479162192901032421731321171287244228167155142229219183173155

508352139650681444123738592723143636653936983440235830736434231319853739830482333114231569446351562823934122129136835558052623487116262242203971377974286196193138265289270150139

3082554653153203441862754481692562654443243162842952752793352541234483341653242671512605025036032613829520522924726039710616473781941108777944122216152116123144191151138105

3082554653153203441862754481692562654443243162842952752793352541234483341653242671512605025036032613829520522924726039710616473781941108777944122216152116123144191151138105

3082554653153203441862754481692562654443243162842952752793352541234483341653242671512605025036032613829520522924726039710616473781941108777944122216152116123144191151138105

3211365283132863151582012731182281982142153651862142152082141949035123114233829299205392168082539118314117816317933216814149701621291137170374415588941111271891347867

3211365283132863151582012731182281982142153651862142152082141949035123114233829299205392168082539118314117816317933216814149701621291137170374415588941111271891347867

296133479298258289145181256113210185195210352176203206201203165863202141293082799319338191759232801651311671581692861441204360149118110666435421268484981201821297065

253491528261320175181319513101197112943117133013612125492111181011510462421610131135622294101377582

14618781898222613711383872102219948171465122018941419135513801494109112531582110075520281578779116611534881076245118992613054801325883591110910641354327766377351572348484373498304137836770543476874526445607564

14618781898222613711383872102219948171465122018941419135513801494109112531582110075520281578779116611534881076245118992613054801325883591110910641354327766377351572348484373498304137836770543476874526445607564

14618781898222613711383872102219948171465122018941419135513801494109112531582110075520281578779116611534881076245118992613054801325883591110910641354327766377351572348484373498304137836770543476874526445607564

14618781898222613711383872102219948171465122018941419135513801494109112531582110075520281578779116611534881076245118992613054801325883591110910641354327766377351572348484373498304137836770543476874526445607564

744399773108163558745245695636770258291676066972164452058778357233491881738455052421046912056843262022363645829752450565315834019716425816822417422814971430353258222432203232285291

3792356266743914322562905882484543695593814123645553404104742922216524492363723851463446934226533213039223116132033739398240981101761041591191578930222234163141256193104191161

3382444994713453641642764502023092694192782742952952312563252362004583121592442441322635627922935312729719413326522230871186827713876101801136636184183122113186130109131112

1704153715641362146615478041106172011331127117620311148141917181197121410811302179574823032135112613231134612166235394290312684931401124972311118061243602114365448213311044690820119353034813951441116176713141650143112441012

1704153715641362146615478041106172011331127117620311148141917181197121410811302179574823032135112613231134612166235394290312684931401124972311118061243602114365448213311044690820119353034813951441116176713141650143112441012

1704153715641362146615478041106172011331127117620311148141917181197121410811302179574823032135112613231134612166235394290312684931401124972311118061243602114365448213311044690820119353034813951441116176713141650143112441012

1704153715641362146615478041106172011331127117620311148141917181197121410811302179574823032135112613231134612166235394290312684931401124972311118061243602114365448213311044690820119353034813951441116176713141650143112441012

53549060870353359833444573932647643065648449951755438849747939725981864525740037619539194398385405146478311203418326464832551221272281391531361369853274328196178336234198224188

1053962799570775842389538821726555622118757782710845807344867441321439122413527858406993731191249415468772295805883492621400626492793473331106387250467210324142871042104092952591013891208975794

1168515789158107811231608196124188879311763929879775026113884835944801012950915211855287280153279559244033331225188797336646827254530

5568511356326016490157993908506660732755405043515846446948334972488942024619439933172213669246462553383541062472421010153749323440621583355633563190429434254139111121761241132220491261160012741409923510232019901565161630022382219820472002

5568511356326016490157993908506660732755405043515846446948334972488942024619439933172213669246462553383541062472421010153749323440621583355633563190429434254139111121761241132220491261160012741409923510232019901565161630022382219820472002

5568511356326016490157993908506660732755405043515846446948334972488942024619439933172213669246462553383541062472421010153749323440621583355633563190429434254139111121761241132220491261160012741409923510232019901565161630022382219820472002

5568511356326016490157993908506660732755405043515846446948334972488942024619439933172213669246462553383541062472421010153749323440621583355633563190429434254139111121761241132220491261160012741409923510232019901565161630022382219820472002

2211642282462342501382112631251762172471691761771451601341461489830721790148151121138501701701765817511786143137168611026651614160445235189691546812370607664

26062282271827902356273618032482279412851812198926801986221722322359187422211998150597429402048120817891867111619844741741149919237531554152215301861153618454809775826029445877355556654272431083901737761135710731052911993

73776474566255761043059764933750760379961062375646849263266044235997454430843863333851410850936349619448842133742942757213925314820626517620519217414165278239216226370278250270234

78870788210768028286466421106405691671982689757797662650691683590303136582739365652930258112957754065226060848136167653170317539521118226620426621721812774398392255258446310333324265

1216119610591242952137589111341261603864871113810151060101012551026941912632479110610105548049265959932547526628153187318158761185794851256449234281513253334266300193110465367303303706651503466446

9053516342515942542809722301024235612750870967277130883978399312268111644812615513355626395018678914213991465172091558491092731011923403001127141423451593236960215918768424684817820512447377377120272110140745531684722076938209125927647443644736948491343398126050929774795729179174518712156205395573890458263713951812957821897728

9053516342515942542809722301024235612750870967277130883978399312268111644812615513355626395018678914213991465172091558491092731011923403001127141423451593236960215918768424684817820512447377377120272110140745531684722076938209125927647443644736948491343398126050929774795729179174518712156205395573890458263713951812957821897728

2009517175231692801625769229172609216212428271674617024243392609019086204402757932097228391599019144141171071526790274613362318487173171823418252135411813215020156514562115356437481015128401230401811645621047553725257140081597462986886608640552333108911079916807855517187141061681989159212

31230832543440835434026662520327325538927030643635927525535120619738348129031430314927312828224120528823855815243235030161154968626627977121875827187118226102214271326135131

31230832543440835434026662520327325538927030643635927525535120619738348129031430314927312828224120528823855815243235030161154968626627977121875827187118226102214271326135131

137157171204189213170145296112141152220150166221193146126161123702251931661611201321537215210012914912825085181155158399340381381385448333098997114701161171447970

289253053302532561826354928326831281623261151223235292227642251722283016364240720125565210957320152418173737159

109148146174136183145113240941151171711221341531621181101389759174171134126911101266611075112127100220691451131183273283382864439282366982905299801076461

9110311314411710879791478890132155931181051401009111669711441201251041107312043739068107861136512511784155125187856293034135535748355479834134

9110311314411710879791478890132155931181051401009111669711441201251041107312043739068107861136512511784155125187856293034135535748355479834134

19714121020825025114420132312716415925321820122019815816419413585267254139185204106193481971431221211412011322072091734289494118710360764735209392110581011741516773

19714121020825025114420132312716415925321820122019815816419413585267254139185204106193481971431221211412011322072091734289494118710360764735209392110581011741516773

964815104613131014103572675614315308127051143954998109795080674786458649311071236632890805411811221797768782417720710396824915879176446238218786541246260288175103453461330313526801766377379

964815104613131014103572675614315308127051143954998109795080674786458649311071236632890805411811221797768782417720710396824915879176446238218786541246260288175103453461330313526801766377379

53140169666166256241350588136947447467460158160461837843353541226581877636556251523944812657740346930448951122455650955912126413211542336717815815411071295332241188376466481254210

2822254144034303372672935502202702654143793473703992362543212291484764972343463241472617638223827220928932413436232731561156756327722710492906640175204157111226285330144121

249176282258232225146212331149204209260222234234219142179214183117342279131216191921875019516519795200187901941822446010857521461407466644431120128847715018115111089

1631142072451852101471502751131641672171771831932071741541731179029323012016419868191511661631399611815377174184155277240431291306757423517100699562891631607071

1631142072451852101471502751131641672171771831932071741541731179029323012016419868191511661631399611815377174184155277240431291306757423517100699562891631607071

28531622356035784305316191541789128801808207023444749237123906967752624072233219818471236472140194305225418451450307234192668183322579925203222714106874492157224655617118607119344759798803843532317145818055918106354371103178611301387

6373551068767156470762638297442051852475648453676653153043745544320511158361592523312485454200775335550136063486919384449459512833418011317019216712015211653372422443209483195244223198

1027463100512261135102560247314335095546049736987978551004618576698509454134513961260623620409598279899734478873513852348877624629177664313188366266296309253196114406511405320460412527371619

1189804148715851606142979269341047387999812163020118910575346599112591220104589557722611787145311089135562020294099476412297692885209935275728103910222517133674103984301335374438220150680872507053444944961015536570

244234323342342298289199413158163184287203194321291227160246107112326377231191221137161109175149163228169370712462221994999494713015559856360221141031299020315520184129

244234323342342298289199413158163184287203194321291227160246107112326377231191221137161109175149163228169370712462221994999494713015559856360221141031299020315520184129

1661001791771611661311182741161511582281411601601781361371359581249191921631129213638139131146721321167715315816947753853101744458392814889673481281121017460

1661001791771611661311182741161511582281411601601781361371359581249191921631129213638139131146721321167715315816947753853101744458392814889673481281121017460

4338495268417030903048527440495557376061463083653159421854293642563235594263555112351574030221526125252826147736382419

4338495268417030903048527440495557376061463083653159421854293642563235594263555112351574030221526125252826147736382419

2211963085223282712731554171891972062642052313252792201802161991312923822322142091342581182071281702611842669920921115659148465019812574687549281451301219714713821698107

2211963085223282712731554171891972062642052313252792201802161991312923822322142091342581182071281702611842669920921115659148465019812574687549281451301219714713821698107

463969494660463968345136554247513438444531387055315125284519523649293930244725398309143221101611157232418123032232620

463969494660463968345136554247513438444531387055315125284519523649293930244725398309143221101611157232418123032232620

336339405399368442268251509196347271391294325309255225257289212147475395180269249120238672652282751492422931343012623046116585591961788595736737205167139116210203216124111

336339405399368442268251509196347271391294325309255225257289212147475395180269249120238672652282751492422931343012623046116585591961788595736737205167139116210203216124111

20521024727823621216120236915520017029320820620925018818221312810328123213219318976153481951921611121642138721920820941113515019516259685753211331151048114121817889105

20521024727823621216120236915520017029320820620925018818221312810328123213219318976153481951921611121642138721920820941113515019516259685753211331151048114121817889105

84781298811969949857137841401538740813113081797497093865678989560343210921148577840777414819240785718646456673831432101689783220842122024562748423227623016482487428355280518633752348355

154142180173179163128172221113149165206195166140161106109147101851961808716812076133461531401046012114376176191149337733501207545424426241007652531021171485959

38142545658540647730931763822330033650333346047442128737937825520142046426035137017939811933330327222231137918846439636297172941192732211051391038736231183140116230282362170184

1156612116614011498942048611811617211213713813310112013187521701681001129060883011079113739211168109109110276234238363233230196646776436773825342

19717923127226923117820133811617319624917721121822316218123916094306336130209197992004518919615710114919810026720121151110595315112559635332169210287681191611606670

1371116817162003165316491523139624698291283123417741460158617451711120711721364100980019352204122014881383689114350114081091124498910801231677137416101412444789506416137099858560456228620880872371860992113331337728631

1371116817162003165316491523139624698291283123417741460158617451711120711721364100980019352204122014881383689114350114081091124498910801231677137416101412444789506416137099858560456228620880872371860992113331337728631

2793043563893183143011834751832202513262432333343342491953092221493064342582252091472631261801842042312294221233442781985413971541211148810567692612718913672182150238116153

2793043563893183143011834751832202513262432333343342491953092221493064342582252091472631261801842042312294221233442781985413971541211148810567692612718913672182150238116153

2133187022953001379124213294150938385387179490602318181822962824681274151594197914221538251329351708618261708920021404981170312641588250861590479617614329583817145361063494730932165872189562039922111691113317177620021146185311671102

2133187022953001379124213294150938385387179490602318181822962824681274151594197914221538251329351708618261708920021404981170312641588250861590479617614329583817145361063494730932165872189562039922111691113317177620021146185311671102

201210225258213269165174281120189189308181205204193151161193165117240269149179161831745216415616411015518612918915516535963543131104556136452012411691601271131728282

201210225258213269165174281120189189308181205204193151161193165117240269149179161831745216415616411015518612918915516535963543131104556136452012411691601271131728282

265197336486319376237270554193263261356301325342389264219281206135415422194304300116259862752532481622272861512783132597615078652842018610887532915718011397192276261150118

265197336486319376237270554193263261356301325342389264219281206135415422194304300116259862752532481622272861512783132597615078652842018610887532915718011397192276261150118

19381594226523662038193913121741294311191565160525551787195521311855152814211725126087223862290131218251827849151155018681486136098615251574732162518101959409909451421151011725554535603362051013984785693116015401565727699

19381594226523662038193913121741294311191565160525551787195521311855152814211725126087223862290131218251827849151155018681486136098615251574732162518101959409909451421151011725554535603362051013984785693116015401565727699

47242659477062259542844185632450847170151357460062240938951829827069074137348844423139815247939242630743047922256151349010124914012935827314817213511042261238261194304379424229216

16215017918721418613017727698216165220158167206178136142157103792452351191621058613342168132143811371286915617117238843445119776455553415937067651011161228778

310276415583408409298264580226292306481355407394444273247361195191445506254326339145265110311260283226293351153405342318631651068423919684117807627168168194129203263302142138

705663694111083181265056811284805616218686186798728726855776704474298031037773659583390698332592569525712547781319798630531299489261259425462385383332208155347383501351465430647402334

705663694111083181265056811284805616218686186798728726855776704474298031037773659583390698332592569525712547781319798630531299489261259425462385383332208155347383501351465430647402334

11312512716313121383682477589971491031211421251101281031026412620513911511391944212711212213310297551481172011463503188502938443114814769506185923246

11312512716313121383682477589971491031211421251101281031026412620513911511391944212711212213310297551481172011463503188502938443114814769506185923246

2732763443633293093562356011672032253302222293674061841942261571214243132932181731402151172551901913501905991243551832514515476611521797980574629142142203105189172203115129

2732763443633293093562356011672032253302222293674061841942261571214243132932181731402151172551901913501905991243551832514515476611521797980574629142142203105189172203115129

582514663836862697521580126536149447369660067268366750342652240826697476948162855824038317364357345239849970420354469272212728217314260966914916419612343330361333223354610715252221

1821892012292492181522353461211551451921852091672161771331441429029927615016816264107542161681319316418778158174229419448441821535955663911104110106691192121758057

31162932473639246825232729182838251910142193533244723152363530201626127254337711774253317211698111543491412

3693094335755664433303218512153163014753974354784263072833642451676404603074133731612531133923753012893095051183614754567917711891385463871081238231210242219143220355491158152

3825385278055876225253661020300429349581522488609624457383581286301588729385483461249436189449366365358368683250648537439125286158177843414183233150115129338425309241406604588327382

276398341553356443316224616205288221412335313373408312265404185223384489241310279175254117263233230204227389183463316290852031161436212701411741037999246305205169288406397219299

1061401862522311792091424049514112816918717523621614511817710178204240144173182741827218613313515414129467185221149408342342221444259473630921201047211819819110883

50257438852346652842935769925438131050939140348055031838642629024740652532334632522237715030233227927929759318648535933384203788521919012313211712147202154198113284191303154197

50257438852346652842935769925438131050939140348055031838642629024740652532334632522237715030233227927929759318648535933384203788521919012313211712147202154198113284191303154197

530539439589573478480421796326375315477447462499674418399522344256384523303359421259425154303369278286281556352476441332882001041362441671371851329338223166207109294258328183216

530539439589573478480421796326375315477447462499674418399522344256384523303359421259425154303369278286281556352476441332882001041362441671371851329338223166207109294258328183216

15541244153724431611162113791033267890312371230172915461564174019051398121116221599821173520171233131313177551349612124111001178105210421772945241014941272278715366398101476645055447131618986676583016589619591181641786

3371532754382372742211483971611802022663412952333323012422405361673242631812142481572211191931842091171782292636702481866914368921929910210786764315411014536717317015488150

767716876142896393574657715014856906879617788501019951715630883686455944123868973570238374230873461062959559810194241106850738126386196209592450228302238171103467484406543495551713372433

1461251001591111131151132857913011819617713612622014212418310860142169118120110491016110194127108791566615515512121594532877034426717137367701068276896469

18412017224318315417311029110514913318015517820226113710919718178193191157130145101196861271291161559722411531613811438783744678156634930188643140562126861147270

120130114175117145124852047388901269510516014110310611988611321568811411265893886839777901447716310311324492021766630403122128661698085761114564

17220826934420921517917132611318116226119215522722014113118812010522422115115916273149451891451341241371791112041581653810542291291095266454010989797621011261588081

17220826934420921517917132611318116226119215522722014113118812010522422115115916273149451891451341241371791112041581653810542291291095266454010989797621011261588081

20718023927224319917820937214718717523021322622726014915121212895268264173231229102205562052301771661782451121982382364484414026023457785932219493128841192562978783

101741221481511011101152238411589133117124136111818412777431411501051351405011032127122871021021664710112012722482319135138283933161443467642711381674535

10610611712492986894149637286979610291149686785515212711468968952952478108906476796597118109223618211259629392616751475242481181304248

24723433036229426424517742713522822238222320630230121821322315913031626720920319916424812419716822625417338012531121121748101454916411175786241231251111718318212616676104

24723433036229426424517742713522822238222320630230121821322315913031626720920319916424812419716822625417338012531121121748101454916411175786241231251111718318212616676104

81368292911311006102768364414535647528111018922932100811048646287885794181231114292082482058766032378667565389271679740992282970619543221019769550124726325215599438390439346516651669347371

2382012513223142862051814621991932623212722522763652491662061671263763553502072102211741162171851854462052649731522419950116725018212254817442301191411419914519217710498

57548167880969274147846399136555954969765068073273961546258241229285578757061761036648620756949046844651153331260760550714531613814751337919318217811369319249298247371459492243273

8995127186161120101932095182961481041191681038786128636011519279133964897511136810375821173410614811729602723155103335437191270806449771171395674

8995127186161120101932095182961481041191681038786128636011519279133964897511136810375821173410614811729602723155103335437191270806449771171395674

8995127186161120101932095182961481041191681038786128636011519279133964897511136810375821173410614811729602723155103335437191270806449771171395674

4789854216709041440828980013687859606419561295593759249911561086982477995933518167610836435025563497242228189323575628814757646410519054740220476448511772237422503229261524693568915256047759889268813845320114112314026221266072626513546140333019224965139116822289483666716076239363108144245530423780738979

4789854216709041440828980013687859606419561295593759249911561086982477995933518167610836435025563497242228189323575628814757646410519054740220476448511772237422503229261524693568915256047759889268813845320114112314026221266072626513546140333019224965139116822289483666716076239363108144245530423780738979

3147247734425629362147262512182154942151257227043678327337313726394321312675361017991526326056581891268023451100243774725832444382810972852236715893619321626926261195909857113862880910139816823131427167192197615761487176715011407

947580249049125069960138486872563512983592482847535115519588117061084910647629177819436561741809485148064906777253222993640318616696685911849269579856357465210326826267221583368724902508340517542332262825521775759416046262816270245993901442938203575

551579021315933908189323046411673623730551483166787226851613557139601452025686494577971374438226164663433712993991979449372176863767368762859835332078775918556010867152164712201436706238319236941555206771274590209910594843414823304196437869811103761959536

6997845119042463602707743784172599642427136889964010063125841992419877211103562472531039119178603081881004449820144751279114081568497935538525387641177339751015470796725135972006361552936538572963893455521972594873858252467139771005598350155276641934215164788411725

11023898512576165781299816731837566721733371659399885413524111881270512069133497213883110327694750621284818851617692326960357784512384791176391120435308984751055911037910145873319984770305827893752211828643350319821811017536056073372337558574561499348544606

11741183771363629101172122732512915119492048510632133381972619971204653137219402191157192188741612739747237140172472990231023392453401100813425112921833144126100761903923329236424013831236163062254443362301336897215294336773367819470722776058150173136716080301797315652135538130

6447791067579519712656567078816083594767725442126903590822256790537932634227521048714421620132582324222053749272745132853933

6447791067579519712656567078816083594767725442126903590822256790537932634227521048714421620132582324222053749272745132853933

6447791067579519712656567078816083594767725442126903590822256790537932634227521048714421620132582324222053749272745132853933

2572074003532712902062264121612622564212933113193242282192812071234663941842972571232246129820024610523718011424927724375164666022614999747955221491741139517721915010395

2572074003532712902062264121612622564212933113193242282192812071234663941842972571232246129820024610523718011424927724375164666022614999747955221491741139517721915010395

2572074003532712902062264121612622564212933113193242282192812071234663941842972571232246129820024610523718011424927724375164666022614999747955221491741139517721915010395

214709104763950254426106141811504093511932102965333931600741188189152789918524153422450258336918165793141268765632164721635403197870845936363045895101369051191612046151106634470971301316150500689543817578968230482453948318005398453085551616568109978195466678017621160129307122361106993097849047

214709104763950254426106141811504093511932102965333931600741188189152789918524153422450258336918165793141268765632164721635403197870845936363045895101369051191612046151106634470971301316150500689543817578968230482453948318005398453085551616568109978195466678017621160129307122361106993097849047

37352545012778176453192478552501873213033763099916396013041370188369436103237357062526676026726125820488322189162424617245752447768887051317141984931215181105142126573241353

1699430934031107053394239213986551151850127423191781179515477431254741011341396910453053156175691666351843081503443316961017848118651225119151003254746681338526158631934853829344324858746224172293737511460746795484316302279

1907285531594642402819898175474381331070395976309981434438128165932535916418143717409937792916313726301132361649142031535672962366063878982844789139346541045710816135635945438341178214857466328938815795899328371409043687308356547455224415378101657655436837004554253778293111931043972881145995

32668263317981004792344117101929520242023836925012838467382071345131561272137231669480524276928818112415789430127128465765200882815347715669470175112743791257770126122886296420

6626387969287939225174511032398651624972697751703817544589730532344935104740558853324160613751049666921754648031867054863015438120615034316725623120012262359419259230389340361320288

6626387969287939225174511032398651624972697751703817544589730532344935104740558853324160613751049666921754648031867054863015438120615034316725623120012262359419259230389340361320288

6626387969287939225174511032398651624972697751703817544589730532344935104740558853324160613751049666921754648031867054863015438120615034316725623120012262359419259230389340361320288

44924479480639615162438927894471628220783180308648273738364233233209279330573215256715924899316917203442320316752836666350731113245123129522467212232693725441683616618617913077179410421004970640342176915941259122922323156236113581487

261120602666202028052416148220803075113017321745250718561941183217781479162716961381862270217389811732164793915223771781156118136561642130311891665178822754558925144391321763563553557381192965875725685116413691131693803

261120602666202028052416148220803075113017321745250718561941183217781479162716961381862270217389811732164793915223771781156118136561642130311891665178822754558925144391321763563553557381192965875725685116413691131693803

1236522813428012777199336751021192271591541371209285797821328123841971176765242641329158130873611520532736824133279169292636226591115572992821825818

1236522813428012777199336751021192271591541371209285797821328123841971176765242641329158130873611520532736824133279169292636226591115572992821825818

163103218153260128102173254991261502421381171141191231191051253426211983139111799024175184141511467769126142208316539271119533303723168378477390132816257

163103218153260128102173254991261502421381171141191231191051253426211983139111799024175184141511467769126142208316539271119533303723168378477390132816257

2325189222201733226521611303170824859561504147620381559167015811539126414231512117880721121496814139614197931367329134212451581547136611391084142414411740388745434379931499501497484336170823686623540975955868573728

2233183320591620203120511245149221828951416136518451419153114641474118413511442109476218781367750121413227491295313113111051493491124610731060132412151458359680398371640321465474455312159746609578483913638717536709

92591611132341105821630361881111931401391176580727084452341296418297447216211140885612066241002262822965368291178362329241177774557623171513719

1653219618261683194517361155202926258141245113519011576141312441244112512351349997641177211735991350131264711712491392131012444591082102284814541577171932063127830113007673994063532181376855424374229111358990562614

2462333442433733121582684731412182264072472722272031752032461541073382471273022598816347278179214107216120872373113436219658432661637258624425116124821051332791729079

2462333442433733121582684731412182264072472722272031752032461541073382471273022598816347278179214107216120872373113436219658432661637258624425116124821051332791729079

2462333442433733121582684731412182264072472722272031752032461541073382471273022598816347278179214107216120872373113436219658432661637258624425116124821051332791729079

1407196314821440157214249971761215267310279091494132911411017104195010321103843534143492647210481053559100820211141131103035286690276112171266137625843522025810346043273482911741125694183553177781079818472535

1407196314821440157214249971761215267310279091494132911411017104195010321103843534143492647210481053559100820211141131103035286690276112171266137625843522025810346043273482911741125694183553177781079818472535

12871828129812781253128090115461555576893751118611669998459558629061005733500101475339585893251892218086695587627270783773011281064105822034616722783148828532625215396482302295219695906718422511

1201351841623191449621559797134158308163142172868812698110344201737719012141862224817615480159653189202318388953312031164222392116871166098831731005024

134137151134160117851082076410913216513813614010210112286117591981287413011940752212096108591368451911071583693343111073632334225559664628387725946

134137151134160117851082076410913216513813614010210112286117591981287413011940752212096108591368451911071583693343111073632334225559664628387725946

134137151134160117851082076410913216513813614010210112286117591981287413011940752212096108591368451911071583693343111073632334225559664628387725946

134137151134160117851082076410913216513813614010210112286117591981287413011940752212096108591368451911071583693343111073632334225559664628387725946

948616312425212067254375709474254168152107858873847230227130662301254968182141448057925834592532642545352034619117222619864813360743421684424

948616312425212067254375709474254168152107858873847230227130662301254968182141448057925834592532642545352034619117222619864813360743421684424

948616312425212067254375709474254168152107858873847230227130662301254968182141448057925834592532642545352034619117222619864813360743421684424

948616312425212067254375709474254168152107858873847230227130662301254968182141448057925834592532642545352034619117222619864813360743421684424

880778071096910810876192486032109691361155469600770411767112769783882988386930897191076289458011598100114804105118682349369431708103067749747327536834524441017502100869504215752862353232413191664229372596283320309154578466134262828503312074746040793546

840475031039110271830187535716105991293552249133728411126107409275834383686557844886105970437910885940545221013982913347657516339951745770942612650149823849709297229072205849772201220012930648927852469268619218744353439932502662479211843722938643397

815664107410498107805246379924988076891069686702704687502807681573361114494440263155227556613162749460621753648831967755066618957620421043625024419223016978364406312316419352302328280

815664107410498107805246379924988076891069686702704687502807681573361114494440263155227556613162749460621753648831967755066618957620421043625024419223016978364406312316419352302328280

11766113140991013976142721048113073867973611038667371597946686825701587588228565439778579216929276043292127245425827574944414030

716510792935856104113578487869410084725410891514311685346744356516635556216054409870792554222453383020212010434324324239294240

45042149846441342528132051625445035553435838437340829340433831719357646221933532915832472316269293109288289172362262341992759098204107127951278446189202173132240190156174156

117521301651441187977126591109922189821059254118959446203123628865395717917110436854632687210426100363465302925332311555944534837374240

60602261886178696095565967987250634240747144429019541734618501170417123474536726163187827275432293122186354444424042393014

13941249185616211394142986512661856861157711971792131612731300131110161397134293970118031499701118510835321031285115288511724021069764610114110321227319788366357900445428391452342136656672536428815817630633589

13941249185616211394142986512661856861157711971792131612731300131110161397134293970118031499701118510835321031285115288511724021069764610114110321227319788366357900445428391452342136656672536428815817630633589

48739561264454150634944174132552842857248547051247338248447436526760757724144839522037310140333743613539831820343340043311728812814131616017213717612147257268225153317305229252231

554394117867539518346567081636664695666635227996931624420321451383913402825496466113613214629131320137272824194662232221

7737551063772687782437679952445903636104670167367368953372071446638298777539161359326357516063645263823457338934459951266616840820015447322421721424418971355324257222408399347327306

401324324224102924132421442023142515203517755241421151314424292172211923242481786259753827207141916111712

3943635638423066563266424947413755301075639185554244136163763829381336182937323815391735402319229119103223202535201519

62063488778663068652499711084237415789169647876806605387336844993659417323869007423184981259186295872265604603265968708221924321801791250566284199203163663903672762454291189723357270

62063488778663068652499711084237415789169647876806605387336844993659417323869007423184981259186295872265604603265968708221924321801791250566284199203163663903672762454291189723357270

62063488778663068652499711084237415789169647876806605387336844993659417323869007423184981259186295872265604603265968708221924321801791250566284199203163663903672762454291189723357270

526546926236644852065584360374368563328256704582696375076243529654394253523755753769281166915883286871935713211942211038701752594522169741123094239744067042614612793052137713931020451301746160117201186563279927951995160629969349542824262137

526546926236644852065584360374368563328256704582696375076243529654394253523755753769281166915883286871935713211942211038701752594522169741123094239744067042614612793052137713931020451301746160117201186563279927951995160629969349542824262137

21817624428019825214715924011524119327318820322224517027820914710727723511515019689158481211051495514912810416213218963135647495545967756321112112964412410470104108

293302369311288305192235410202312316403376293277344255348297227172355290171256289127261712262083258723619117623423324077174668014992939298663618615311684171174145158150

251224334331973338322283039462241291219223521157115176003411424866234920173619141868636521231410158178114101534123113221914777041010

17031349178720091496163710981147201194015971368199015631466157515581311145616071164845176216748281285134664412383451140116812885031201851713138411141296426916367428928485572517525353213872803580475891779614732672

47540552159048050331840763727951739565649648147849740647152834422459457625239338817141090362327392184368279241438351422842371161082961641301261339943238249164163244239206218163

21002024278626282204236114794386423613922467188829883911309322622287170922002397153511963051255912394149281190017493834149277619347011766136994718064162319950412686305806898342672465972550020811241214851689128763513488985887

42240246355946846734034759931345236957647442541646335441644829023854349222633431316235585338298391138333238183331344402922631121042371191351121409336238218158121247212189204132

2922423829282022471956233837413733294654411438422026291226819282312231919332033123689209191899217151781313121515

310264338367261274200263416160338238386267270363271248274328190141306347165230201103259542371902077022417619727222821179129746114098838681613114415913167133136146120121

310264338367261274200263416160338238386267270363271248274328190141306347165230201103259542371902077022417619727222821179129746114098838681613114415913167133136146120121

310264338367261274200263416160338238386267270363271248274328190141306347165230201103259542371902077022417619727222821179129746114098838681613114415913167133136146120121

403304578539460495316370676322467420641536508486470373523497319201713606282372391146368753552923791413332622524103644329930915212426115315212714710941225262176166241231231215149

403304578539460495316370676322467420641536508486470373523497319201713606282372391146368753552923791413332622524103644329930915212426115315212714710941225262176166241231231215149

403304578539460495316370676322467420641536508486470373523497319201713606282372391146368753552923791413332622524103644329930915212426115315212714710941225262176166241231231215149

403304578539460495316370676322467420641536508486470373523497319201713606282372391146368753552923791413332622524103644329930915212426115315212714710941225262176166241231231215149

16221237193419782065179810241871263292113861262220415701765153815231195119412441087582324516119641674158663312112411866123514246961307980617120616561972347743443326165097343134543925414872896754067989416441096650474

16221237193419782065179810241871263292113861262220415701765153815231195119412441087582324516119641674158663312112411866123514246961307980617120616561972347743443326165097343134543925414872896754067989416441096650474

16221237193419782065179810241871263292113861262220415701765153815231195119412441087582324516119641674158663312112411866123514246961307980617120616561972347743443326165097343134543925414872896754067989416441096650474

16221237193419782065179810241871263292113861262220415701765153815231195119412441087582324516119641674158663312112411866123514246961307980617120616561972347743443326165097343134543925414872896754067989416441096650474

28622539436736732221733251417529024544828527929129521323722519811558534316030024812222138335247275150261171111231304347681377046320170806483491915217510510817725417611285

2021722112872772291253082901161781312612292321721581421341451327345017813624518578133442361751641021681316817322820740101533827914945426424229012354871142541687359

85461091031321468098179506761130801207892544959531718284651021362964811972824987601865102146284131211128217141611173858315752151915017

38322851944652842124243566624436034261939749542737532132232626314610014232614353991632965655731037519337124617625442857910523212499445295123941026034189289151199240470303201128

66656670177576168036069898333649148374657963957060346545248944123110275833425926182414979561943152820242037224448359469310623216512249427716613117411056259322199228311515358214185

15961192209523661777189510581137225410851719153621491642173417131953133515611645123490023691993983141813836281311329137712651483585139910187801422126515344251030514444858443633507551374150917984647583892648658774670

15961192209523661777189510581137225410851719153621491642173417131953133515611645123490023691993983141813836281311329137712651483585139910187801422126515344251030514444858443633507551374150917984647583892648658774670

15961192209523661777189510581137225410851719153621491642173417131953133515611645123490023691993983141813836281311329137712651483585139910187801422126515344251030514444858443633507551374150917984647583892648658774670

99567512231480105811226347021394667102791412819231011102911237708659567445471410123358583879035075217985776888535186261745686974095625761430825846324735228332220685542627389344536381330442393

460323610878517549304324674312570438625469511564554405410480369258664596316438435160374994443794021654242932144383764931262991421162311241641321668640291317170161255183183229185

261175215221168280144195325163166184227182219187263153189190158121231286102146161781663013218318861157129120170152169511205871104467462714317951097559111857189107

753615270108644246965174751188884756957815276342019253683929569974610042915834596582266538214637241622269396745524927193627

10584125202154106797516968751051569910699142791001488079140127621001003786168793107451048746107761192880392541204135383713786557417038264846

9457121109111123656213073142112155859110495768586615517413252865546702597678838865042957193265031254120493825146396942315148314028

60151787288671977342443586041869262286871972368483056569668949035395976039858059327855915052049759823453740132455352557816841620618639519628122422916865375357258239356267328332277

41839461763050655230029658429045143462650051848760841448451135625265954728640845321237310336438041017336628823439037340511730914713028615421718016212352271261190157244186253234198

1831232552562132211241392761282411882422192051972221512121781341013002131121721406618647156117188611711139016315217351107595610942644467451310496688211281759879

9087061796130016751150645808183358910288201967774896121976570880371374828940179906051339594368690154112768710424551413557288655760155322062935321036129228920730419867692775492877541402288489234

9087061796130016751150645808183358910288201967774896121976570880371374828940179906051339594368690154112768710424551413557288655760155322062935321036129228920730419867692775492877541402288489234

773628152511151396995519667156150386868617216207551072663613709600610253365681851711655012935851379475978923801246462241561641132419254327717629224525818327016254582679407770459326237431191

773628152511151396995519667156150386868617216207551072663613709600610253365681851711655012935851379475978923801246462241561641132419254327717629224525818327016254582679407770459326237431191

773628152511151396995519667156150386868617216207551072663613709600610253365681851711655012935851379475978923801246462241561641132419254327717629224525818327016254582679407770459326237431191

1357827118527915512614127286160134246154141147102959411313836361172881749375105171809015075167954794119229288676346947312434361311096851078276515843

1357827118527915512614127286160134246154141147102959411313836361172881749375105171809015075167954794119229288676346947312434361311096851078276515843

1357827118527915512614127286160134246154141147102959411313836361172881749375105171809015075167954794119229288676346947312434361311096851078276515843

5444157337165876583563857973785955417835635975325514525146093802748416543144835452144879648841553123851236833249442350911430112814827716816517617012756297358241180314247245256217

5444157337165876583563857973785955417835635975325514525146093802748416543144835452144879648841553123851236833249442350911430112814827716816517617012756297358241180314247245256217

5444157337165876583563857973785955417835635975325514525146093802748416543144835452144879648841553123851236833249442350911430112814827716816517617012756297358241180314247245256217

5444157337165876583563857973785955417835635975325514525146093802748416543144835452144879648841553123851236833249442350911430112814827716816517617012756297358241180314247245256217

3382844284453874212162474892533773404893483773413592873344052451825744172192903401243136429823734314130124723333427328062185839416692112120112843517123914991199131162169134

2041303052681982351401363071222162002892122191861911631762021349126523495193205901733218817018696210116991601492294911645541107552565743201261199289110114818581

21322213215315124211231282115131111152222
